# Automated chromatin profiling with spa-ChIP-seq uncovers the impacts of condition variations

**DOI:** 10.1101/2025.08.14.670415

**Authors:** Yuwei Cao, Lauren Patel, Lauren Alcoser, Eric Mendenhall, Christopher Benner, Sven Heinz, Alon Goren

## Abstract

Chromatin immunoprecipitation followed by sequencing (ChIP-seq) is widely used to study the genomic localization of DNA-associated proteins. However, conventional protocols include multiple manual steps that can introduce inconsistency and limit scalability, thereby restricting the inclusion of appropriate replicates and controls. Although the introduction of liquid handling platforms has improved reproducibility, most existing efforts have automated only a subset of the workflow, and extending automation to efficiently map non-histone proteins, such as chromatin regulators, remains challenging. Here, we present a fully automated implementation of our previously developed single-pot ChIP-seq protocol (Texari et al. 2021), named spa-ChIP-seq, which enables scalable processing of 8 to 96 ChIP-seq samples from crosslinked cells to sequencing-ready library in approximately three days with an estimated cost of $70 per sample. Benchmarking spa-ChIP-seq against manual ChIP-seq performed in parallel demonstrates comparable signal-to-noise ratio between the two workflows. Using spa-ChIP-seq, we systematically evaluate multiple parameters including shearing and crosslinking conditions, buffer compositions, and the ratio of antibody to cell-number. We find, for the first time to our knowledge, that weaker genomic localization signals are sensitive to changing the antibody to cell-number ratio, whereas the stronger signals remain unaffected. This finding underscores the importance of maintaining consistent antibody-to-cell-number ratio for comparative studies, such as treatment responses or chromatin-QTL mapping. The spa-ChIP-seq protocol is publicly available, including deck setups, operational parameters, and scripts. We envision that this robust, cost-efficient protocol will facilitate high-throughput, reproducible ChIP-seq analyses, supporting large-scale studies of antibody validation, compound screening, population genomics, and diagnostic frameworks.

## INTRODUCTION

Chromatin immunoprecipitation followed by sequencing (ChIP-seq) is a well-established method for studying the genomic localization of DNA-associated proteins. ChIP-seq is widely used to study histone modifications, chromatin regulators (CRs), transcription factors (TFs), and other DNA-associated proteins (Barski et al. 2007; Johnson et al. 2007; Mikkelsen et al. 2007; Furey 2012). The commonly used ChIP-seq method has led to significant insights about principles of gene regulation and function of the epigenome in both normal and disease conditions (Park 2009; Mundade et al. 2014). While ChIP-seq is useful, the overall experimental workflow involves multiple steps that increase the risk of introducing inconsistency within experiments and between groups, making analysis across ChIP-seq datasets challenging. These challenges were partially addressed by the incorporation of automated liquid handling platforms to improve the robustness of the process (Garber et al. 2012; Blecher-Gonen et al. 2013; Aldridge et al. 2013; Gasper et al. 2014; Busby et al. 2016). Yet, most of the previous efforts (Garber et al. 2012; Blecher-Gonen et al. 2013; Gasper et al. 2014) have automated only a subset of the ChIP-seq steps (**Table 1**). Additionally, the majority of current automated ChIP-seq protocols are limited in their ability to efficiently map non-histone proteins, such as chromatin regulators that may interact only indirectly with DNA (e.g., by being associated with modified histones or histone variants).

**Table 1.** Comparison of spa-ChIP-seq protocol to other automated ChIP-seq approaches. For each liquid handler based ChIP-seq method, the table shows the automated steps, methods for sonication, crosslinking, and library prep, as well as protocol availability, noting the potential shortcomings that limits the high-throughput ability of these approaches: (1) Most of the methods shown only automated a subset of the entire ChIP-seq process, but spa-ChIP-seq (highlighted in grey) is fully automated from cross-linked cells to sequencing ready libraries. (2) For sonication, a probe sonicator shears one tube at a time and thus limits throughput. The Covaris E220 machine is costly and time consuming. Our method uses the PIXUL device (Bomsztyk et al. 2019), which is both low cost ($19/plate vs $640/plate for Covaris E220) and has a short turnaround (∼60-108 min/96 samples vs ∼9.5-10 hours (∼600min) for Covaris E220). (3) The methods shown in the table are optimized for single crosslinking and does not include an option for double crosslinking. (4) All of these methods do not include on-bead library preparation and require reverse-crosslinking and DNA purification after immunoprecipitation prior to sequencing library construction. This leads to an increase in process time and cell loss. Note, we do not discuss here there several variations of ChIP-seq (e.g., AutoRELACS) or other approaches for mapping DNA-associated proteins (e.g., AutoCUT&RUN, AutoCUT&Tag) due to key the differences between the experimental protocols.

| Method | Automated Steps | Sonication | Crosslinking | Library Prep | Protocol Availability |
| --- | --- | --- | --- | --- | --- |
| **HT-ChIP**<br>(Garber et al. 2012; Blecher-Gonen et al. 2013) | Uses Agilent Bravo for library construction and DNA cleanup steps; the rest of the protocol is not automated | Branson Sonifier (probe sonicator) | Single crosslinking with 1% FA | Requires purification beforehand | Automated protocols are available as a written text in the Supplemental Experimental Procedures. |
| **AHT-ChIP-seq**<br>(Aldridge et al. 2013) | Uses Agilent Bravo NGS for ChIP process, Beckman FxP dual arm instrument for pre-PCR library construction, and Caliper Zephyr liquid handler for PCR cleanup | Misonix Sonicator 3000 (probe sonicator) | Single crosslinking with 1% FA | Requires purification beforehand | Agilent Vworks automated ChIP protocol files and Beckman library prep are available in Additional files. |
| **R-ChIP**<br>(Gasper et al. 2014) | Uses Tecan Freedom EVO 200 for ChIP process; library construction and DNA cleanup are not automated | Probe sonicator | Single crosslinking with 1% FA | Requires purification beforehand | Automated protocols are available as text in Methods and Supplementary Information |
| **FloChIP**<br>(Dainese et al. 2020) | Uses an “in house” custom built microfluidic device (non-commercial) for ChIP process; library construction and DNA cleanup are not automated | Covaris E220 | Single crosslinking with 1% PFA | Requires purification beforehand | Automated protocols are available as text in Supplementary Information |
| **spa-ChIP-seq**<br>(this study) | Uses Agilent Bravo NGS for entire ChIP-seq process including sequencing-ready library construction | Active Motif PIXUL | Double crosslinking with 2mM DSG and 1% FA | On-bead library preparation | User-friendly spa-ChIP-seq Agilent Vworks form and protocol files are available in Supplemental files. |

Commonly, ChIP-seq crosslinking involves a single step of formaldehyde (FA) to create covalent bonds between proteins and DNA. However, FA is a small molecule and only captures interactions in the range of 2.3–2.7 Å, which limits the distance of protein interactions captured (Sutherland et al. 2008). Thus, to enable detection of protein interactions occurring between molecules that are further apart, larger FA-compatible crosslinkers, such as disuccinimidyl glutarate (DSG) that can form crosslinks of approximately 7.7 Å, can be combined with FA. Double crosslinking was shown to be a highly effective and reproducible method for capturing the binding of protein complexes or transcription factors (e.g., *NFKB1* and *STAT3*) and identifying genomic targets for protein-DNA binding with high signal-to-noise ratios (Nowak et al. 2005; Tian et al. 2012).

Moreover, effective chromatin solubilization is also critical in the ChIP-seq workflow as it directly influences the immunoprecipitation performance, sonication efficiency, and experiment reproducibility. A wide range of lysis buffers using different detergents have been developed to facilitate nuclear disruption and the release of crosslinked chromatin. These lysis buffers help to generate soluble chromatin preparation, balancing effective lysis and nuclear disruption with preservation of chromatin-protein interactions.

Another important aspect of ChIP-seq workflow is the ratio of antibody to cell-number, as this ratio can influence enrichment efficiency during immunoprecipitation. Usually, the vendor experimentally validates and titrates the antibody to yield an optimal titer for each antibody. Yet, different vendors may have different validation methods, and the available recommendations may not be suitable for all applications (Marx 2019). Using fixed amounts of antibody when cell numbers differ across samples or conditions can lead to suboptimal enrichment, increased background, and variability in ChIP-seq quality (Caride et al. 2023).

We recently developed and published a detailed single-pot ChIP-seq protocol that incorporates an on-bead library preparation step (Texari et al. 2021). Here we adapted this single-pot ChIP-seq protocol to a liquid handler operation and created an end-to-end fully automated version that uses a 96-well sonicator. We first benchmarked our single-pot automated ChIP-seq (spa-ChIP-seq) against manual ChIP-seq to ensure high quality data is produced. The spa-ChIP-seq protocol enables simultaneous processing of a maximum of 96 samples (a full 96-well plate) in a time and cost-effective manner that would not be feasible with manual ChIP-seq. It allowed us to evaluate multiple parameters, including double crosslinking, shearing and crosslinking conditions, lysis buffer compositions, and antibody to cell-number ratios. We demonstrated the utility of our automated method by identifying optimal ChIP-seq conditions for both histone and non-histone proteins. We have streamlined the automation process and made the protocols generalizable, tailored the deck settings and packaged the individual protocols into a user-friendly interface. Our automated ChIP-seq protocol is publicly available, including the specific deck setups, software files and parameters (**Supplemental Files**). We envision that our robust, cost-efficient protocol can advance research efforts via multiple fronts. These include: (***i***) allowing the scaling up the number of replicates and conditions tested in each experiment, (***ii***) improving quantification precision when using spike-in normalization in ChIP-seq experiments (Patel et al. 2024), (***iii***) enabling core facilities to provide high-throughput ChIP-seq as a service, and (***iv***) being incorporated into antibody evaluation procedures (Landt et al. 2012; Wardle and Tan 2015; Mendoza-Parra et al. 2016), compound screening (Zou et al. 2022), population genomics (Waszak et al. 2015) and diagnostic frameworks (Yan et al. 2016).

## RESULTS

### Establishment and evaluation of a single-pot automated (spa)-ChIP-seq protocol

We built on our previous automated ChIP-seq setup with a liquid handling robot (Busby et al. 2016) and modified the parameters to accommodate our recent single-pot ChIP-seq protocol (Texari et al. 2021) to establish spa-ChIP-seq. Both the manual and automated workflows take roughly three days (starting from fixed cells) to produce next generation sequencing (NGS) ready indexed libraries. spa-ChIP-seq workflow automates all steps of the ChIP-seq process with a liquid handler, although it does require the manual preparation of master mixes and buffers prior to the run (**Fig. 1**). While both workflows can process a similar number of samples within a similar timeframe, the incorporation of a liquid handler substantially reduces pipetting and handling errors, minimizing experimental variability, especially in larger-scale experiments. Our automated process is scalable to between 8 to 96 ChIP-seq reactions (ideally in increments of 8) and at a cost of approximately $70 per sample (**Supplemental Table S1**). We first identified multiple experimental variables that could be optimized and tested (**Fig. 1**). We then assessed the performance of our automated ChIP-seq protocol by benchmarking it against the manual ChIP-seq using the exact same input material. To conduct this initial benchmarking, we focused on both H3K27ac, a histone modification that is associated with open chromatin, and H3K27me3, a histone mark that is linked to repressive heterochromatin. The ChIP-seq signal-to-noise ratio for H3K27ac and H3K27me3 were highly concordant between the manual and automated ChIP-seq workflows, validating the liquid handler settings we optimized and the reproducibility of our automated protocol (**Fig. 2A**).

**Fig. 1:**
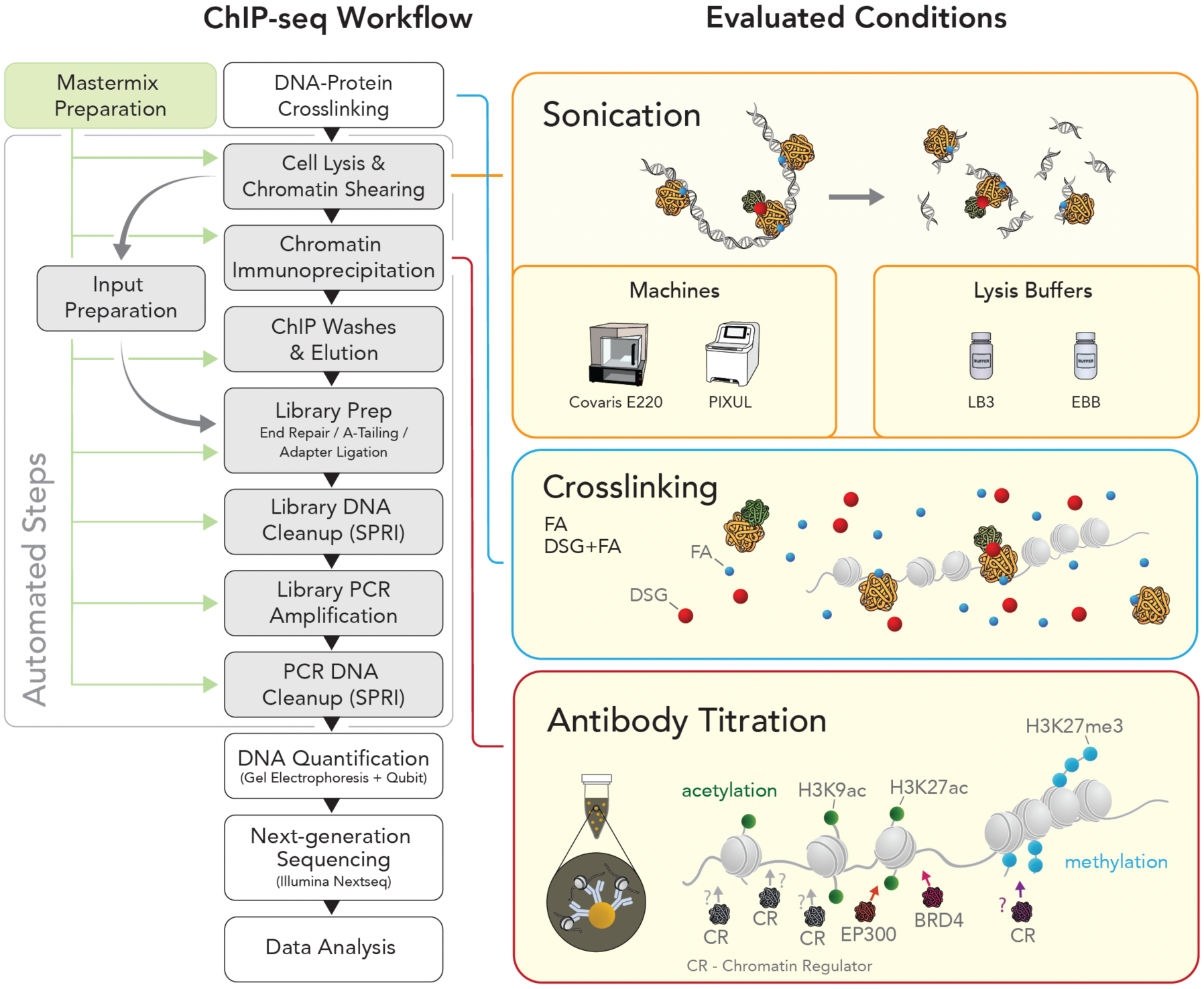
Overview of the spa-ChIP-seq workflow and evaluated ChIP-seq experimental conditions. The automated steps of ChIP-seq workflow are highlighted in the gray boxes of the flowchart on the left and the evaluated experimental conditions in the yellow boxes. The green box shows the required manual preparations of master mixes prior to the start of each automated step. The crosslinking reagent formaldehyde (FA) is represented as a small blue dot, and disuccinimidyl glutarate (DSG) is represented as a large red dot.

**Fig. 2:**
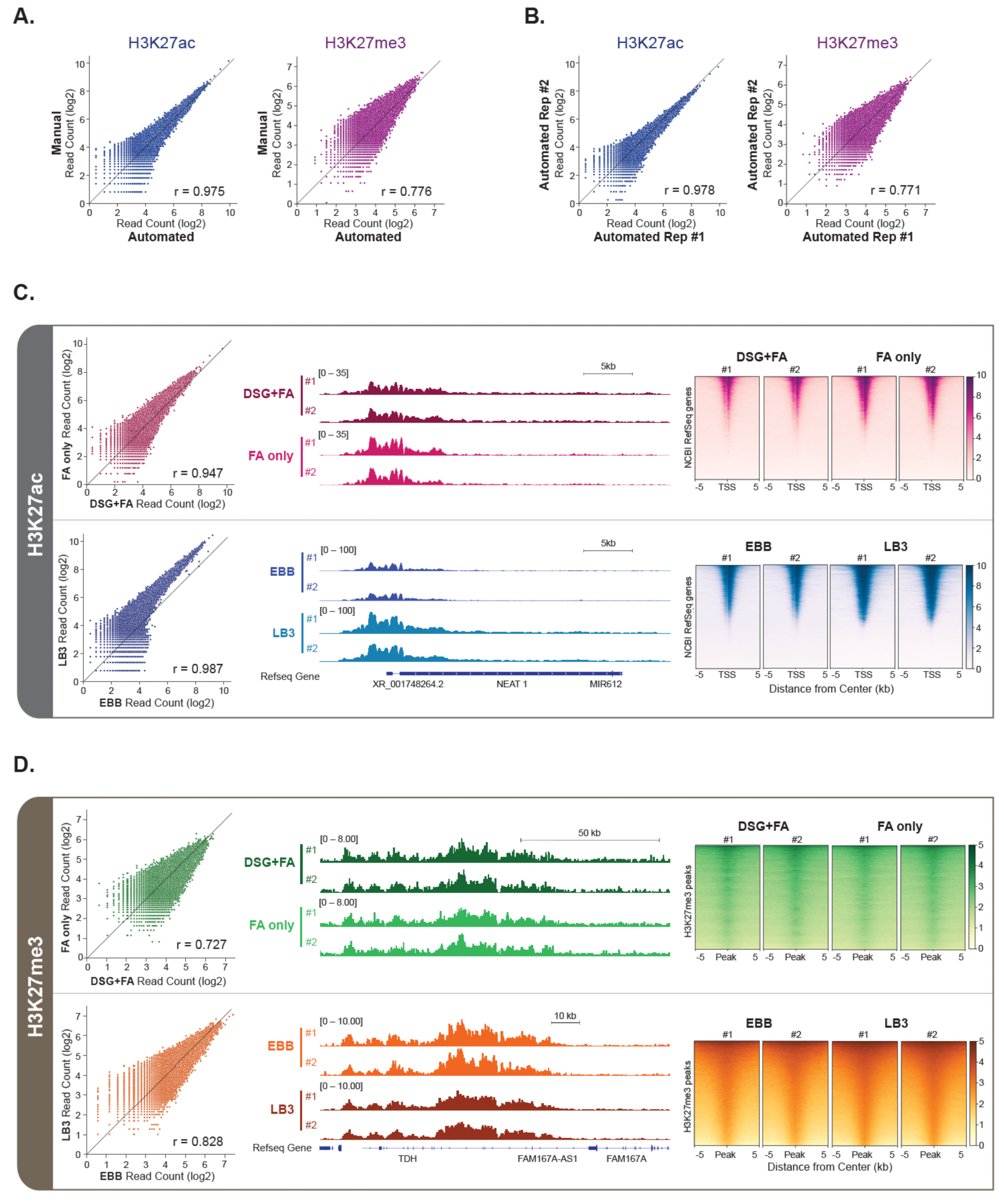
Systematic evaluation of spa-ChIP-seq with comparison to manual workflow, crosslinking conditions and buffer compositions. **A-B.** Scatterplots comparing normalized read counts (log2) between (A) automated and manual ChIP-seq samples and (B) automated replicates at H3K27ac or H3K27me3 shared peaks. **C-D.** Scatterplots of read counts (log2) on shared peaks, IGV (Robinson et al. 2011) genome browser tracks and heatmaps of ±5 kb from the center of transcription start sites (TSSs) or peaks showing the automated ChIP-seq results of (C) H3K27ac and (D) H3K27me3 for evaluating crosslinking conditions, DSG+FA vs FA only (top), and buffer compositions, EBB vs LB3 (bottom). The Pearson r correlation coefficients are reported on the lower right corner of the scatterplots. A total of 32 ChIP-seq experiments were performed, 16 of them using spa-ChIP-seq and 16 of them using manual ChIP-seq, with additional data in **Supplemental Fig. S2**.

### Identification of optimal shearing conditions, buffer composition and cross-linking using spa-ChIP-seq

Most of the initial automation efforts (Garber et al. 2012; Blecher-Gonen et al. 2013; Aldridge et al. 2013) has focused on using a probe sonicator such as Branson, which is limited to shearing one sample at a time. Here, we aimed to use a 96-well format sonicator. In previous work (Busby et al. 2016), we used the Covaris E220 device that allows shearing of up to 96 samples. However, as the Covaris E220 sonicates only one sample at a time, shearing of 96 samples requires approximately 10 hours. This limitation both leads to the samples being kept for a long time within the Covaris E220 before the addition of antibodies, as well as extends the time it takes to perform the protocol. A recently introduced sonication device, PIXUL (Bomsztyk et al. 2019) enables the shearing of 96 samples in only 1-2 hours, and also reduces the costs of the plate used for sonication by 30-fold (**Supplemental Table S1**). We decided to evaluate the ability of PIXUL to reproducibly shear samples in a manner that is at least comparable to the Covaris E220. For both devices we additionally tested the impact of placing samples at different locations within the 96-well plate (**Supplemental Fig. S1A**). While the samples sheared by the different devices had distinct size distributions (∼250 bp for Covaris E220 and ∼350 bp for PIXUL), both were within the range of 200-600 base pairs that is normally used for ChIP-seq (**Supplemental Fig. S1A**). As the PIXUL machine was significantly faster and about 30-fold lower in costs per plate, we proceeded to use this device for shearing in our tests of subsequent variables using our spa-ChIP-seq.

Next, we compared the ChIP-seq results of the histone modifications H3K27ac and H3K27me3 under single and double crosslinking conditions as well as two different lysis buffer compositions, EBB and LB3 (**Methods**; **Fig.2C-D**; **Supplemental Fig. S2**). We used PIXUL to shear approximately 1 million crosslinked HeLa-S3 cells per well; to reduce variation, following the shearing step we merged the sheared chromatin from all wells. Next, we split the pool of sheared chromatin between the two conditions (manual or automated ChIP-seq). In total, for this initial evaluation, we performed 32 ChIP-seq experiments, comparing 16 different conditions, each in duplicates.

We observed that the samples treated with the additional crosslinker required longer sonication time in LB3, the lysis buffer we routinely use (**Supplemental Fig. S1B**). We evaluated the ability of EBB buffer to assist the shearing process. Sonication of the double crosslinked samples in EBB for 60 minutes provided optimal fragmentation patterns. This fits with prior work (Métivier et al. 2003). We overcame the inefficient fragmentation of double crosslinked samples in LB3 buffer (**Supplemental Fig. S1B**) by increasing the sonication time to 90-120 minutes. The longer shearing time provided optimal shearing patterns. Thus, both buffers were able to generate fragment sizes of 200-600bp and were subsequently used to determine their performance in ChIP-seq.

Next, we used the double crosslinked samples and performed ChIP-seq of H3K27ac and H3K27me3, either lysed with EBB or LB3. Overall, we observed that the samples sheared in EBB had lower signal-to-noise ratio (SNR) compared to the ones sonicated in LB3, potentially due to the high SDS levels in the EBB buffer (1% SDS; **Methods**; **Fig. 2C-D**; **Supplemental Fig. S2A-D, G-H**). Given the lower SNR we obtained when using EBB, we conducted the rest of the experiments only with LB3 buffer and sonicated the double crosslinked samples for 108-120 minutes. Using these conditions, the H3K27me3 and H3K27ac ChIP-seq results from the double crosslinked samples were mostly indistinguishable from the single crosslinked ones, providing evidence that the additional crosslinker or longer shearing time did not introduce bias in either active or repressive chromatin environments (**Fig. 2C-D**; **Supplemental Fig. S2A-F**). Since double crosslinking with DSG and FA extends the range of captured interactions in ChIP-seq of DNA-associated proteins and does not negatively impact histone modification ChIP-seq results, using double-crosslinked cells for both target types offer a feasible and consistent protocol for experiments involving chromatin regulators, transcription factors and histone marks.

### Identification of optimal conditions for genomic mapping of non-histone DNA-associated proteins

Next, we used the ChIP-seq conditions identified above to map non-histone proteins in double-crosslinked (DSG+FA) HeLa-S3 cells. As a proof of concept, we focused on 5 DNA-associated proteins (CTCF, HAT1, HDAC3, RPB1 (POLR2A), and YAP1) to perform ChIP-seq, each in two technical replicates (**Fig. 3** and **Supplemental Fig. S3**). Following our previous efforts to increase ChIP-seq standardization (Busby et al. 2016), we aimed to mainly focus on monoclonal antibodies and used our automated approach to evaluate between 2-4 different antibodies for each target protein. Technical replicates were highly reproducible SNR across six of the seven samples and the variability observed in the remaining sample was within the expected range of experimental variability (**Fig. 3** and **Supplemental Fig. S3**). Together, we were able to map all of the DNA-associated proteins we tested, although for a subset of the antibodies we obtained only a weak signal under the set of conditions we evaluated (**Supplemental Table S2**).

**Fig. 3:**
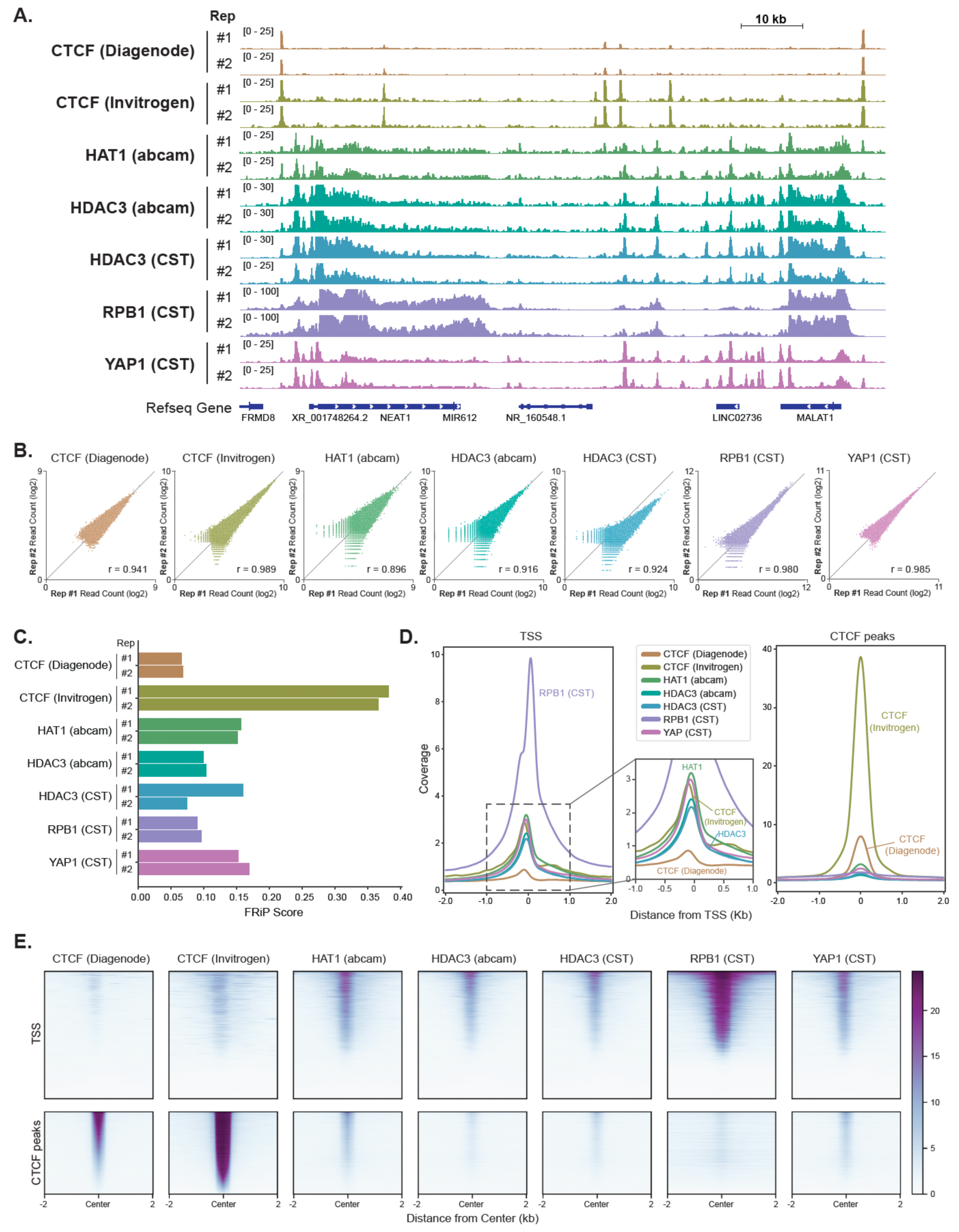
Mapping of DNA-associated proteins using spa-ChIP-seq. **A.** IGV genome browser tracks showing the ChIP-seq of DNA-associated proteins: CTCF, HAT1, HDAC3, Rpb1, and YAP1. CTCF and HDAC3 ChIP-seq experiments were performed with two antibodies from two different vendors. Each antibody condition has two technical replicates (labeled as #1 and #2), with a total of 14 ChIP-seq samples. **B.** Scatterplots showing the comparison of the 2 technical replicates at shared peaks for each antibody condition. **C.** Bar plot showing Fraction of Reads in Peaks (FRiP) for the ChIP-seq of DNA-associated proteins. For each antibody, both technical replicates (labeled as #1 and #2) are plotted. **D.** Histograms showing the average normalized read coverage of the replicates at ± 2 kb from the center of TSSs and CTCF peaks. **E.** Heatmaps showing the average normalized read density of the replicates at ±2 kb from the center of TSSs (top row) and CTCF peaks (bottom row).

### The antibody to cell-number ratio may impact ChIP-seq results

Another important, yet often overlooked, aspect of ChIP-seq is the ratio between the amount of antibody used for immunoprecipitation and the number of cells (or amount of chromatin) in the reaction. Our automated ChIP-seq protocol significantly simplifies the task of performing multiple antibody titrations which could be tedious otherwise. We first titrated H3K9ac antibody in HeLa-S3 cells with four different amounts of the antibody, each in replicates. Initial evaluation of the genome browser tracks on IGV and FRiP scores show that different amounts of antibodies had minimal impact on the global ChIP-seq signal, and the overall trend is that higher amounts of antibodies may lead to only slightly higher ChIP enrichment (**Supplemental Fig. S4**). However, when we equally divided the ChIP-seq peaks into four quartiles based on the normalized read counts at peaks, we found that the amount of antibody had a greater impact on ChIP-seq peaks with weaker enrichment, whereas the enrichment signal is constant for the strongest peaks in the experiment (**Fig. 4A**).

**Fig. 4:**
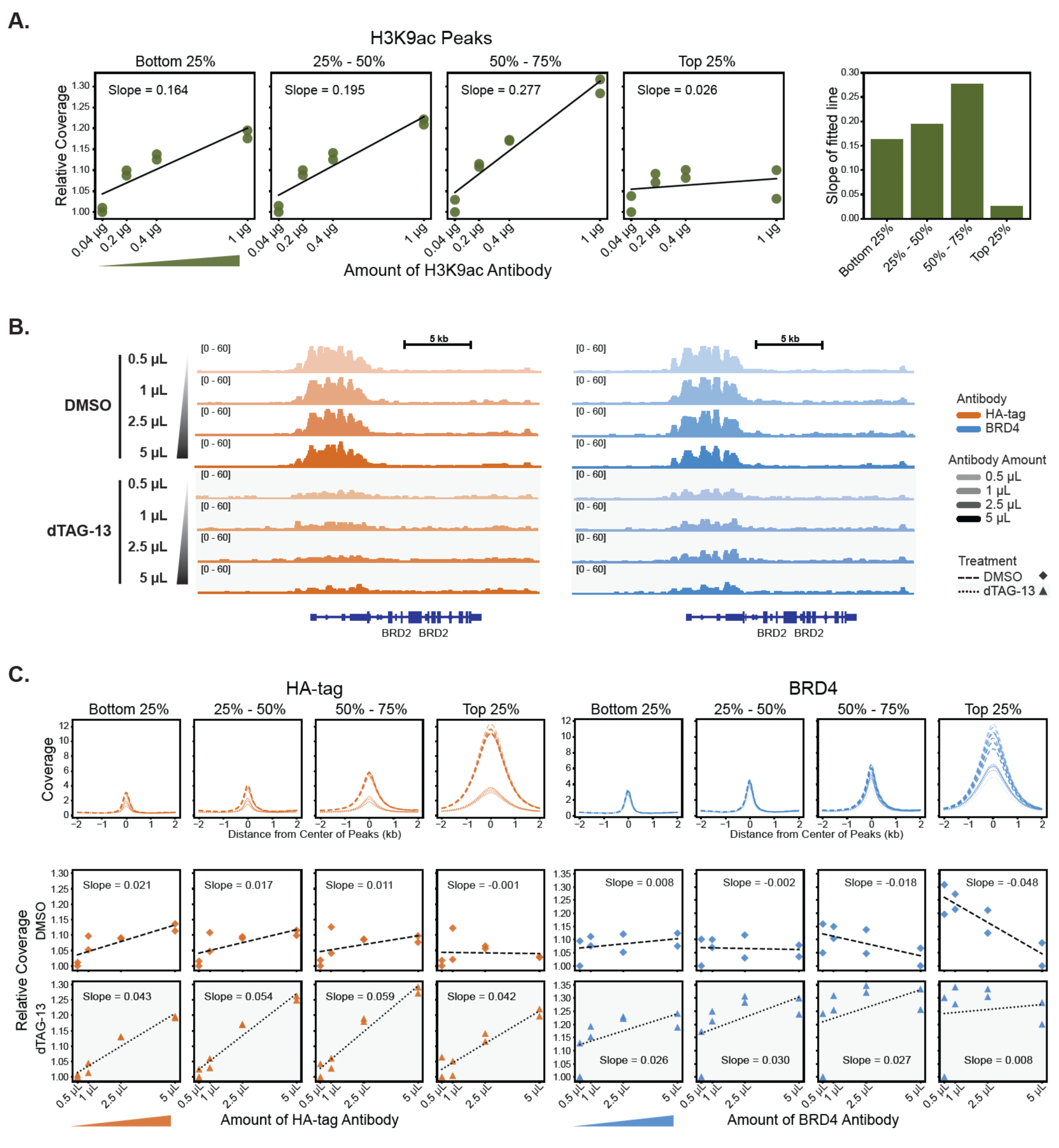
Antibody titration with spa-ChIP-seq reveals the weaker signals are more sensitive to the amount of antibody. **A.** Relative total coverage of H3K9ac signal with 0.04 μg, 0.2 μg, 0.4 μg, and 1 μg of antibody for H3K9ac peaks separated into quartiles based on normalized read counts at peaks (the values are relative to the minimum, which is set to 1). Each condition has 2 technical replicates, with a total of 8 ChIP-seq samples. Each plot is fitted with a linear regression line, and the corresponding slope of fitted line is shown in the bar plot on the left. **B.** IGV genome browser tracks for HA-tag and BRD4 ChIP-seq in dTAG-13 and DMSO treated HEK293T-BRD4-FKBP12^F36V^-2xHA cells with 0.5 μL, 1 μL, 2.5 μL, and 5 μL of the HA-tag and BRD4 antibodies, with each amount of antibody performed in replicates for a total of 32 ChIP-seq samples. The tracks of 2 replicates for each condition are overlaid on one another. **C.** Histograms of normalized read coverage for HA-tag and BRD4 antibody titration ChIP-seq signals centered on peaks obtained by using the DMSO samples. The peaks are separated into quartiles according to normalized read counts at peaks. Relative total coverage of HA-tag and BRD4 signals with 0.5 μL, 1 μL, 2.5 μL, and 5 μL of antibody for each quartile of DMSO peaks shown in the histogram above. The orange and blue triangles represent the increase in the ratio of antibody to cell-number in HA-tag and BRD4, respectively. Each condition has 2 technical replicates.

We hypothesized that the weaker signals are more sensitive to the amount of antibody, and to investigate this further, we incorporated a cell line with a knock-in that allows an inducible degradation of the BRD4 protein (HEK293T-BRD4-FKBP12^F36V^-2xHA (Nabet et al. 2018)). Upon treatment of dTAG-13, the amount of BRD4 epitopes is greatly reduced (Nabet et al. 2018). We performed titration experiments for four different amounts of antibodies targeting HA-tag and BRD4 in both dTAG-13 and DMSO treated cells. Regardless of the amounts of antibody used, the ChIP-seq signal is reduced in dTAG-13 treated cells for both the HA-tag and BRD4 antibodies (**Fig. 4B**). When the DMSO peaks for each antibody were separated by quartiles based by their DMSO enrichment, the dTAG-13 treated samples showed an increasing higher signal as the ratio of antibody to cell-number increased in all 4 quartiles regardless of the trend in the DMSO samples (**Fig. 4C**).

Next, to further study our hypothesis regarding the weaker signals being more sensitive to the antibody to cell-number ratio, we focused on the EP300, that binds strongly to non-promoter cis-regulatory elements and is weakly associated with promoter regions (Heintzman et al. 2007; Mendenhall and Bernstein 2008). Using spa-ChIP-seq we conducted a comprehensive antibody titration experiment, where we varied both the number of cells and the amounts of EP300 antibody (**Fig. 5A**). In particular, we titrated 0.05 μg, 0.5 μg, 2.5 μg, and 5 μg of EP300 antibody with 20K, 200K, and 2M of HEK293T cells. To minimize variability, we used the same chromatin preparation that was split across the different conditions. We observed that, in general, the higher amount of antibody and higher number of cells tend to generate better ChIP-seq signal to noise ratios. Yet, when comparing the normalized read counts on shared EP300 peaks for the different combinations of antibody to cell-number ratios with each other, we noticed that samples with distinct ratios did not share similar set of peaks, and the Pearson r correlation coefficients varied across the combinations (**Fig. 5A**). Thus, given that the same original chromatin extract was used for all conditions, the ratio of EP300 antibody to cell-number appears to cause a discrepancy in the ChIP-seq results.

**Fig. 5:**
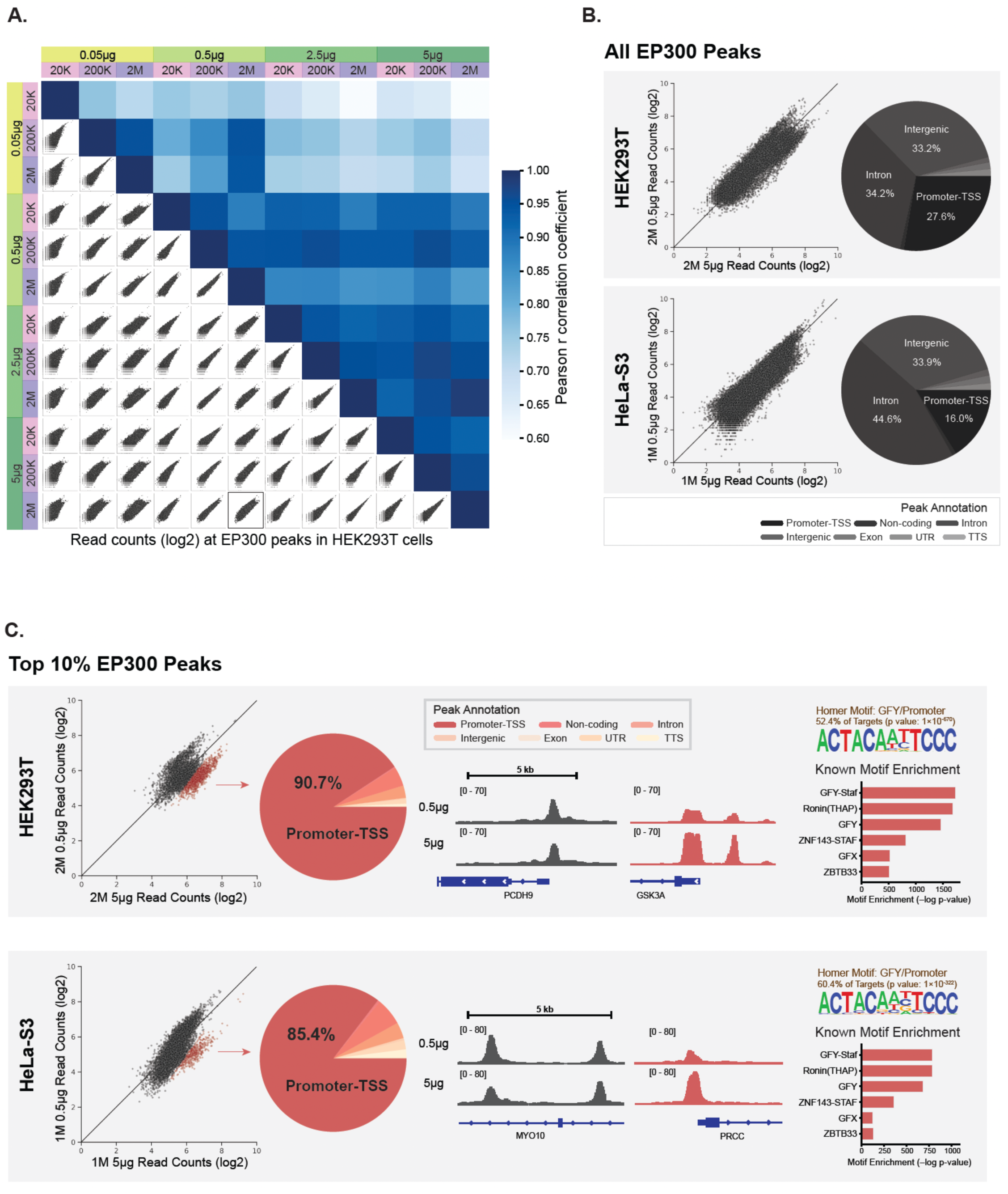
EP300 antibody titration shows that higher antibody to cell-number ratio better captures weaker promoter signals. **A.** A matrix of scatterplots showing normalized read counts (log2) at EP300 peaks and heatmaps showing Pearson r correlation coefficients for EP300 antibody titrations in HEK293T cells with various antibody to cell-number ratios. We conducted a comprehensive comparison of conditions using 0.05 μg, 0.5 μg, 2.5 μg, and 5 μg of EP300 antibody (green shades) and 20K, 200K, and 2M of HEK293T cells (pink/purple shades), resulting in a total of 12 ChIP-seq samples. The scatterplot outlined in bold corresponds to the HEK293T sample shown in (B). Note that a similar matrix for HeLa-S3 cells is presented in **Supplemental Fig. S6B**. For HeLa-S3 samples, each antibody amount was tested in replicates, yielding a total of 8 ChIP-seq samples, and the replicates for each amount were merged for subsequent analysis. **B.** Scatterplot of normalized read counts (log2) at all EP300 peaks comparing 0.5 μg and 5 μg of antibody in 2M HEK293T cells (top) and 1M HeLa-S3 cells (bottom). The pie chart on the right shows the genomic annotations for all EP300 peaks. The peaks are annotated with HOMER and the annotations include promoter-TSS, non-coding, intron, intergenic, exon, UTR (5’ UTR and 3’ UTR), and TTS (see legend below the pie chart). **C.** Scatterplot of normalized read counts (log2) at the top 10% of EP300 ChIP-seq peaks comparing 0.5 μg and 5 μg of antibody in 2M HEK293T cells (top) and 1M HeLa-S3 cells (bottom). The subset of peaks that has higher signal in 5 μg antibody than 0.5 μg antibody is highlighted in red. The pie chart shows the distribution of peak annotations in the red subset highlighted in the scatterplot to the left. IGV genome browser tracks show 2 peak loci in the red subset. HOMER motif analysis identified same top enriched known motifs in the red subset of peaks for both HeLa-S3 and HEK293T cells. The motif enrichments are shown in the bar plot as –log p-values.

We conducted a similar antibody titration experiment using a different cell line, HeLa-S3. Here, we only varied the amount of antibody (ranging between 0.5 μg, 1 μg, 2.5 μg and 5 μg of EP300 antibody, **Supplemental Fig. S6**) with 1M HeLa-S3 cells, splitting the exact same chromatin extract across the different antibody amounts to minimize variability, as described above. In a similar manner to our observations in HEK293T cells, we noticed a discrepancy between the genomic patterns between the different ratios of antibody to cell-number (**Supplemental Fig. S6B**).

In both experiments – either when we used HEK293T or HeLa-S3 – we noticed that the major discrepancy between antibody to cell-number ratios was observed in the regions with the higher EP300 read counts, in particular when comparing the lowest vs. the highest ratios (0.5 µg vs 5 µg EP300 antibody per 1M HeLa-S3 cells, respectively, **Fig. 5B**). To further study this discrepancy, we focused on the top 10th quantile and compared the normalized read counts between highest and lowest antibody to cell-number ratios. This analysis revealed a subset of peaks with stronger enrichment in the higher antibody to cell-number ratio (**Fig. 5C**). Next, we used PCA (principal component analysis) to separate out the subset of peaks that showed an increased signal in samples with higher ratio of antibody to cell-number in both cell lines (**Supplemental Fig. S6D-E**). Annotation of the loci associated with the subset of peaks impacted by the antibody to cell-number ratio showed that these regions are predominantly present at promoter-TSS regions with 90% in HEK293T cells and 85% in HeLa-S3 cells, whereas the annotation of the entire set of EP300 peaks showed high localization at non-promoter intergenic or intron regions, and only 27% (HEK293T) and 16% (HeLa-S3) in promoter regions (**Fig. 5B-C**). Next, to better define the subset of peaks most impacted by the antibody to cell-number ratio, we conducted motif analysis, which identified a GFY motif commonly associated with some active promoter regions. Moreover, in both cell lines, this subset of EP300 peaks was enriched for the same top known motifs (**Fig. 5C**) further supporting the notion that the weaker or less frequent localization of EP300 at promoter regions leads to the higher sensitivity of these peaks to the ratio of antibody to cell-number.

## DISCUSSION

We presented here spa-ChIP-seq, a single-pot automated protocol for chromatin profiling. We benchmarked spa-ChIP-seq by performing parallel manual ChIP-seq reactions and demonstrated a high reproducibility and nearly indistinguishable signal-to-noise ratio between manual and automated workflows. We then used spa-ChIP-seq we systematically evaluate several aspects of the ChIP-seq process, including shearing conditions, crosslinking methods, and buffer compositions.

In our hands, the optimal conditions included (***i***) shearing with the Active Motif PIXUL device (Bomsztyk et al. 2019), that provides a short incubation time and low cost per sample, (***ii***) lysing the cells with LB3. Although this buffer requires longer sonication time compared to EBB (potentially due to the weaker reagent), the ChIP-seq results had higher signal-to-noise ratio than EBB, (***iii***) using DSG+FA double crosslinking for all ChIP-seq profiling to allow a consistent the experimental procedure regardless of the target epitope. We demonstrated that for histone modifications the signal-to-noise ratio was indistinguishable between single (FA) and double (DSG+FA) crosslinking. As non-histone DNA-associated proteins such as chromatin regulators me be less amenable to get crosslinked to the DNA, double crosslinking by DSG and FA can yield ChIP-seq results with robust signal-to-noise ratios. While FA only forms crosslinks of 2.3–2.7 Å, DSG can form crosslinks of 7.7 Å thus potentially capturing longer interactions (Sutherland et al. 2008). The combination of DSG and FA, DSG first stabilizes the longer protein-protein interactions and then FA crosslinks shorter DNA-protein interactions (Nowak et al. 2005; Tian et al. 2012).

Using spa-ChIP-seq and our optimal set of conditions, we identified a set of mostly monoclonal antibodies for an array of non-histone DNA associated proteins, including CTCF, BRD4, HA-tag, HDAC3, HAT1 and YAP1. As we discussed in our previous work (Busby et al. 2016), the use of monoclonal antibodies is critical to allow reproducibility of studies e.g., between groups or over time, as polyclonal reagents are non-renewable and each batch may differ from the other. We thus envision that the validation of this set of antibodies and conditions to allow reproducible studies of these targets.

A key observation of our study relates to the impact of antibody to cell-number ratio in chromatin profiling. Using spa-ChIP-seq we conducted a set of comprehensive experiments to systematically interrogate the effect of this ratio. For each experiment, we used the exact same chromatin prep, thus only varying the ratios of the antibodies or the number of cells. First, we demonstrated that only the weaker quartiles of H3K9ac peaks are impacted by the antibody to cell-number ratio, while the upper quartile is not. We then used a BRD4 HA-tagged perturbation system to reduce the amount of epitope by degrading HA-tagged BRD4 and show that when we titrate either the HA-tag or BRD4 antibodies following depletion of the target, the normalized ChIP enrichment signal is impacted by the antibody to cell-number ratio much more then prior to depletion. Lastly, we focused on EP300 that has both strong (enhancer/cis regulatory elements) and weak (promoter) binding sites (Heintzman et al. 2007; Mendenhall and Bernstein 2008) and systematically titrated both the number of cells and the amount of antibody in two cell lines. This allowed us to identify a subset of the peaks that show enhanced sensitivity to the antibody to cell-number ratio and show that they are highly enriched for promoter signatures. Thus, our results highlight the importance of maintaining the ratio of antibody to cell-number as constant as possible, in particular in chromatin profiling experiments aimed at studying the impact of perturbations (e.g., by small molecules or CRISPRi) to allow proper comparisons.

Altogether our study highlights the usefulness of spa-ChIP-seq in conducting comprehensive, systematic studies. The method reduces hands-on time and cost per sample, and the increased throughput (up to 96 samples) were key components in our ability to conduct our systematic studies. We anticipate that spa-ChIP-seq will allow the scaling up of ChIP-seq studies to improve statistical power and facilitate novel experimental work including assaying entire population cohorts and large compound libraries as well as potentially advance diagnostic approaches.

## METHODS

### Cell culture and fixation (crosslinking)

HeLa-S3 (ATCC CCL-2.2™), HEK293T (CRL-3216™), and HEK293T-BRD4-FKBP12^F36V^-2xHA adherent cells were cultured in Dulbecco’s Modified Eagle’s Medium (Gibco™ DMEM, high glucose, GlutaMAX™ Supplement, with 4.5 g/L D-Glucose), supplemented with 10% fetal bovine serum (FBS), 110mg/L sodium pyruvate, 2 mM L-glutamine, and 1% Antibiotic-Antimycotic. Cells were incubated at 37°C with 5% CO2 and 95% Air. HEK293T-BRD4-FKBP12^F36V^-2xHA cells were treated with 500 nM of dTAG-13 (Tocris, cat #6605) or DMSO for 24 hours. When the cells were grown to confluency, they were either single or double crosslinked. For single crosslinking, cells were incubated with 1% formaldehyde solution for 10 minutes at room temperature. For double crosslinking, cells were incubated first with 2mM DSG solution for 30 minutes and then with 1% formaldehyde solution for 10 minutes at room temperature. The crosslinking reaction was quenched by 2.625 M glycine to a final concentration of 0.125 M. The cell pellets were washed with PBS and stored in -80°C.

### Chromatin shearing

Cells were lysed with either the LB3 lysis buffer (10 mM Tris-HCl pH 7.5, 100 mM NaCl, 1 mM EDTA, 0.1% Deoxycholate, 0.5% N-Lauroylsarcosine, 1× protease inhibitor cocktail), or the EBB lysis buffer (0.5% Empigen BB (Sigma-Aldrich, #30326), 1% SDS, 50 mM Tris-HCl pH 7.5, 10 mM EDTA, 1× protease inhibitor cocktail; Métivier et al. 2003). Cells were sonicated either on the Active Motif PIXUL sonicator for 60-120 minutes per plate, or on the Covaris E220 sonicator for 6 minutes per well. After sonication, lysates in EBB lysis buffer were diluted 2.5-fold in EBB dilution buffer (20 mM Tris-HCl pH 7.5, 100 mM NaCl, 0.5% Triton X-100, 2 mM EDTA, 1× protease inhibitor cocktail).

### Analysis of sheared chromatin / input preparation

#### Manual workflow

Per input sample, a reaction mix (0.5% SDS, 18.75 mM EDTA, 280 mM NaCl, 250 μg/mL Proteinase K, 125 μg/mL RNase A) was added and incubated in a thermocycler at 55°C for 1 hour and then 65°C for at least 2 hours. To clean up the input sample, Speedbead mastermix (2 μL SpeedBeads (Cytiva # 65152105050250), 12% PEG8000 final, 1 M NaCl final) was added to each input and incubated at room temperature for 10 minutes. Then, the input sample was washed twice with 80% EtOH and the Speedbeads pellet was air-dried. Lastly, the input sample was eluted with TT buffer (0.05% Tween 20, 10 mM Tris pH 8.0).

#### Bravo workflow

Reaction mix plate was prepared by equally distributing the total required amount of reaction mix to each of the well in the first column and placed on Bravo deck 6. After running Bravo protocol *1a_Input_Prep*, the input plate was incubated in a thermocycler at 55°C for 1 hour and then 65°C for at least 2 hours. Speedbeads mastermix plate was prepared by equally distributing the total required amount of Speedbeads mastermix to each well in the first column and placed on Bravo deck 5. After incubation, the input plate was placed back on Bravo deck 4, and Bravo protocol *1b_Input_PEG* was run. Ethanol reservoir was placed on deck 9, and TT buffer reservoir was placed on deck 3. Immediately after incubation with Speedbeads mastermix, Bravo protocol *1c_Input_Speedbead_Cleanup* was run to clean up the input samples. At the end of the protocols, the input plate should contain input samples eluted in TT buffer.

### Chromatin immunoprecipitation

See **Supplemental Table S2** for all the antibodies used in the immunoprecipitation step.

#### Manual workflow

Dynabeads™ Protein A or G were resuspended either in LB3 and 1 % Triton X-100 for LB3 ChIP samples or in EBB dilution buffer for EBB ChIP samples. Resuspended beads were combined with each antibody to make antibody/beads mix. Per each ChIP sample, sheared chromatin was combined with the antibody/bead mix.

#### Bravo workflow

The antibody/beads mix was made the same way as the manual workflow. The mix was transferred to tube strips, which were then placed on deck 4. PIXUL lysate plate was placed on deck 5. Bravo protocol *2_Chromatin_Beads_Antibody_On_Mag* was run to combine the chromatin lysate with the antibody/beads mix.

The ChIP samples were incubated overnight, rotating overhead on HulaMixer™ Sample Mixer at 4°C in the cold room.

### ChIP Washes

#### Manual workflow

After overnight incubation, the sample was washed three times with WBI (20mM Tris pH7.5, 1% Triton X-100, 2 mM EDTA, 150 mM NaCl, 0.1% SDS, 1× protease inhibitor cocktail), three times with WBIII (10 mM Tris pH7.5, 0.7% DOC, 1 mM EDTA, 250mM LiCl, 1× protease inhibitor cocktail), and twice with TET (10 mM Tris pH7.5, 0.2% Tween-20, 1 mM EDTA, 1× protease inhibitor cocktail). The Dynabeads loaded with each ChIP sample were resuspended in 25 μL of TT buffer (10 mM Tris-HCl pH 8.0, 0.05% Tween 20).

#### Bravo workflow

Excessive amounts of wash buffers (WBI, WBIII, and TET) and TT buffer were prepared. Waste reservoir was placed on deck 1, elution TT buffer reservoir was placed on deck 3, empty ChIP plate was placed on deck 4, IP tube strips were placed on deck 6, and wash buffer reservoir was placed on deck 9. Bravo protocol *3_ChIP_Wash_Elution_Off_Mag was* run. At the end of the protocol, each well of the ChIP plate should contain 25 μL of ChIP sample/Dynabeads resuspended in TT buffer.

### Library construction, reverse crosslinking, and library cleanup

For the IP and input samples, libraries were constructed with the NEBNext® Ultra™ II DNA Library Prep Kit for Illumina (Catalog #E7645L). Indexing adapters were from NEXTflex DNA Barcodes: 48 (NOVA-514104) and NEXTflex Unique Dual Index Barcodes (8NT index, 97-192) (NOVA-514151).

#### Manual workflow

**(1) End prep:** End prep mastermix (NEBNext Ultra II End Prep Enzyme Mix and NEBNext Ultra II End Prep Reaction Buffer) was added to the ChIP and input samples and incubated for 30 minutes at 20°C, and then 30 minutes at 65°C in a thermal cycler. **(2) Adapter ligation:** 0.625 μM of indexing adapters and ligation mastermix (NEBNext Ultra II Ligation Master Mix and NEBNext Ligation Enhancer) were added to the ChIP and input samples, which were then incubated for 15 minutes at 20°C. **(3) Reverse crosslinking:** Per sample, ligation reaction was stopped by adding 27μL of ligation stop solution mastermix (0.5% SDS, 18.75 mM EDTA, 250 μg/mL Proteinase K, 125 μg/mL RNase A) and 285 mM NaCl. ChIP samples were incubated for 1 hour at 55°C and then at least 2 hours at 65°C. **(4) Library cleanup:** The supernatant of each sample was separated from Dynabeads Protein A/G and incubated with Speedbeads mastermix (2 μL SpeedBeads, 8.6% PEG8000 final, and 0.8 M NaCl final) for 10 minutes at room temperature. Then, the input sample was washed twice with 80% EtOH and the Speedbeads pellet was air-dried. Lastly, the input sample was eluted with TT buffer. <u>Bravo workflow:</u> **(1) End prep:** End prep mastermix plate was prepared by equally distributing the total required amount of the mastermix to each well in the first column and placed on Bravo deck 6. The ChIP sample plate was placed on deck 4. Bravo protocol *4_End_Repair_MM_Dispense* was run and the ChIP plate was incubated for 30 minutes at 20°C, and then 30 minutes at 65°C in a thermal cycler. **(2) Adapter ligation:** Ligation mastermix plate was prepared by equally distributing the total required amount of the mastermix to each of the well in the first column and placed on Bravo deck 6. The ChIP sample plate was placed on deck 4, and the index plate was placed on deck 5. Bravo protocol *5_Adapter_Ligation_MM* was run, and the ChIP plate was incubated for 15 minutes at 20°C. **(3) Reverse crosslink:** Ligation stop solution mastermix plate was prepared by equally distributing the total required amount of the mastermix to each of the well in the first column and placed on Bravo deck 6. The ChIP sample plate was placed on deck 4, and the 5 M NaCl reservoir was placed on deck 9. was prepared as a Buffer Plate and the plate was placed on deck 5. Bravo protocol *6_Ligation_Stop_Solution_Disp* was run and the sample was incubated for 1 hour at 55°C and then at least 2 hours at 65°C. **(4) Library cleanup:** Speedbeads mastermix plate was prepared by equally distributing the total required amount of Speedbeads mastermix to each of the well in the first column and placed on Bravo deck 5. After incubation, the ChIP plate was placed back on Bravo deck 4, and Bravo protocol *7a_Adapter_Ligation_Cleanup* was run. Ethanol reservoir was placed on deck 9, and TT buffer reservoir was placed on deck 3. Immediately after incubation with Speedbeads mastermix, Bravo protocol *7b_Adapter_Ligation_Cleanup* was run to clean up the input samples. At the end of the protocols, the ChIP plate should contain ChIP samples eluted in TT buffer.

### PCR Amplification and final cleanup of ChIP and Input libraries

#### Manual workflow

**(1) PCR amplification:** The PCR mastermix [Ultra II Q5 Mastermix, and 1 μM sequencing platform-compatible forward and reverse primers (Solexa 1GA and Solexa 1GB)] was added to each library sample. A PCR program (Initial denaturation at 98°C for 30 seconds; 12 cycles of denaturation at 98°C for 10 seconds, annealing at 60°C for 15 seconds, extension at 72°C for 30 seconds; and final extension at 72°C for 2 minutes) was run to amplify the libraries using a thermal cycler. **(2) PCR cleanup:** The PCR products were incubated with Speedbeads mastermix (2 μL SpeedBeads, 8.5% PEG8000 final, and 1.06 M NaCl final) for 10 minutes at room temperature. Then, the input sample was washed twice with 80% EtOH and the Speedbeads pellet was air-dried. Lastly, the input sample was eluted with TT buffer.

#### Bravo workflow

**(1) PCR amplification:** PCR mastermix plate was prepared by equally distributing the total required amount of the mastermix to each of the well in the first column and placed on Bravo deck 6. The ChIP library plate was placed on deck 4. Bravo protocol *8_PCR_Enrichment_MM_Dispense* was run. Then, a PCR program (Initial denaturation at 98°C for 30 seconds; 12 cycles of denaturation at 98°C for 10 seconds, annealing at 60°C for 15 seconds, extension at 72°C for 30 seconds; and final extension at 72°C for 2 minutes) was run to amplify the libraries using a thermal cycler. **(2) Library cleanup:** Speedbeads mastermix plate was prepared by equally distributing the total required amount of Speedbeads mastermix to each of the well in the first column and placed on Bravo deck 5. After incubation, the ChIP library plate was placed back on Bravo deck 4, and Bravo protocol *9a_PCR_Enrichment_Cleanup* was run. Ethanol reservoir was placed on deck 9, and TT buffer reservoir was placed on deck 3. Immediately after incubation with Speedbeads mastermix, Bravo protocol *9b_PCR_Enrichment_Cleanup* was run to clean up the input samples. At the end of the protocols, the ChIP library plate should contain ChIP library samples eluted in TT buffer.

### DNA Quantification and sequencing

The size of the DNA libraries was estimated by running on a 2% agarose gel pre-stained with GelGreen. DNA libraries should have a smear with high intensity around 250-500 bp on the gel (**Supplemental Fig. S7**). The concentration of the library (ng/μL) was determined using a Qubit fluorometer with Qubit HS DNA assay kit (Invitrogen). The libraries were pooled according to the concentration calculations. The pooled libraries were prepared and sequenced on Illumina NextSeq 500 or NextSeq 2000.

### Sequencing data analysis

Samples were demultiplexed with Illumina’s bcl2fastq program, adapters were removed with Trimmomatic (Bolger et al. 2014), and FastQC was run for sequencing quality control. Reads were mapped to human reference genome hg38 using Bowtie2 (Langmead and Salzberg 2012) with default parameters and duplicated reads were removed with SAMtools (Li et al. 2009) (samtools markdup). HOMER (Heinz et al. 2010) was used to make tag directories, which contain filtered read counts for primary alignments with MAPQ > 10. HOMER makeBigWig was used to create bigwig files for visualization on IGV genome browser (Robinson et al. 2011). Peaks were called with HOMER findPeaks program with parameters “-style histone”, “-style histone -size 1000 -minDist 2500” for broad peaks, and “-style factor” for transcription factors, using an input control as background. Histograms centered on transcription start site (TSS) or peaks were generated with HOMER annotatePeaks.pl program set in “TSS” mode or with peak files (tss or peaks, hg38, -size 4000, -hist 10). The HOMER annotatePeaks.pl program was used to quantify the read-depth normalized read counts at the merged peaks from all samples for each antibody (made with mergePeaks program). Correlation scatterplots were created with python seaborn. Both HOMER programs automatically normalize each tag directory by the total number of mapped reads. Heatmaps of H3K27ac and H3K27me3 were generated using deepTools (Ramírez et al. 2016) computeMatrix and plotHeatmap with either the TSS or shared peaks. Other plots were made with customized python scripts.

## DATA ACCESS

Data generated for this study are available on NCBI Gene Expression Omnibus (GEO) under accession number GSE304259.

## Supporting information

Cao-etal-revised-2025-biorxiv-automation-files

Cao-etal-revised-2025-biorxiv-Supplemental-Tables-submit

## ACKNOWLEDGEMENTS

Research reported in this publication was supported in part by NIH/NIMH grants R01MH127077 (A.G. and C.B.), R35GM149520 (C.B), R01GM129523 (S.H.), NSF grant 2003358 (A.G.). We thank T. Xu and T. Dishon for their input, and R. Wachs for her help with the illustrations. We thank the Stem Cell Genomics Core at the Sanford Stem Cell Institute and the Epigenomics Core at UC San Diego for providing sequencing services. We also thank N. Gray and B. Nabet (Dana-Farber Cancer Institute) for the HEK293T-BRD4-FKBP12^F36V^-2xHA cell line and for the insightful input.

## Author contributions

A.G. and Y.C. conceptualized and designed the study. Y.C. and L.P. performed the experiments and analyses. Y.C. programmed individual Bravo protocols, and L.A. organized them into a user-friendly interface on VWorks and a distributable package. E.M, C.B, S.H. provided key input in designing, interpreting and analyzing the results. A.G. supervised the study. All authors contributed to writing and reviewing the manuscript.

## Supplemental Material - Table of Contents

**Supplemental Tables (provided as a separate .xlsx file)**

Supplemental Table S1: Experimental materials used for spa-ChIP-seq.

Supplemental Table S2: All antibodies used in this study.

**Supplemental Methods (provided as a separate .pdf file)**

Step-by-step protocol for spa-ChIP-seq.

**Supplemental File (provided as a separate .zip file)**

Required VWorks files for running the Agilent Bravo liquid handlers, which include protocol files, device files, forms, scripts and setup files (labwares, liquid classes, and profiles).

## SUPPLEMENTAL FIGURES

**Supplemental Fig. S1:**
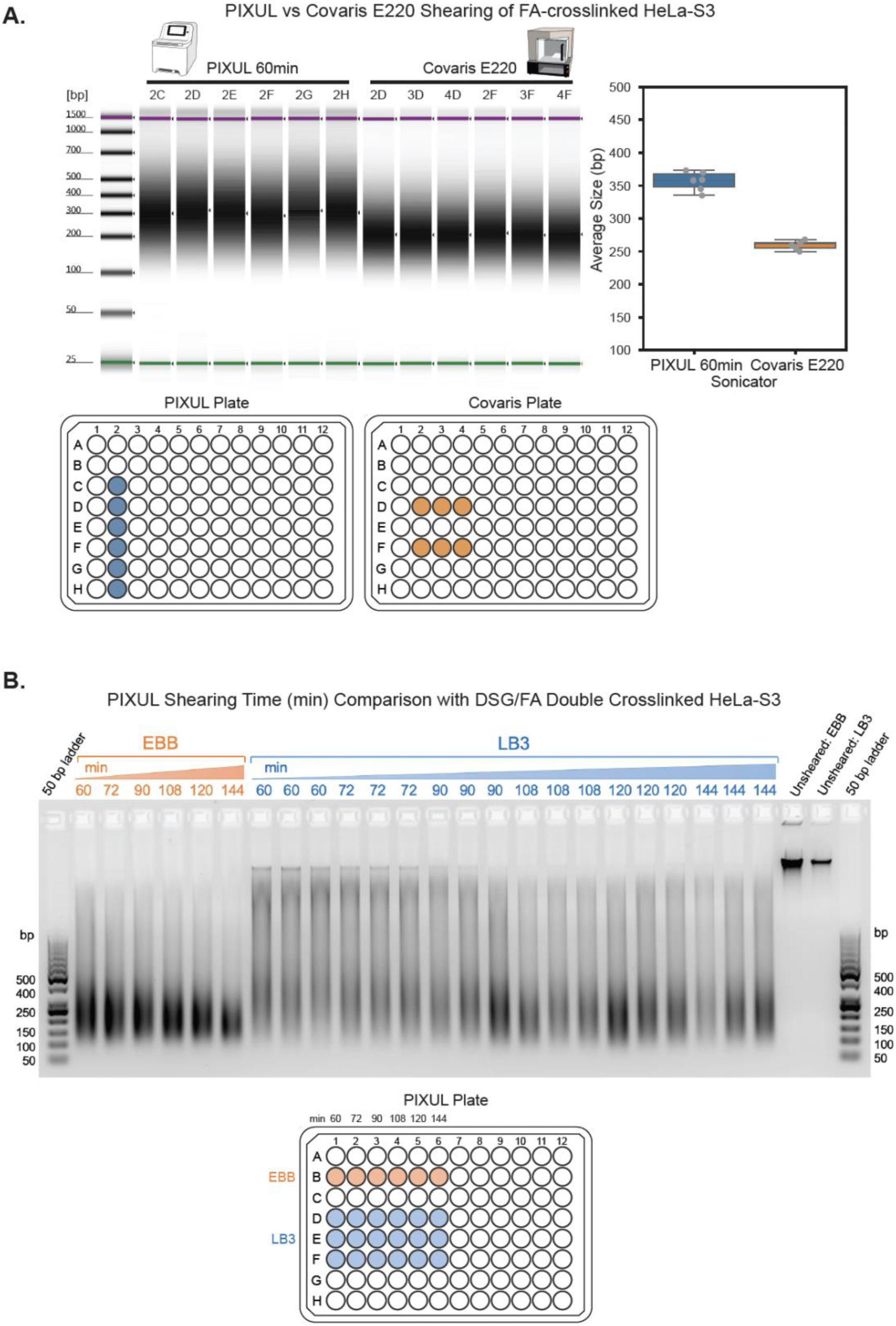
Comparison of 96-well sonicators and lysis buffers for shearing of crosslinked cells. **A.** TapeStation analysis comparing the two 96-well sonicators: PIXUL and Covaris E220 for shearing of FA-crosslinked HeLa-S3. 96-well plate schematics show wells used for the PIXUL (blue) and Covaris (orange) plates during sonication. Boxplot on the right shows the average size (bp) of sheared chromatin on each device based on TapeStation analysis. Error bars represent standard deviations. **B.** 2% agarose gel showing the comparison of sonication time on the PIXUL for lysis buffers EBB and LB3 with DSG/FA double crosslinked HeLa-S3 cells. Schematic of the 96-well PIXUL plate shows wells containing either EBB (orange) or LB3 (blue) lysis buffer during sonication.

**Supplemental Fig. S2:**
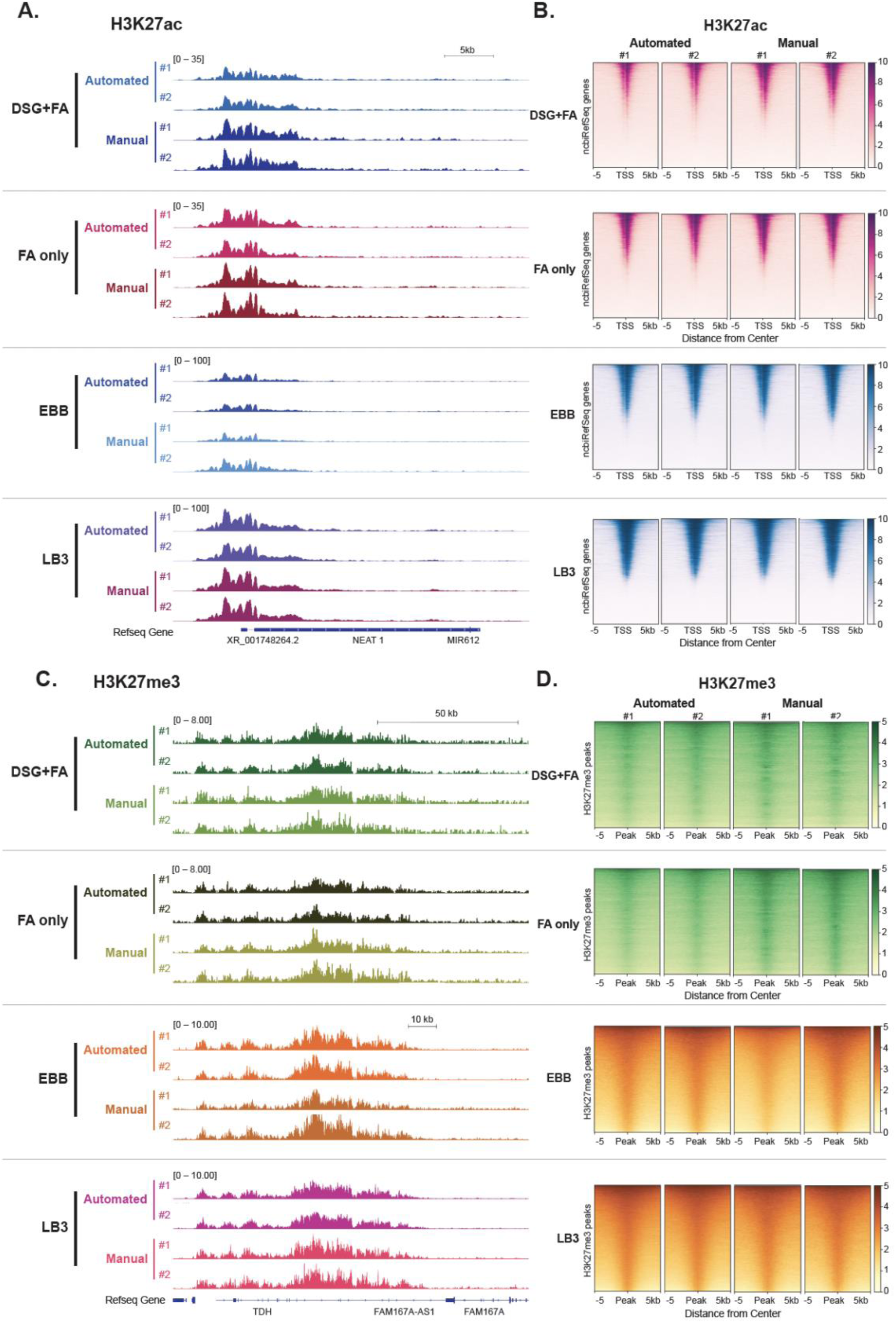
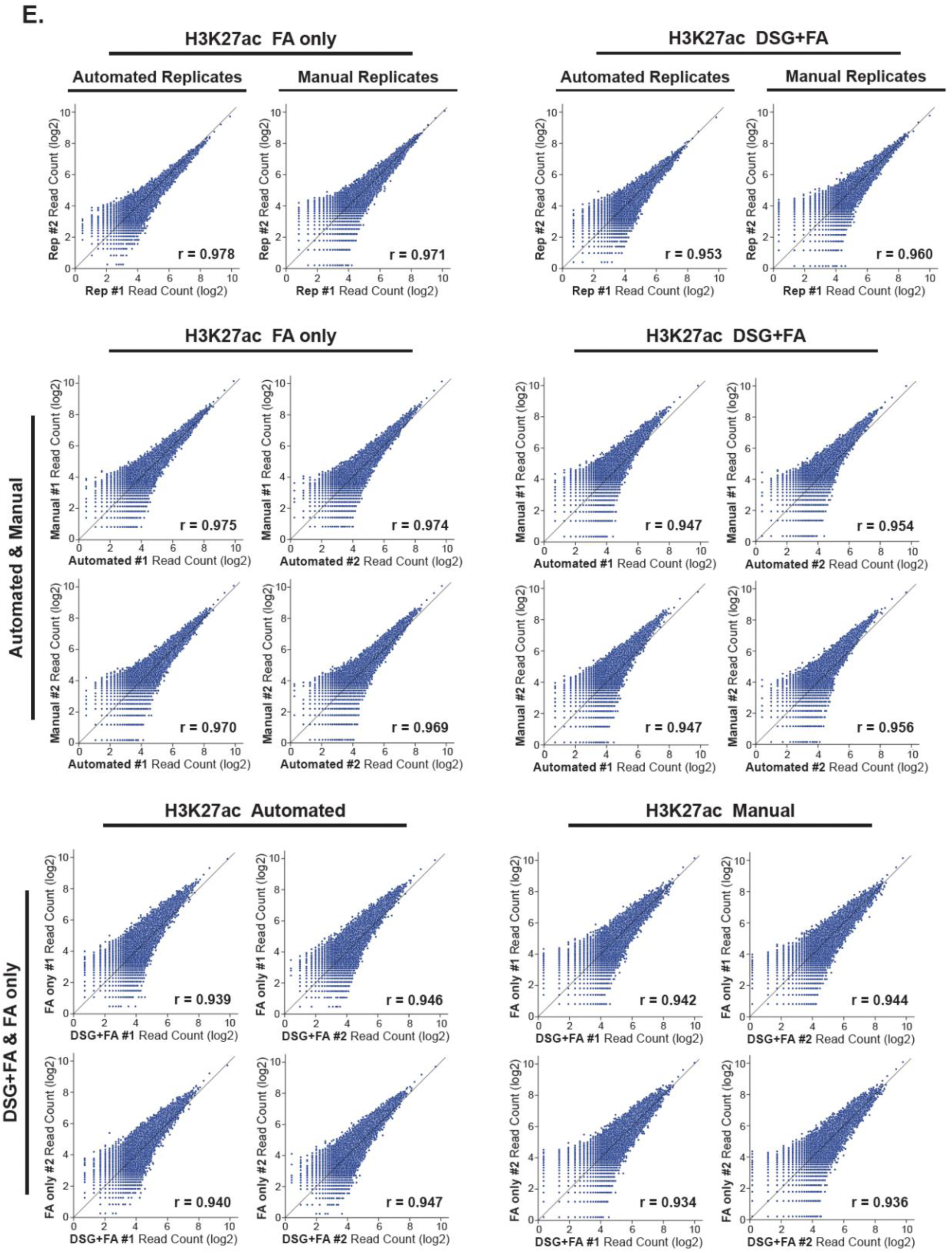
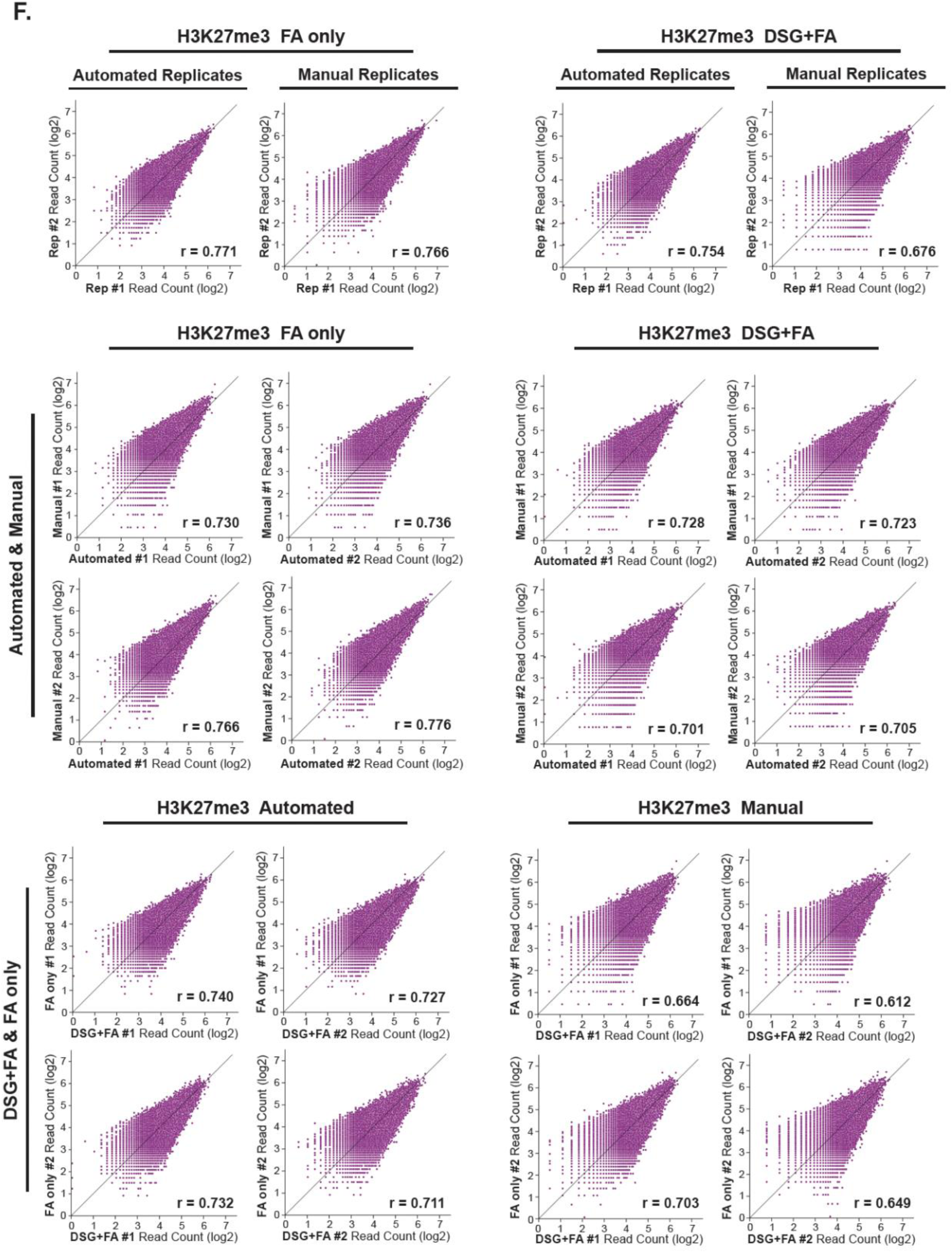
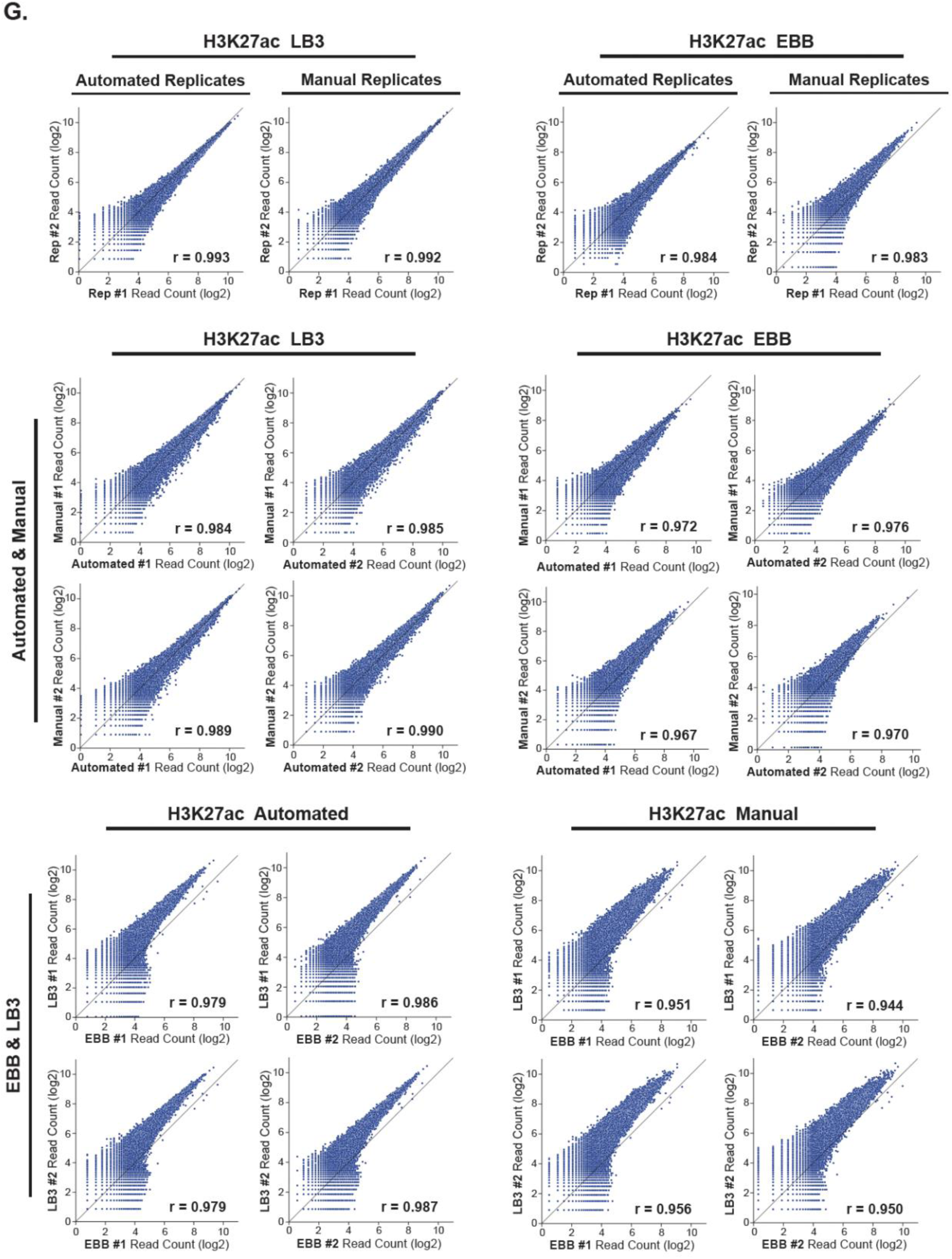
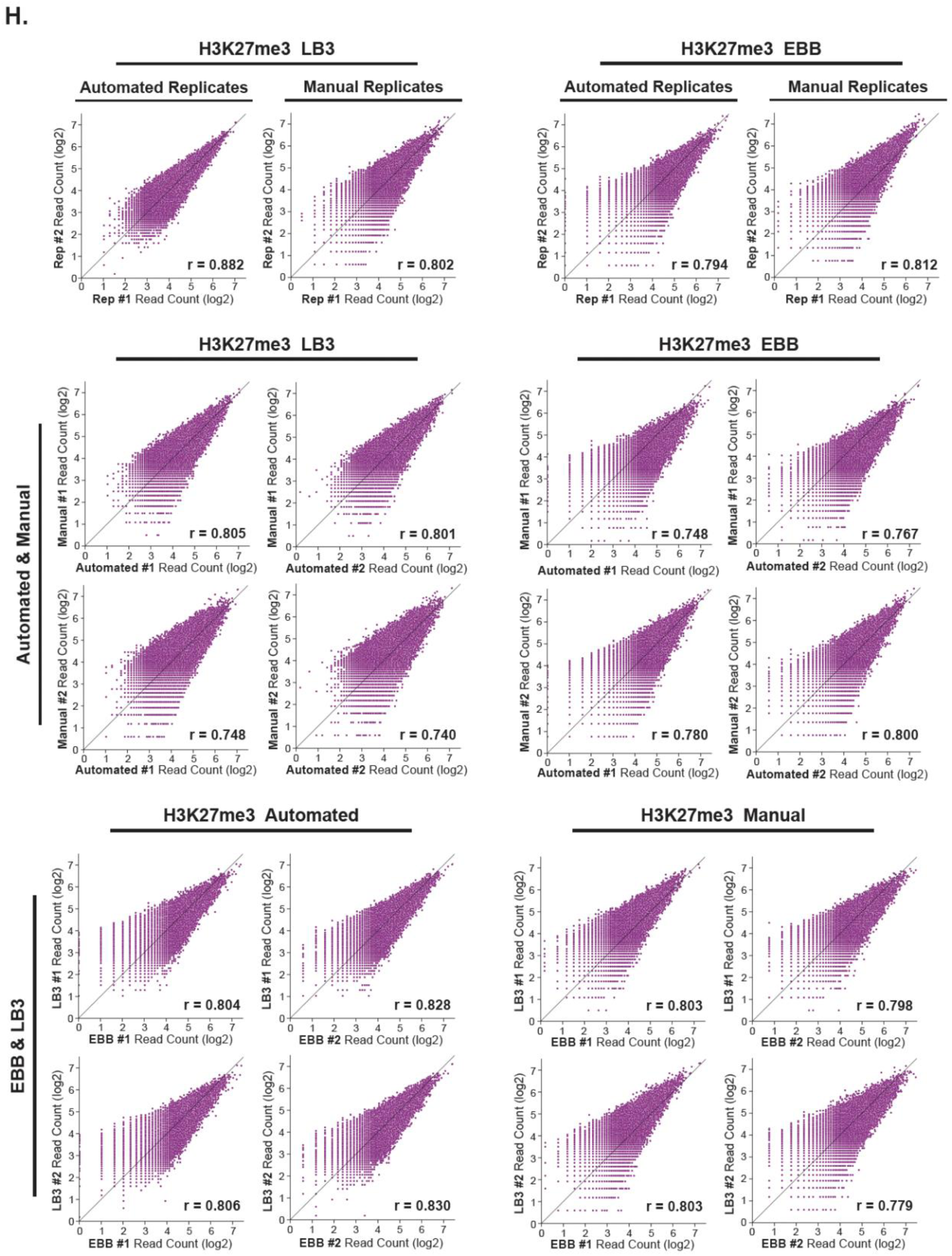
Comprehensive data of our systematic evaluations of spa-ChIP-seq by comparing automated vs. manual workflows, crosslinking conditions and buffer compositions. **A-B.** IGV genome browser tracks (A) and heatmaps of ±5 kb from the center of TSSs (B) showing the automated and manual H3K27ac ChIP-seq results for comparing crosslinking conditions, DSG+FA and FA only, and buffer compositions, EBB and LB3. **C-D.** IGV genome browser tracks (C) and heatmaps of ±5 kb from the center of peaks (D) showing the automated and manual H3K27me3 ChIP-seq results for comparing crosslinking conditions, DSG+FA and FA only, and buffer compositions, EBB and LB3. **E.** Scatterplots of normalized read counts (log2) at shared H3K27ac peaks comparing automated replicates, manual replicates, automated and manual workflows for FA only and DSG+FA ChIP-seq samples, as well as comparing FA only and DSG+FA samples under automated and manual workflows. **F.** Scatterplots of normalized read counts (log2) at shared H3K27me3 peaks comparing automated replicates, manual replicates or automated vs. manual workflows for FA only and DSG+FA ChIP-seq samples, as well as comparing FA only and DSG+FA samples under automated vs. manual workflows. **G.** Scatterplots of normalized read counts (log2) at shared H3K27ac peaks comparing automated replicates, manual replicates, automated and manual workflows for LB3 and EBB ChIP-seq samples, as well as comparing LB3 and EBB samples under automated and manual workflows. **H.** Scatterplots of normalized read counts (log2) at shared H3K27me3 peaks comparing automated replicates, manual replicates, automated and manual workflows for LB3 and EBB ChIP-seq samples, as well as comparing LB3 and EBB samples under automated vs. manual workflows. The Pearson r correlation coefficients are reported on the lower right corner of the scatterplot.

**Supplemental Fig. S3:**
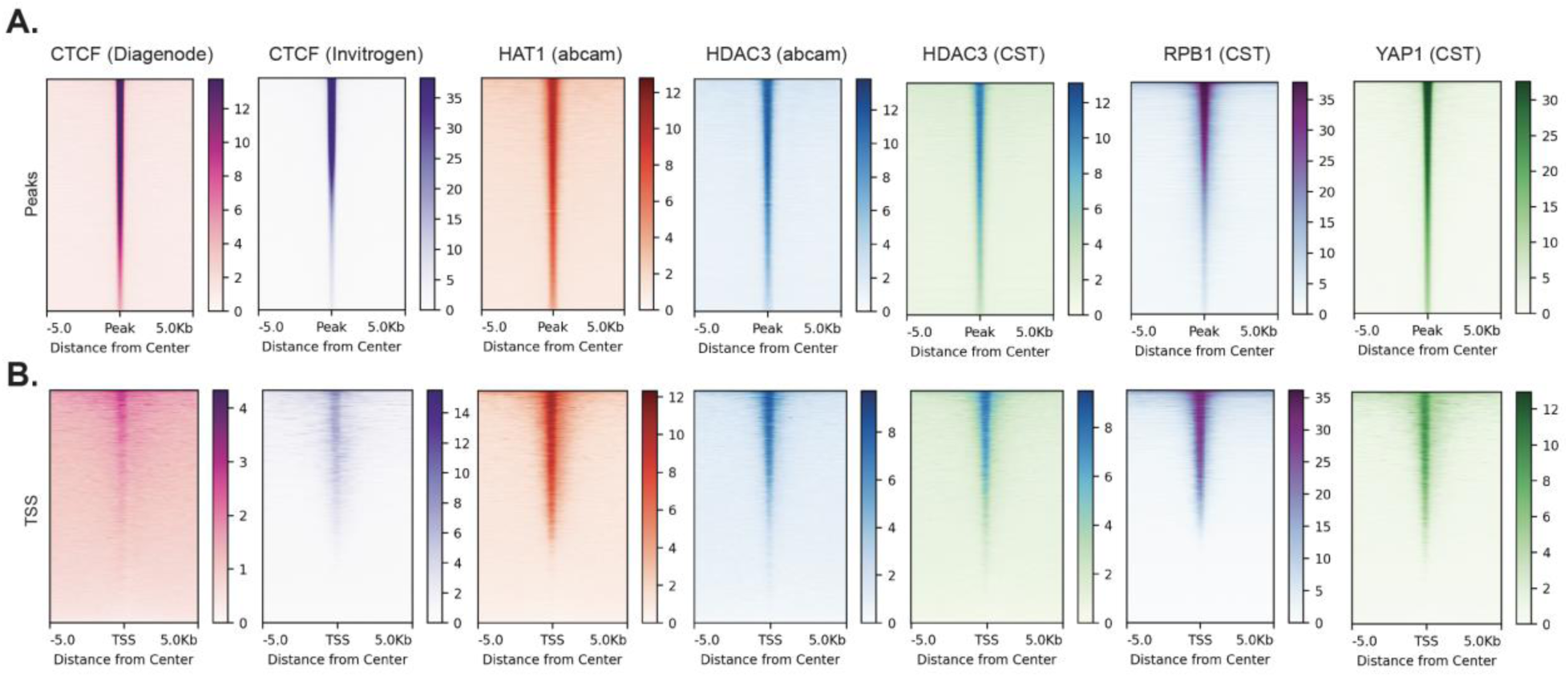
Extended data for mapping of DNA-associated proteins using spa-ChIP-seq. **A-B.** Heatmaps showing average normalized read density of the replicates at ±5 kb from the center of peaks (A) and TSSs (B) for ChIP-seq of DNA-associated proteins: CTCF, HAT1, HDAC3, Rpb1, and YAP1.

**Supplemental Fig. S4:**
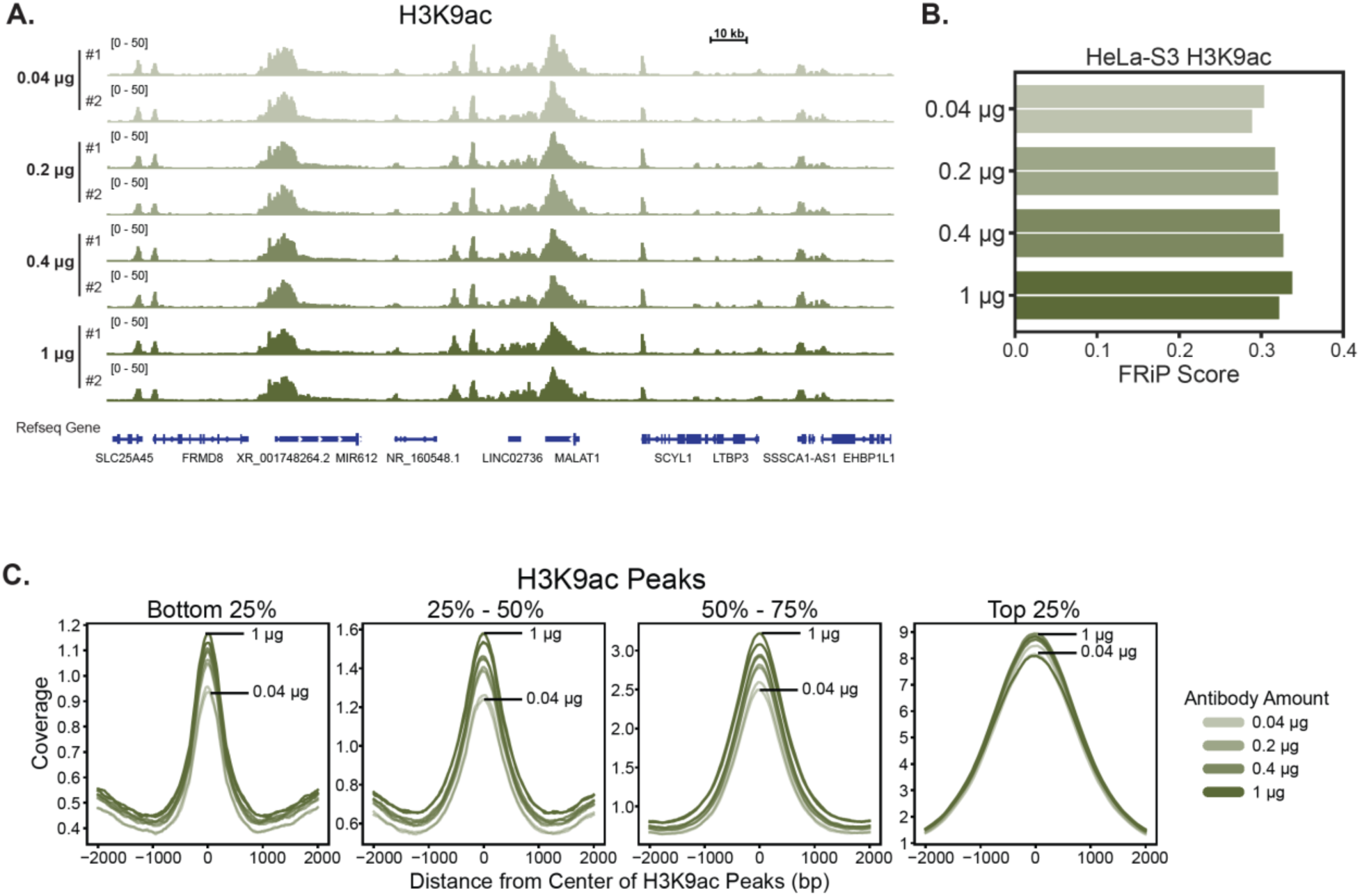
Extended data for H3K9ac antibody titration ChIP-seq experiments. **A.** IGV genome browser tracks for H3K9ac ChIP-seq done with various amounts of antibody (0.04 μg, 0.2 μg, 0.4 μg, 1 μg), each in replicates. **B.** Bar plot of FRiP scores for ChIP-seq of H3K9ac antibody titrations shown in (A). **C.** Read coverage histograms for H3K9ac antibody titration ChIP-seq signal centered on H3K9ac peaks, separated into quartiles based on the normalized read counts at the peaks.

**Supplemental Fig. S5:**
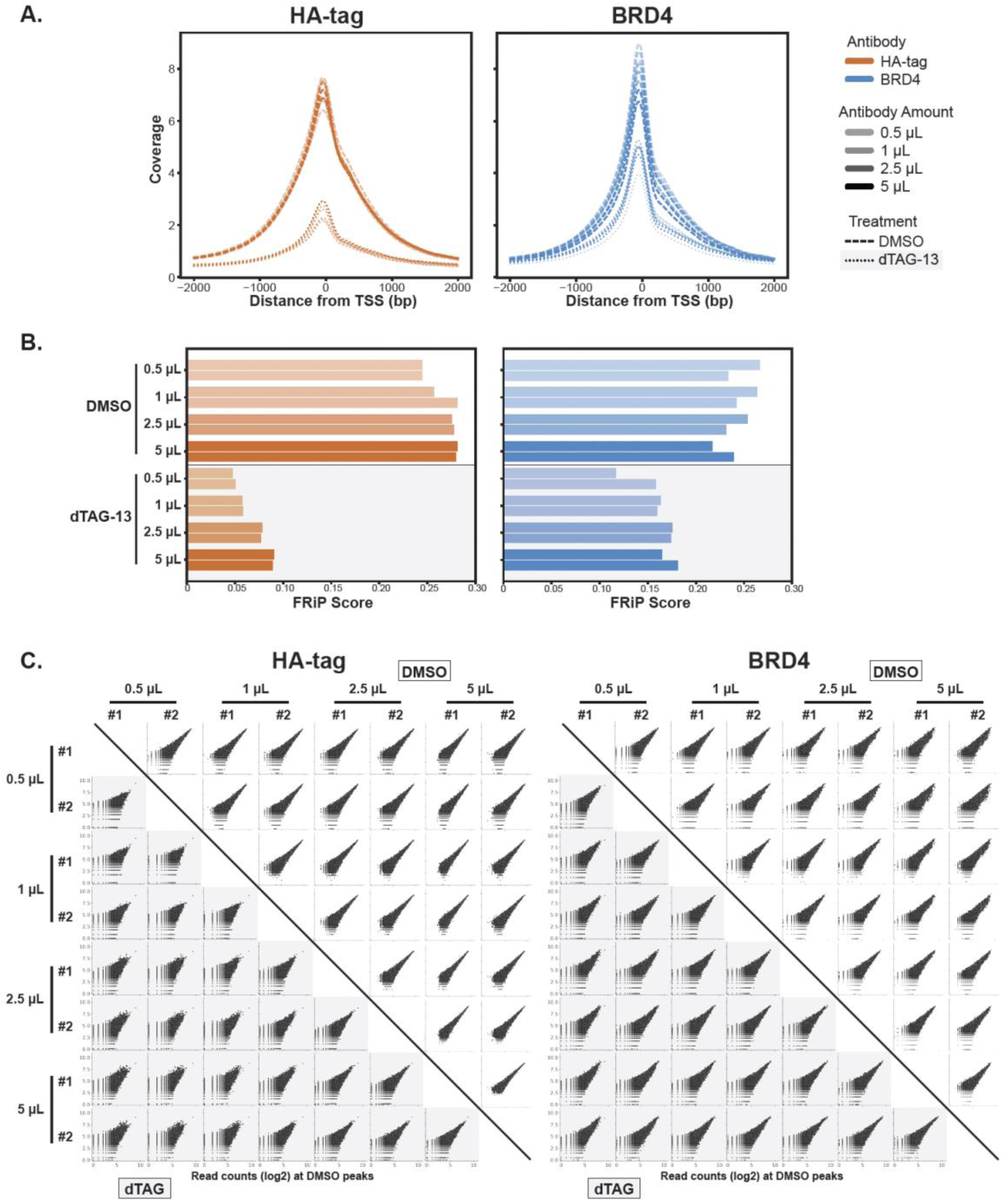
Extended data for HA-tag and BRD4 antibody titration ChIP-seq experiments. **A.** Metagene plot of ±2000 bp from the center of TSS for BRD4 and HA-tag ChIP-seq in dTAG-13 and DMSO treated HEK293T-BRD4-FKBP12^F36V^-2xHA cells with 0.5 μL, 1 μL, 2.5 μL, and 5 μL of HA-tag and BRD4 antibodies. **B.** Bar plot of FRiP scores for ChIP-seq of HA-tag and BRD4 antibody titrations shown in (A). **C.** Scatterplots showing normalized read counts (log2) at DMSO peaks for HA-tag and BRD4 antibody titration. The DMSO samples are in the upper right corner and dTAG samples are in the lower left corner shaded with a light gray background.

**Supplemental Fig. S6:**
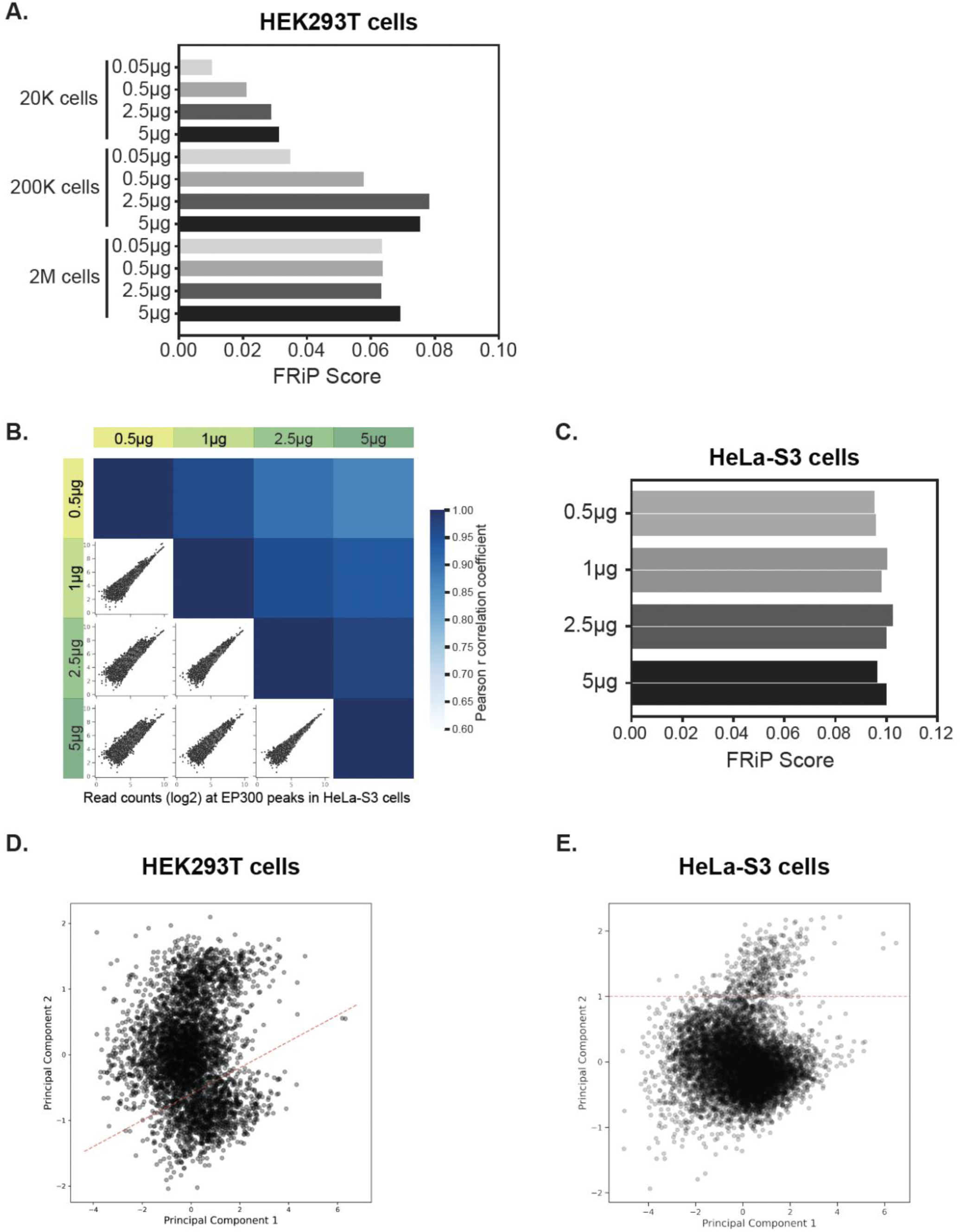
Extended data for EP300 antibody titration ChIP-seq experiments in HeLa-S3 and HEK293T cells. **A.** Bar plot of FRiP scores using the peaks called in each sample for ChIP-seq of EP300 antibody titrations with 0.05 μg, 0.5 ug, 2.5 ug, and 5 ug of antibody and 20K, 200K, and 2M of HEK293T cells. **B.** A matrix of scatterplots showing normalized read counts (log2) at EP300 peaks and heatmaps showing Pearson r correlation coefficients for EP300 antibody titrations in 1M HeLa-S3 cells with various antibody amounts 0.05 μg, 0.5 μg, 2.5 μg, and 5 μg of EP300 antibody (green shades). **C.** Bar plot of FRiP scores using the merged peaks of the replicates for each condition for ChIP-seq of EP300 antibody titrations with 0.05 μg, 0.5 μg, 2.5 μg, and 5 μg of EP300 and 1M of HeLa-S3 cells. Each condition is done in replicates represented as two bars with the same shading. **C-D.** Principal component analysis (PCA) on the normalized read counts (log2) at the top 10% of EP300 ChIP-seq peaks for 0.5 μg and 5 μg of antibody in 2M HEK293T cells (C) and 1M HeLa-S3 cells (D). The red dashed line shows how the red subset of peaks in **Fig. 4C** is isolated for each of the samples (the region under the line in d and the region above the line in e).

**Supplemental Fig. S7:**
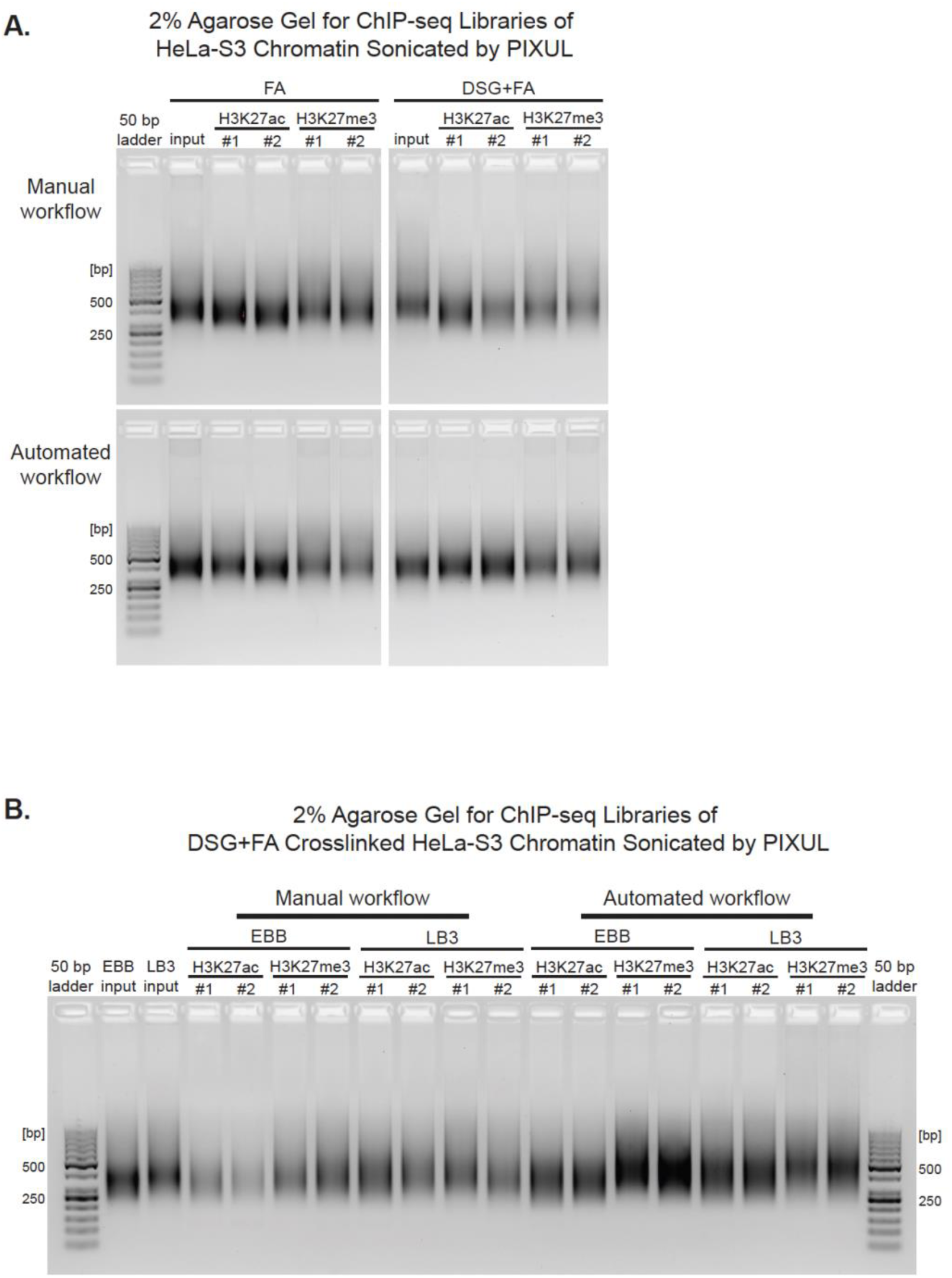
Size estimation of DNA libraries from automated and manual ChIP-seq with agarose gel. **A-B.** 2% agarose gel showing DNA libraries generated from manual and automated H3K27ac and H3K27me3 ChIP-seq. (A) Libraries prepared from HeLa-S3 chromatin crosslinked with either FA or DSG+FA and sonicated with LB3 lysis buffer on PIXUL. (B) Libraries prepared from DSG+FA crosslinked HeLa-S3 chromatin lysed in either EBB or LB3 buffer and sonicated using PIXUL. The DNA libraries showed a smear with high intensity around 250–500 bp on the gel, consistent with the expected fragment size distribution.

## spa-ChIP-seq with Bravo

spa-ChIP-seq protocol is based on the following published protocol:

Texari L, Spann NJ, Troutman TD, Sakai M, Seidman JS, Heinz S. 2021. An optimized protocol for rapid, sensitive and robust on-bead ChIP-seq from primary cells. *STAR Protocols* 2: 100358. https://doi.org/10.1016/j.xpro.2021.100358.

### Preparing buffers/solutions

- 10% BSA in PBS: Mix 5 grams of BSA with 50 mL of PBS at room temperature.
- 2.625 M Glycine: Mix 9.85 g glycine with 50 mL of UltraPure DNase/RNase-Free Distilled at room temperature.
- Lysis buffer LB3: Freshly prepare before sonication. Table below shows an example of the amount of each reagent required for making 10 mL of LB3. 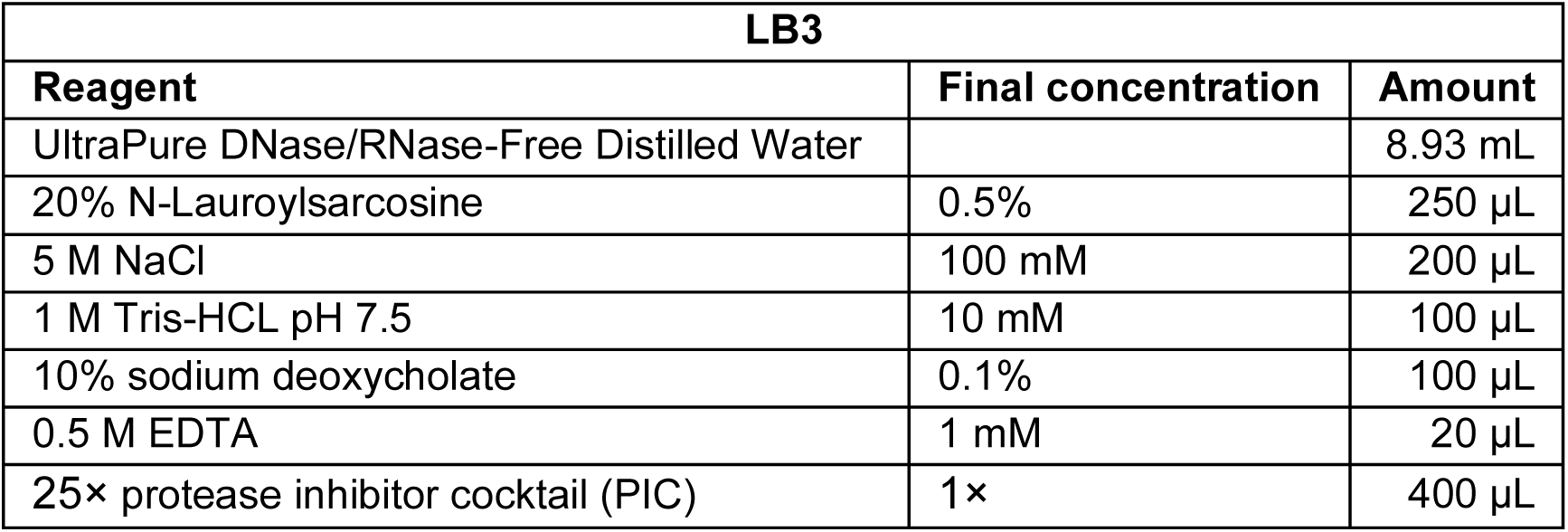
- 10% Triton X-100: Mix 5 mL of Triton X-100 with 45 mL of UltraPure water to prepare a total of 50 mL.
- ChIP wash buffer 1 (WBI): Make a stock of 500 mL according to the table below. Store at 4°C. 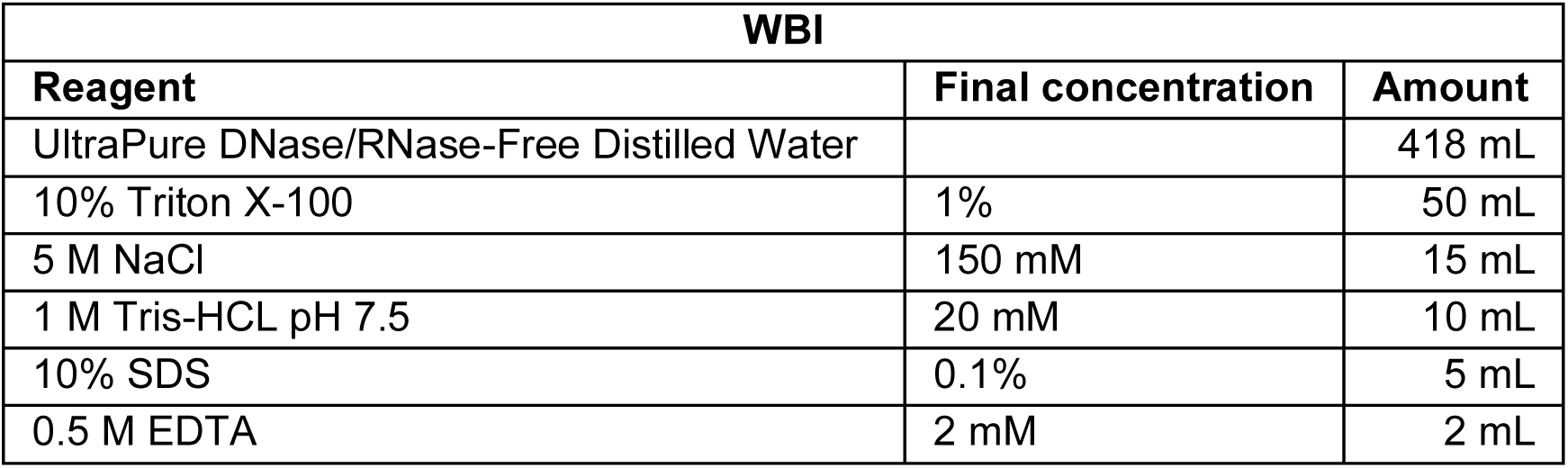
- ChIP wash buffer 3 (WBIII): Make a stock of 500 mL according to the table below. Store at 4°C. 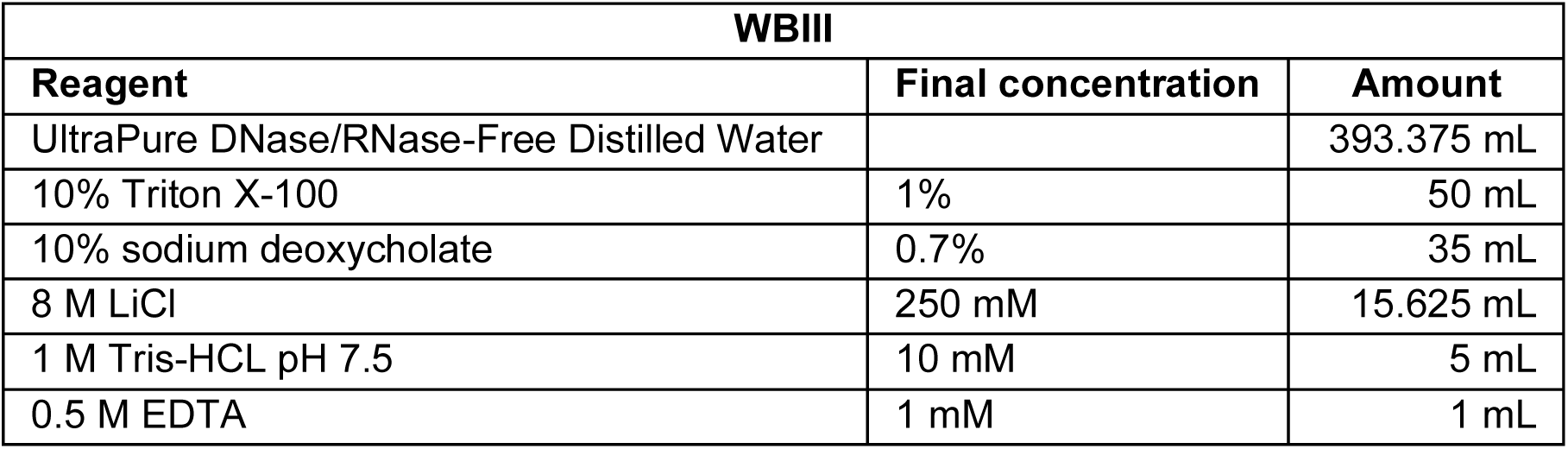
- TET buffer: Make a stock of 500 mL according to the table below. Store at 4°C. 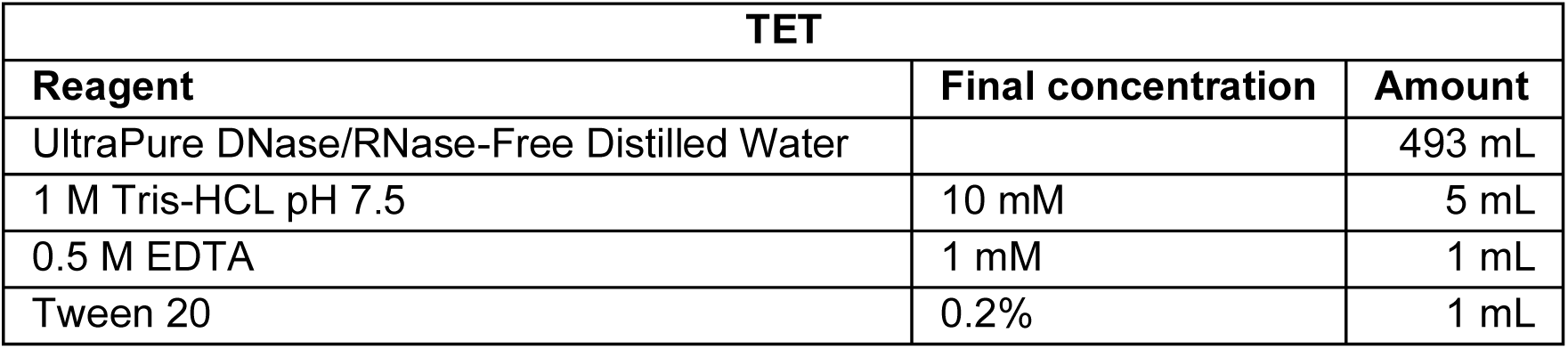
- TT buffer: Make a stock of 50 mL according to the table below. Store at 4°C. 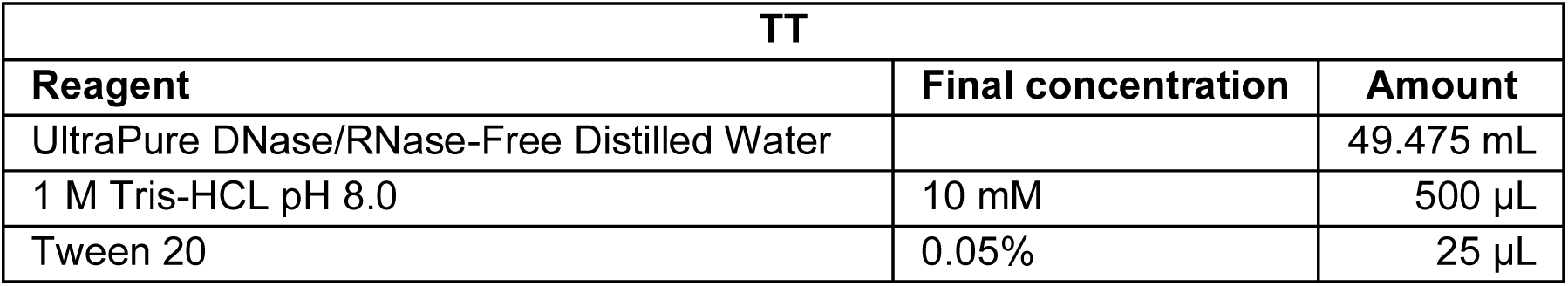

### Crosslink cells

1. Grow cells till ∼80%-100% confluency. Count cell numbers before harvest if necessary.
2. Prepare crosslinking reagents for either double or single crosslinking.

a. **Double crosslinking (DSG+FA)**: For each **1 million cells**, freshly prepare a minimum of **1 mL of a 2 mM DSG solution in PBS**. First warm up the glass vial of 100 mg DSG to room temperature before opening (to avoid water condensation into the vial). Each 1 million cells require at least 0.654 mg DSG dissolved in 3.92 µL anhydrous DMSO (adjust the amount of DMSO to the amount of DSG in the tube). Place 1 mL PBS per 1 million cells in a conical tube. Using a micropipette, swiftly ‘‘shoot’’ **3.92 µL of the DSG/DMSO solution** into the PBS, immediately close the tube, and invert multiple times to rapidly disperse the DSG. If this is done too slowly, the DSG will form a white precipitate, and the DSG/PBS preparation procedure will need to be repeated.
b. **Single crosslinking**: For each **1 million cells**, freshly prepare a minimum of **1 mL of a 1% formaldehyde solution in PBS**. Place 1 mL PBS per 1 million cells in a conical tube and add 66.67 µL of 16% formaldehyde solution per 1 mL PBS.
3. Prepare cells for crosslinking either on tissue culture plates or in conical tubes.

a. **Crosslink adherent cells on plate**: Aspirate old media from the plate. Wash cells once with 5-10 mL PBS.
b. **Crosslink adherent cells in tube**: Harvest adherent cells by trypsinizing and neutralize trypsin with 4 volumes of FBS-containing media, transfer cell suspension to 15 mL or 50 mL conical, depending on volume. Pellet cells by centrifuging for 8 min at 300 × g at 20°C–25°C. Aspirate or decant supernatant. Wash cells once by resuspending in 5-10 mL PBS and centrifugation for 8 min at 300 × g at 20°C–25°C.
c. **Crosslink suspension cells in tube**: Transfer cell suspension to 15 mL or 50 mL conical, depending on volume. Pellet cells by centrifuging for 8 min at 300 × g at 20°C– 25°C. Aspirate or decant supernatant. Wash cells once by resuspending in 5-10 mL PBS and centrifugation for 8 min at 300 × g at 20°C–25°C.
4. Double crosslinking with DSG and FA:

a. Make sure the PBS wash is aspirated.
b. Per 1 million cells, add at least 1 mL of freshly made 2mM DSG/PBS solution, and incubate at room temperature for 30 minutes while shaking the plates on a platform shaker or rotating the tubes overhead at 8 rpm.
c. Next, per 1 million cells, add 66.67 µL of 16% formaldehyde to each 1 ml cell suspension in DSG/PBS, and incubate at room temperature for an additional 10 minutes while shaking the plates on a platform shaker or rotating the tubes overhead at 8 rpm.
5. Single crosslinking with FA only:

a. Make sure the PBS wash is aspirated.
b. Per 1 million cells, add at least 1 mL of freshly made 1% formaldehyde/PBS solution, and incubate at room temperature for 10 minutes while shaking the plates on a platform shaker or rotating the tubes overhead at 8 rpm.
6. Quench the crosslinking reaction by adding 1/20th volume of 2.625 M Glycine and 1/20^th^ volume 10% BSA in PBS (50 μL each per each 1 mL crosslinking reaction).
7. If crosslinking on plate, scrape the cells and transfer the cell slurry into a conical tube. Pellet cells by centrifuging for 5 minutes at 1000–1500 × g at 4°C.
8. Aspirate and discard supernatant, then resuspend fixed cells in 1 mL ice-cold 0.5% BSA/PBS and transfer the cell suspension to a 1.5 mL microcentrifuge tube.
9. Pellet cells by centrifuging for 5 minutes at 1000–1500 × g at 4°C.
10. Aspirate and discard supernatant and resuspend fixed cells in 1 mL ice-cold 0.5% BSA/PBS.
11. Pellet cells by centrifuging for 5 minutes at 1000–1500 × g at 4°C.
12. Aspirate and discard supernatant (can leave behind ∼10 µL).
13. If proceeding to next steps immediately, place the cell pellet on ice. If saving the cells for future use, snap-freeze the cell pellet and store at −80°C.

### Shearing Crosslinked Cells with PIXUL™ Multi-Sample Sonicator

1. Per 1 million fixed cells, freshly prepare at least 100 µL of lysis buffer LB3 with protease inhibitor cocktail (PIC) added. Mix by vortexing and keep on ice.
2. Add ∼100 µL of lysis buffer LB3+PIC into 1 million fixed cell pellet and suspend the mixture thoroughly.
3. Load 100 µL of cells in lysis buffer into each well of the PIXUL shearing plate (∼10,000 - 5,000,000 cells per well) and seal the plate with a PCR plate pressure seal.

- **Note**: All wells lacking sample in the columns being sonicated MUST be filled with liquid (water, coupling buffer, etc.) prior to starting the sonication run. Be sure to keep the outside bottom of the plate clean and free of lint or other debris.
4. Turn on the power switch on the back side of the PIXUL instrument to the ON position. The light around the center power button on the front of the instrument should illuminate.
5. Press the main power button on the front side of the PIXUL instrument. The touchscreen will start initializing and load the home screen.
6. Once PIXUL is on, the first step is to ensure that the PIXUL Coupling Fluid level is about an inch (2.5 cm) below the reservoir top (or just above the circular cut-out in the plastic column within reservoir). The reservoir capacity is approximately 600 mL, and the fluid level should be topped off as needed. Do not overfill.
7. Lift the white external lid of the PIXUL instrument and place the shearing plate on the black grid with well A1 in the upper left corner, secure down the black hold-down pressure distribution lid on top of the plate with plate-securing tension rods/clamps (fastening both sides at the same time), and close the white external lid.
8. On the touchscreen, press “CIRCULATE” to initiate PIXUL Coupling Fluid cooling and allow PIXUL to cool to at least 15°C, which takes up to 5 minutes. You cannot start the sonication run without circulation active. You can monitor Coupling Fluid temperature in the upper right-hand corner of the touchscreen.
9. On the touchscreen, select the plate columns for which you would like to set sonication parameters. Columns selected together will be outlined in the same color.
10. On the left side of the touchscreen, you can use the left and right arrows to select from saved presets of sonication parameters. You can also use the add and delete buttons to add or remove a row of process settings. Adjust the following parameters: 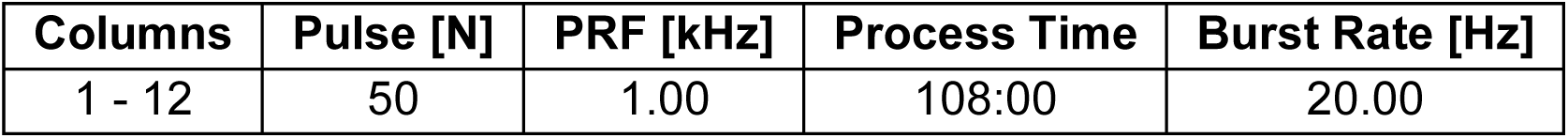

- **Note**: The software will not allow settings that could damage the transducers, and the upper limits are a maximum Pulse of 50 or a maximum PRF of 1.00. Recommended a minimum Burst Rate is 20.00. Varying the Process Time is typically the only parameter that needs adjusting.
11. Once the PIXUL Coupling Fluid has reached approximately 15°C, press “START” on the touchscreen. The run-time to completion will appear in the top left-hand corner of the touchscreen. Do not lift lid while PIXUL runs.
12. Once the sonication run has completed, pop-up appears on screen, click OK, open the external lid. PIXUL Coupling Fluid will drain from underneath the sample plate for the next few seconds (this will happen automatically and will take 5-10 seconds).
13. Unload the sample plate by lifting the pressure distribution plate cover by the lift handle and simply pulling up. Close lid but do not fasten the tension rods.
14. To turn off the PIXUL instrument, press the main power button on the front side of the PIXUL instrument and switch the power switch off on the back side of the PIXUL instrument.
15. After sonication, spin down the plate for at least 1 minute at 1000–1500 × g.

### Analysis of Sonicated Chromatin

1. Freshly prepare the following **Reaction Mix** at room temperature. Each sample requires 29 μL and it is recommended to prepare 20% more reaction volumes per sample. Aliquot 1/8 of the total volume of the mastermix to each well in the first column of the **Reaction Mix Plate**. Place the **Reaction Mix Plate** on **Bravo deck 6**. (The example below shows the amount required for a full 96-well plate.) 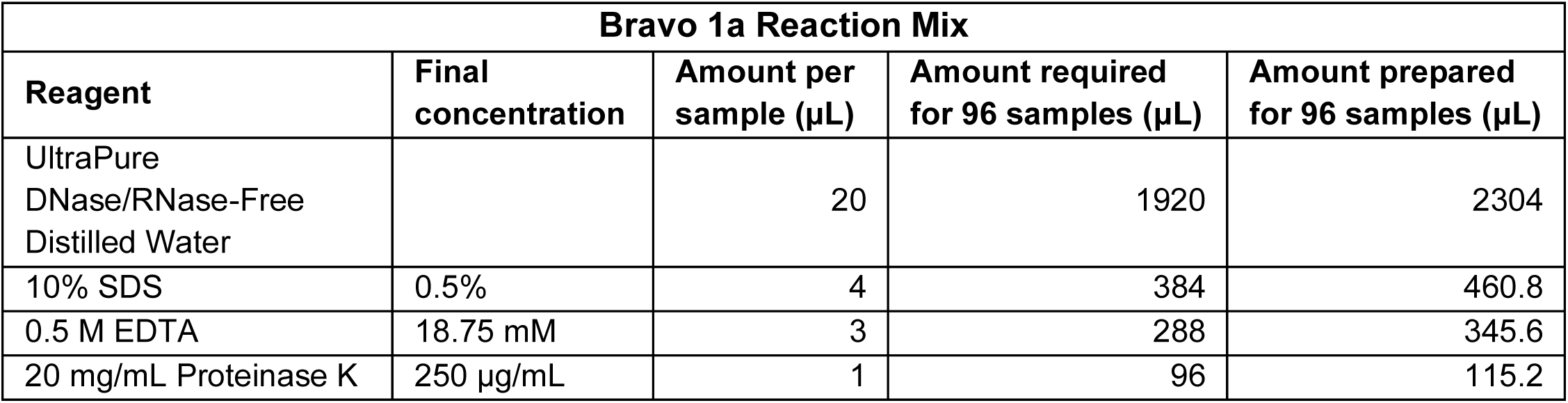
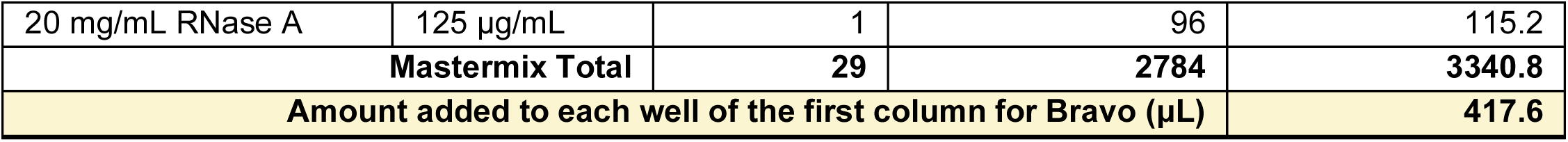
2. Place the **PIXUL Shearing Plate** on **Bravo deck 5** and **5 M NaCl Reservoir** (filled with ∼5-10 mL 5 M NaCl) on **Bravo deck 9**. Ensure the reservoirs contain at least enough liquid to form a thin layer across the bottom.
3. Select Bravo protocol **1a_Input_Prep_vA1.0.1.pro** from the drop down manual in step 1 and click the green check symbol to display the Bravo Deck Setup. Make sure the deck is set up as shown. Once ready, click on the right-pointing triangle (▷) at step 7 to start the protocol. 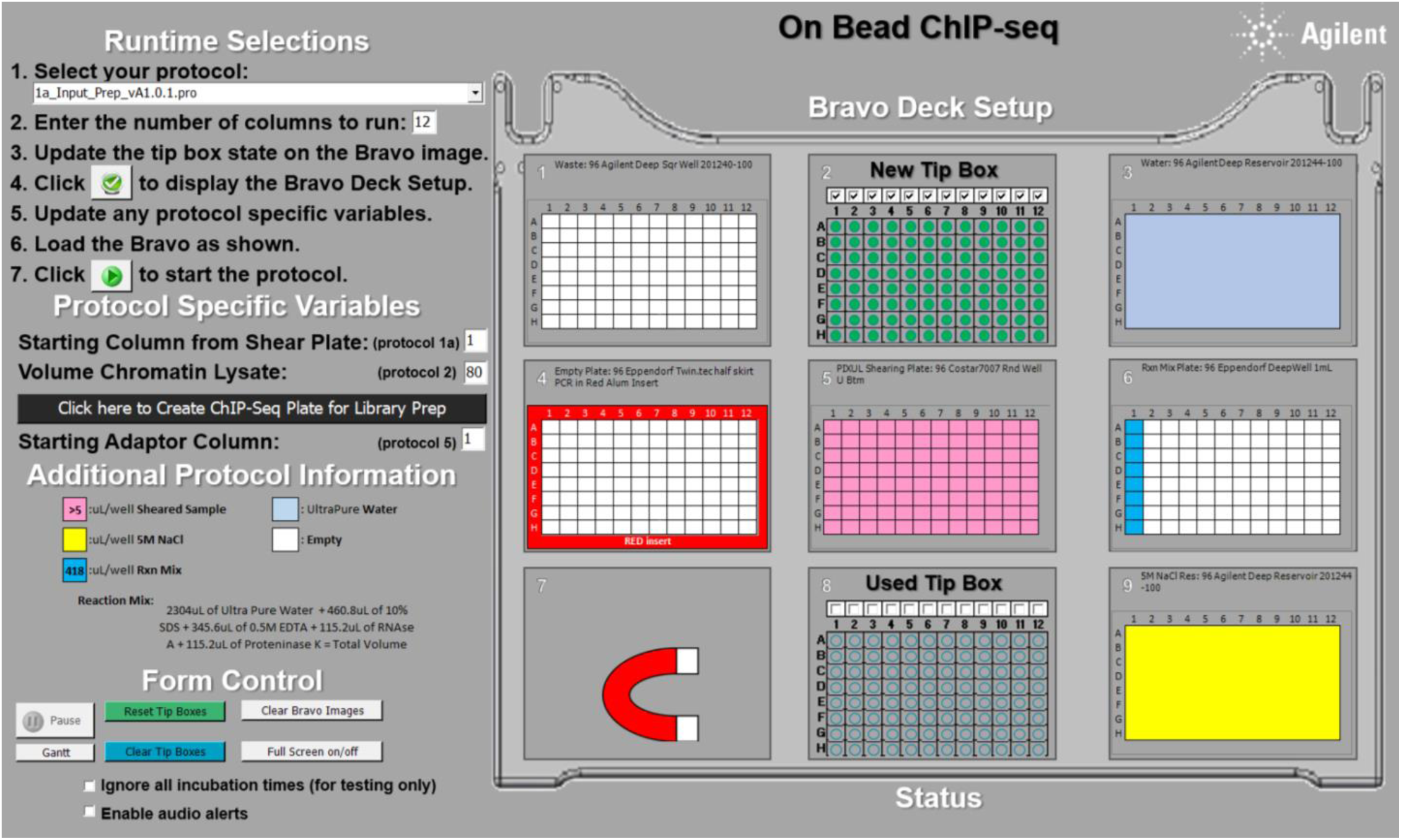 Bravo will perform the following steps:

- Combine 5 μL sonicated chromatin lysate from each well and 41.5 μL of UltraPure water to a final volume of 46.5 μL.
- Add 29 μL of the Reaction Mix to each sample.
- Add 4.5 μL of 5 M NaCl to each sample.
- Mix up and down at least 10 times (the mix cycle can be changed in the parameters).
4. **Reverse crosslinking**: Retrieve the plate with samples from **Bravo deck 4**. Incubate the plate at 37°C for 15 minutes and at 55°C for 30 minutes to digest RNA and proteins, then at 65°C for 30 minutes to 1 hour in a PCR cycler with the heated lid set to at least 75°C to prevent evaporation and drying out of the samples. This can be left at 12°C or 4°C overnight.
5. Freshly prepare the following **Speedbeads Mastermix** at room temperature. Each sample requires 80 μL and it is recommended to prepare 20% more reaction volumes per sample. Aliquot 1/8 of the total volume of the mastermix to each well in the first column of the **Speedbeads Mastermix Plate**. Place the **Speedbeads Mastermix Plate** on **Bravo deck 5**. (The example below shows the amount required for a full 96-well plate.) 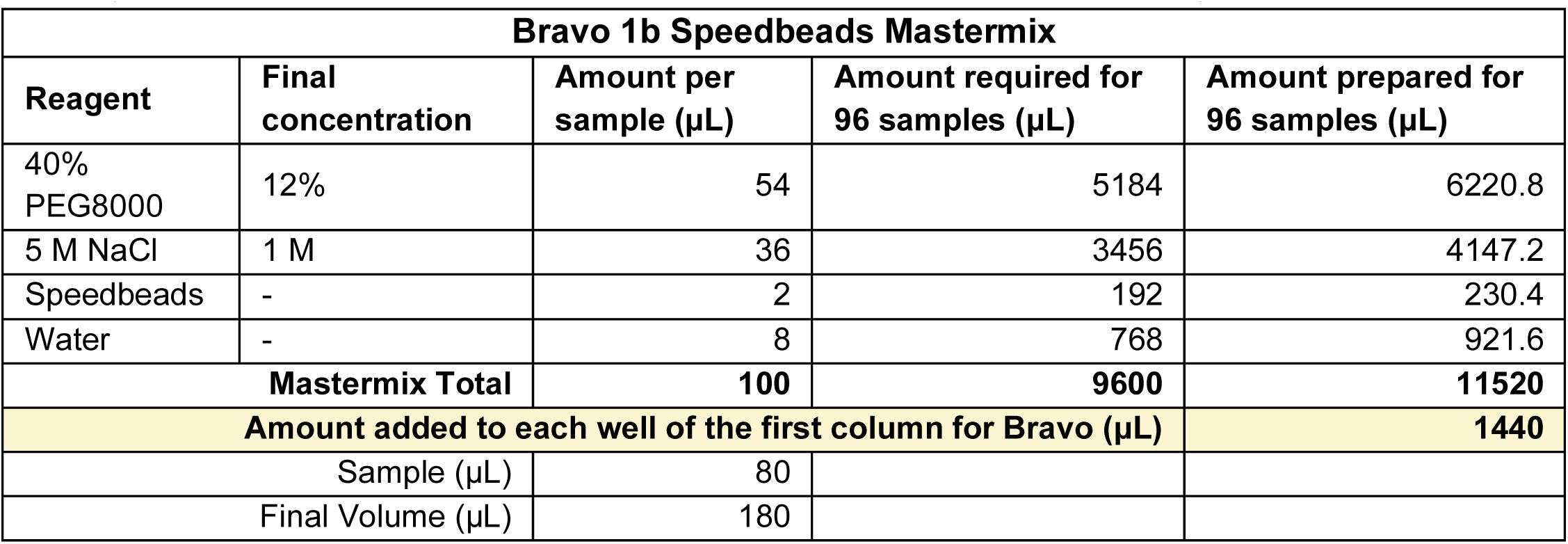
6. Remove the **Sample Plate** from reverse crosslinking incubation and place it back to **Bravo deck 4**.
7. Select Bravo protocol **1b_Input_PEG_vA1.0.1.pro** from the drop down manual in step 1 and click the green check symbol to display the Bravo Deck Setup. Make sure the deck is set up as shown. Once ready, click on the right-pointing triangle (▷) at step 7 to start the protocol. 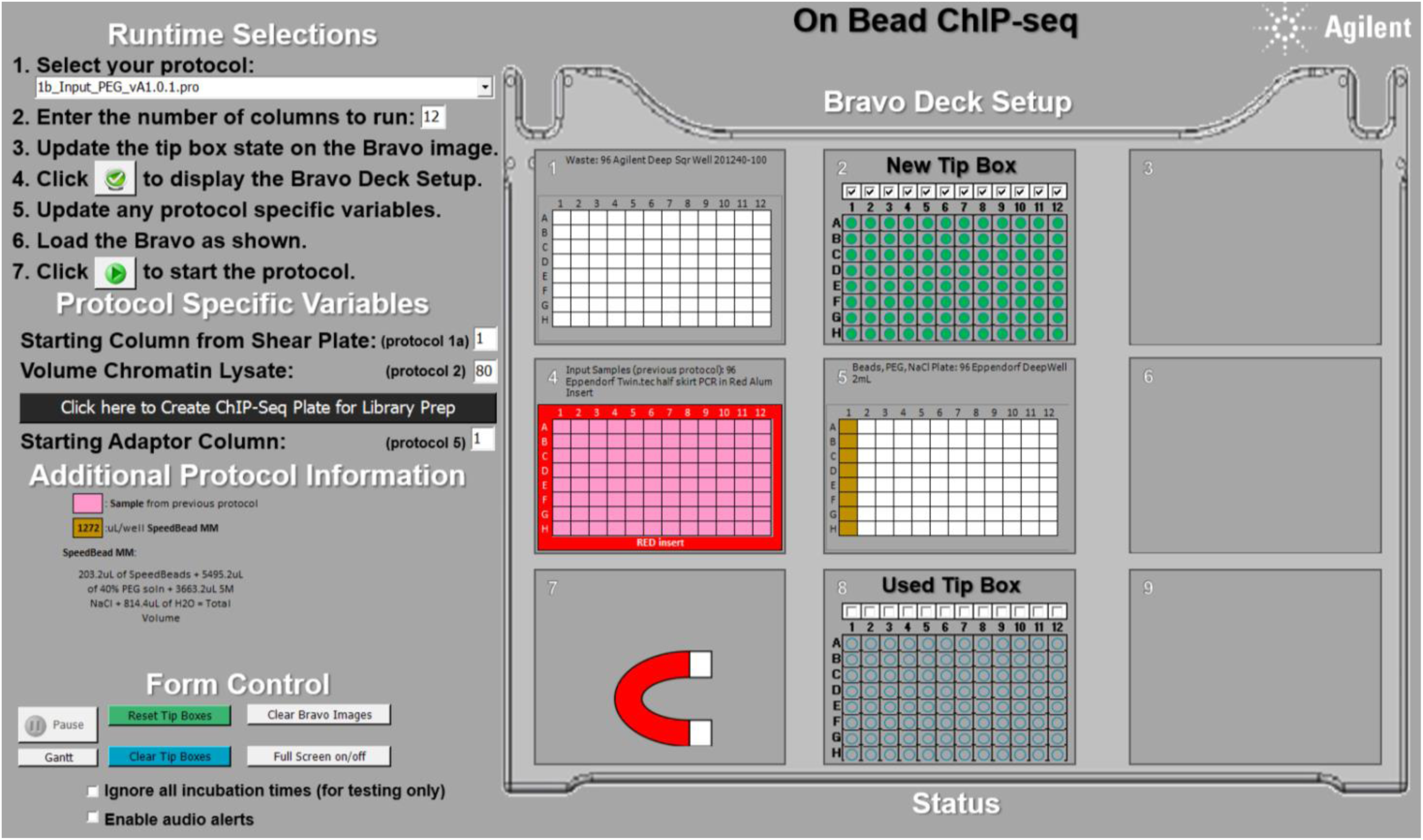 Bravo will perform the following steps:

- Add 100 μL of SpeedBead/PEG mastermix to each 80 μL ChIP input sample. The final concentration of PEG will be 12% and the final concentration of NaCl will be 1 M.
- Mix the samples to homogeneity by repetitive pipetting. Incubate at room temperature for 10 minutes.
8. During incubation, place the **Waste Plate** on **Bravo deck 1**, **TT Buffer Reservoir** on **Bravo deck 3**, **empty Elution Plate** on **Bravo deck 6**, and **80% Ethanol Reservoir** on **Bravo deck**
9. Ensure the reservoirs contain at least enough liquid to form a thin layer across the bottom.
10. Immediately after protocol 1b is completed, select Bravo protocol **1c_Input_Speedbead_Cleanup_vA1.0.1.pro** from the drop down manual in step 1 and click the green check symbol to display the Bravo Deck Setup. Make sure the deck is set up as shown. Once ready, click on the right-pointing triangle (▷) at step 7 to start the protocol. 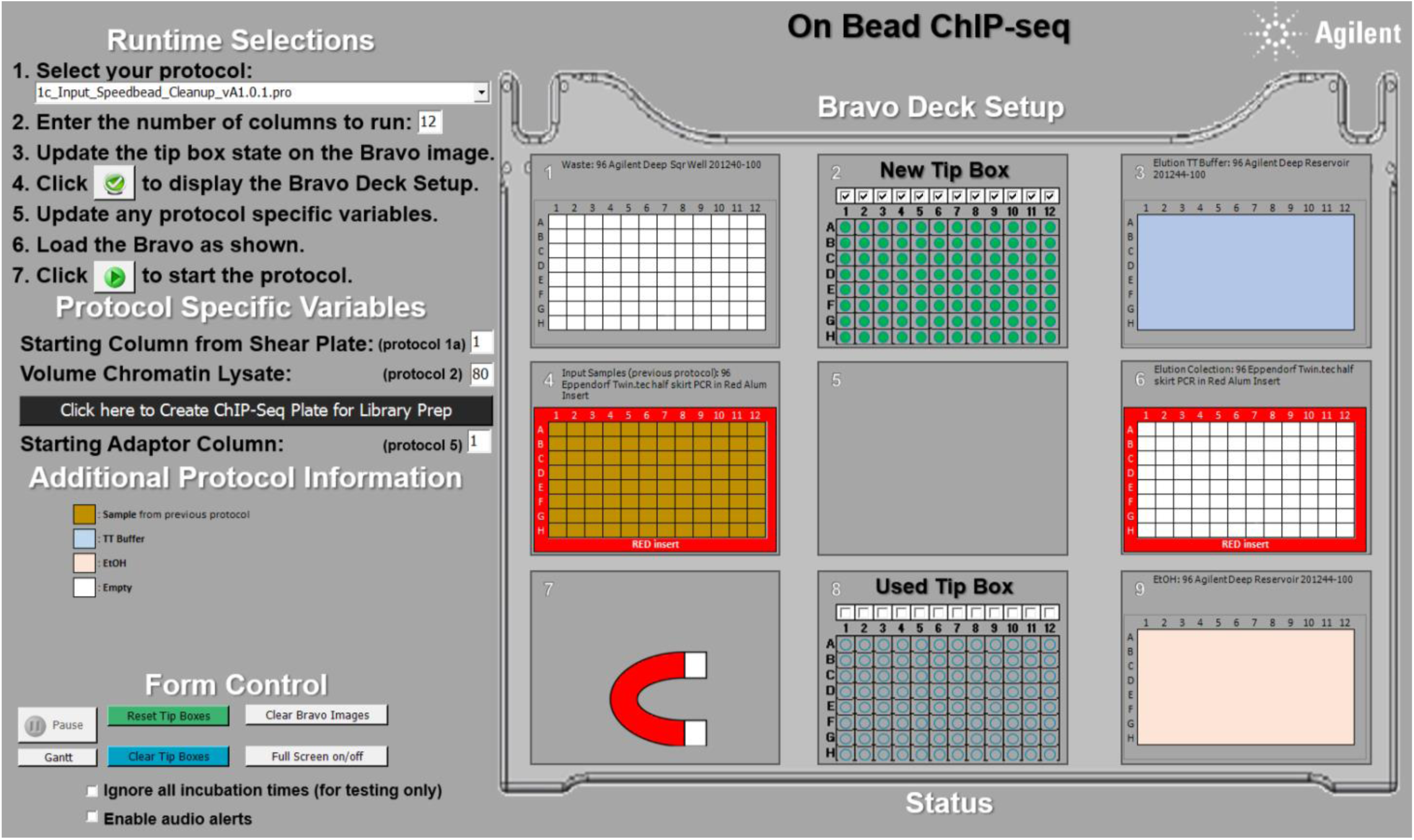 Bravo will perform the following steps:

- Place the plate to a magnet deck and aspirate the cleared supernatant.
- Wash the beads by adding 180 μL of 80% ethanol at room temperature and pipetting up and down to mix. Collect the beads on the magnet and discard the wash supernatant once cleared. Repeat this wash one additional time.
- After removing the second ethanol wash, air-dry the beads for ∼10 minutes, or until “cracks” just begin to appear in the packed beads.
- Elute DNA by adding 25 μL TT Buffer. Mix then incubate at room temperature for 5–10 minutes.
- Apply the plate to the magnet. Transfer the supernatant to a new plate and discard the Speedbeads.
11. At the end of the protocols, the **Sample Elution Plate** should contain cleaned DNA samples eluted in 25 μL of TT buffer.
12. To check the shearing, run 5 μL of the sample from each well on a 2% agarose gel at 120V for 10 minutes & 30 minutes.
13. The recommended length of shearing chromatin is 200 - 500 bp for ChIP assays. If shearing pattern is as expected, proceed to the next step. If storing the sheared chromatin for future use, combine the wells of the PIXUL shearing plate with the same cell type/condition into one tube and place at −80°C.

### Immunoprecipitation

1. Prepare 10 μL of Dynabeads Protein A or Dynabeads Protein G per sample into a microcentrifuge tube.

- **Note**: The choice of protein A and/or protein G Dynabeads, and the volume to use, is dictated by the antibody used for ChIP. Protein A binds rabbit antibodies with highest affinity, while protein G has higher affinity for mouse, rat, goat, and sheep antibodies.
2. Wash Dynabeads twice on magnet with equal volume of lysis buffer LB3/1% Triton X-100 + PIC.
3. For each antibody, prepare a mixture of antibody and Dynabeads Protein A/G in LB3 + PIC and 1% Triton X-100. Add the mixture to **Antibody/Dynabeads Mix Plate**.
4. Place **Antibody/Dynabeads Mix Plate** on **Bravo deck 5**, **empty tube strips** (with detached lid strips, without the lids) on **Bravo deck 6**, and **Chromatin Lysate Plate** on **Bravo deck 9**.
5. Select Bravo protocol **2_Chromatin_Beads_Antibody_On_Mag_vA1.0.1.pro** from the drop down manual in step 1 and click the green check symbol to display the Bravo Deck Setup. Update the protocol specific variables if necessary. The default volume of chromatin lysate is 80. Make sure the deck is set up as shown. Once ready, click on the right-pointing triangle (▷) at step 7 to start the protocol. 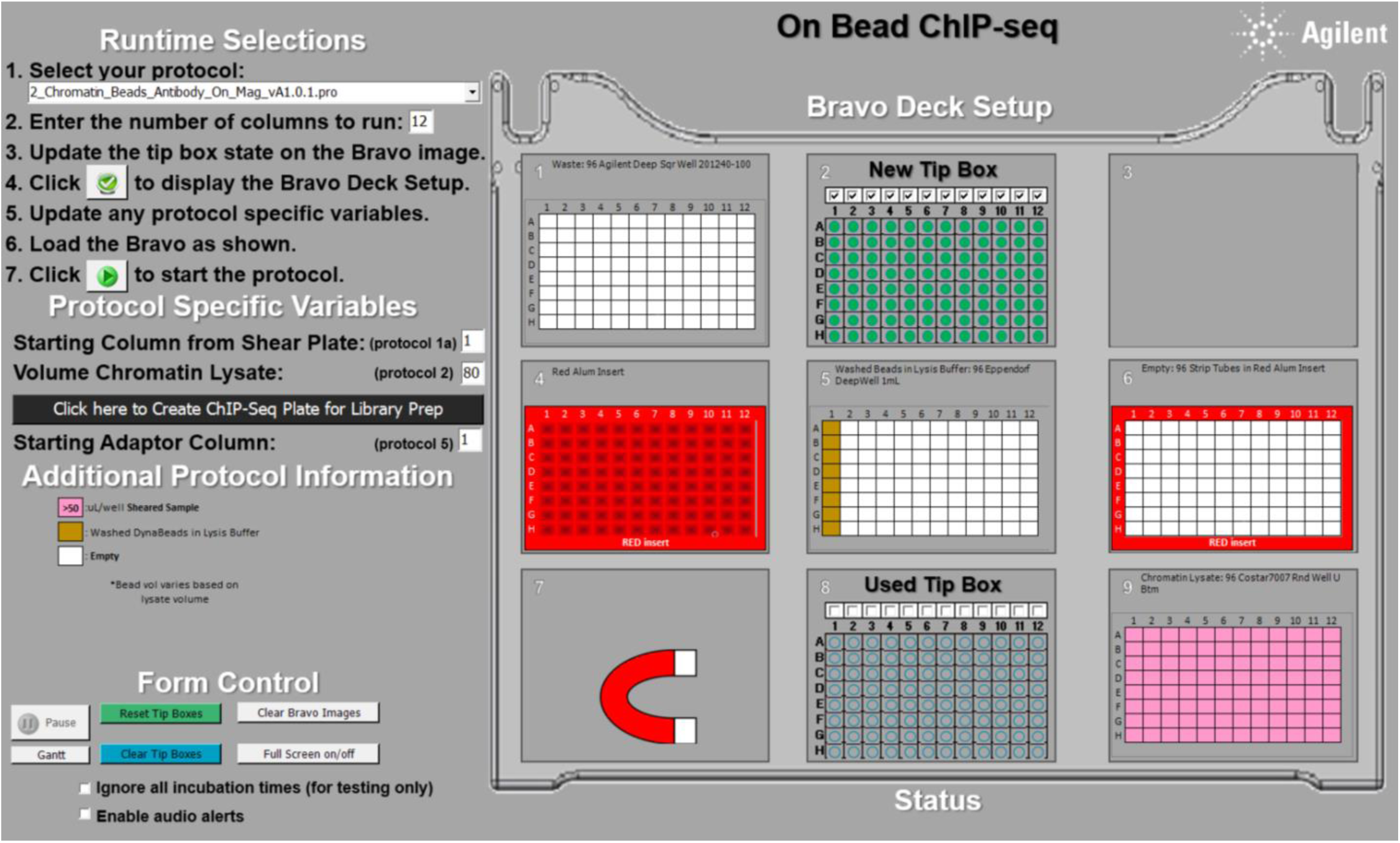 Bravo will perform the following steps:

- Distribute antibody/Dynabeads mix to each tube of an 8-tube strip accordingly.
- Aliquot appropriate amount (50-150μl) of sheared chromatin into each tube with the antibody/Dynabeads mix.
- Mix by pipetting up and down.
6. Incubate IP tube strips overnight (up to 16 h) at 4°C by rotating on a HulaMixer (setting below) in the cold room. 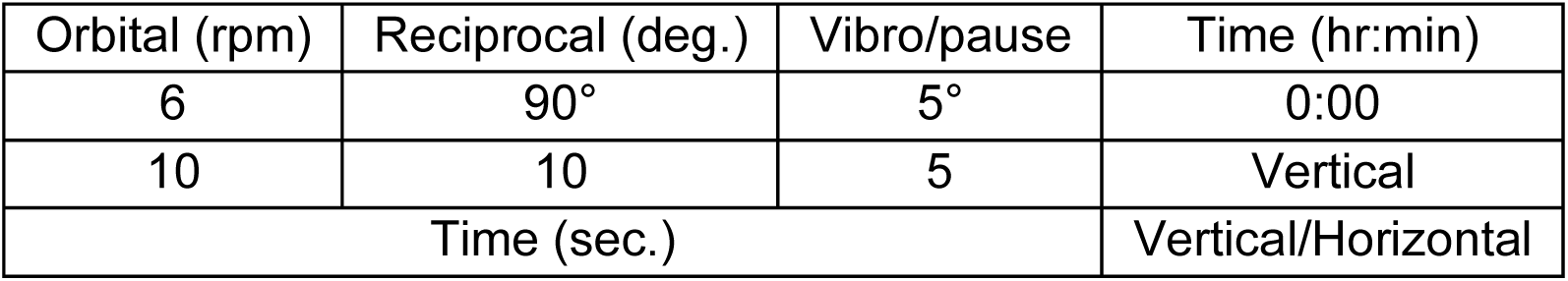

### Wash ChIP Samples

1. Freshly prepare the following amounts of ChIP Wash Buffer 1 (WBI), ChIP Wash Buffer 3 (WBIII), and TET Buffer by adding 25× protease inhibitor cocktail (PIC). Mix by vortexing and keep on ice. (The example below shows the amount required for a full 96-well plate.) 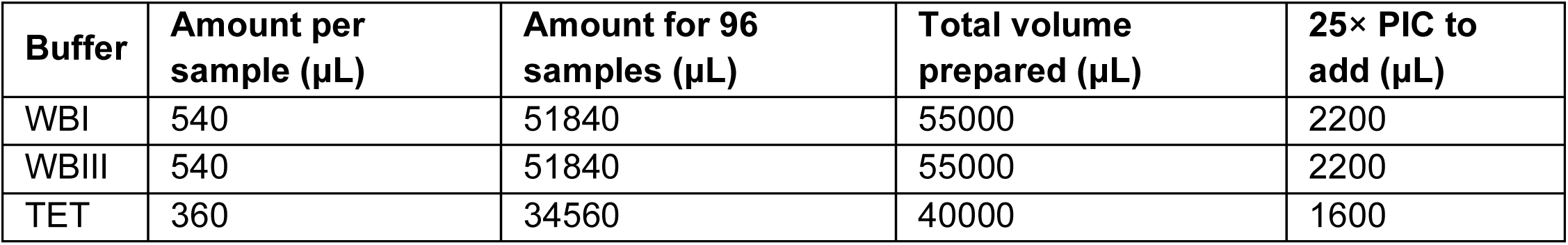
2. Place **Waste Plate** on **Bravo deck 1**, **TT Buffer Reservoir** on **Bravo deck 3**, **empty ChIP Plate** on **Bravo deck 4**, **IP tube strips** (without the lids) on **Bravo deck 6**, and **Wash Buffer Reservoir** (with WBI + PIC first) on Bravo deck 9. Ensure the reservoirs contain at least enough liquid to form a thin layer across the bottom.
3. Select Bravo protocol **3_ChIP_Wash_Elution_Off_Mag_vA1.0.1.pro** from the drop down manual in step 1 and click the green check symbol to display the Bravo Deck Setup. Make sure the deck is set up as shown. Once ready, click on the right-pointing triangle (▷) at step 7 to start the protocol. 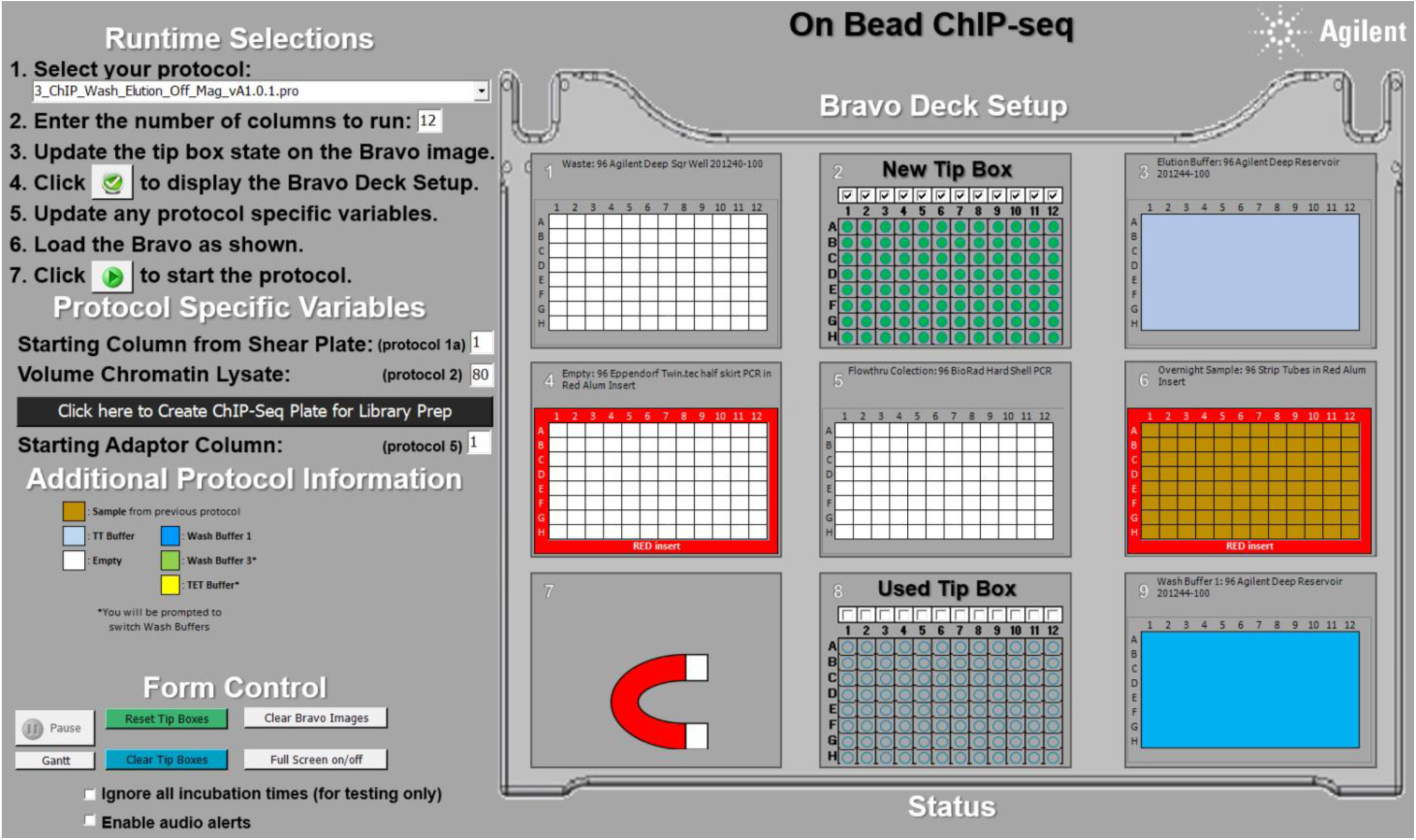 Bravo will perform the following steps:

- Wash the ChIP samples 3 times with 180 μL WBI + PIC
- Wash the ChIP samples 3 times with 180 μL WBIII + PIC
- Wash the ChIP samples 2 times with 180 μL TET + PIC
- Resuspend beads in 25 μL of TT buffer
4. At the end of the protocol, each well of the **ChIP plate** (deck 4) should contain 25μL of ChIP sample eluted in TT buffer.

### Library Preparation

1. Add each of the 25 μL of diluted input chromatin sample to the **ChIP Plate** manually or run Bravo protocol **4_prePrep_Combine_Samples_with_DilutedInputControls_vA1.0.1.pro** Manual steps for diluting chromatin lysate to use as input: For each condition, take 4 μL of the sheared chromatin lysate and add 46 μL of TT buffer to make a 1:12.5 dilution. Take 5 μL of the 1:12.5 diluted chromatin lysate and add 20 μL of TT buffer to make another 1:5 dilution. The total dilution of the chromatin lysate is 1:62.5.
2. Place the ChIP Plate on Bravo deck 4.
3. Freshly prepare the following **End Prep Mix** on ice. Each sample requires 5 μL and it is recommended to prepare at least 20% more reaction volumes per sample. Aliquot 1/8 of the total volume of the mastermix to each well in the first column of the End Prep Mix Plate. Place the **End Prep Mix Plate** on **Bravo deck 6**. (The example below shows the amount required for a full 96-well plate.) 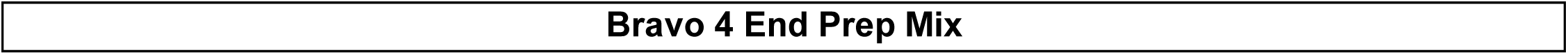
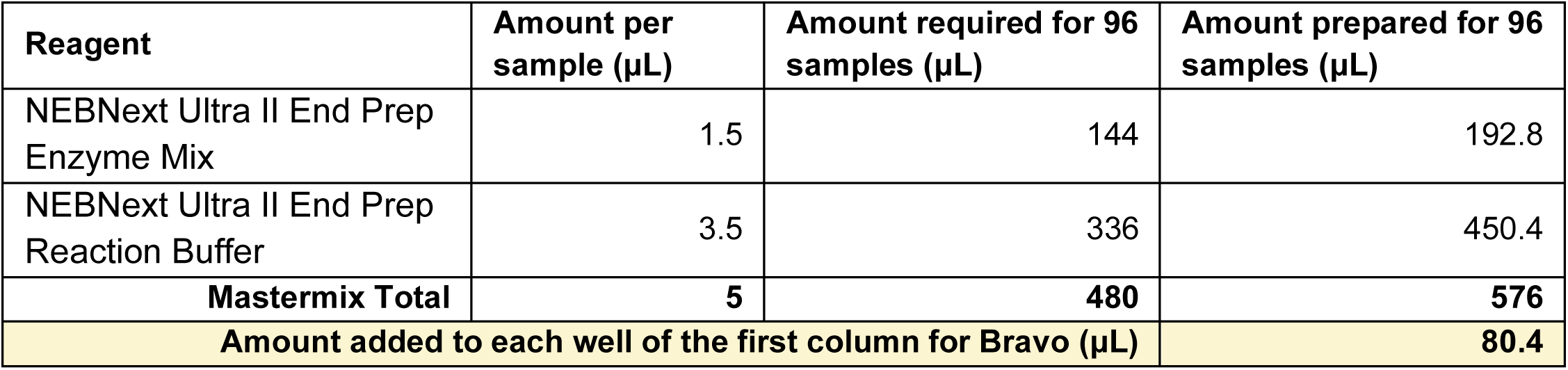
4. Select Bravo protocol **4_End_Repair_MM_Dispense_vA1.0.1.pro** from the drop down manual in step 1 and click the green check symbol to display the Bravo Deck Setup. Make sure the deck is set up as shown. Once ready, click on the right-pointing triangle (▷) at step 7 to start the protocol. 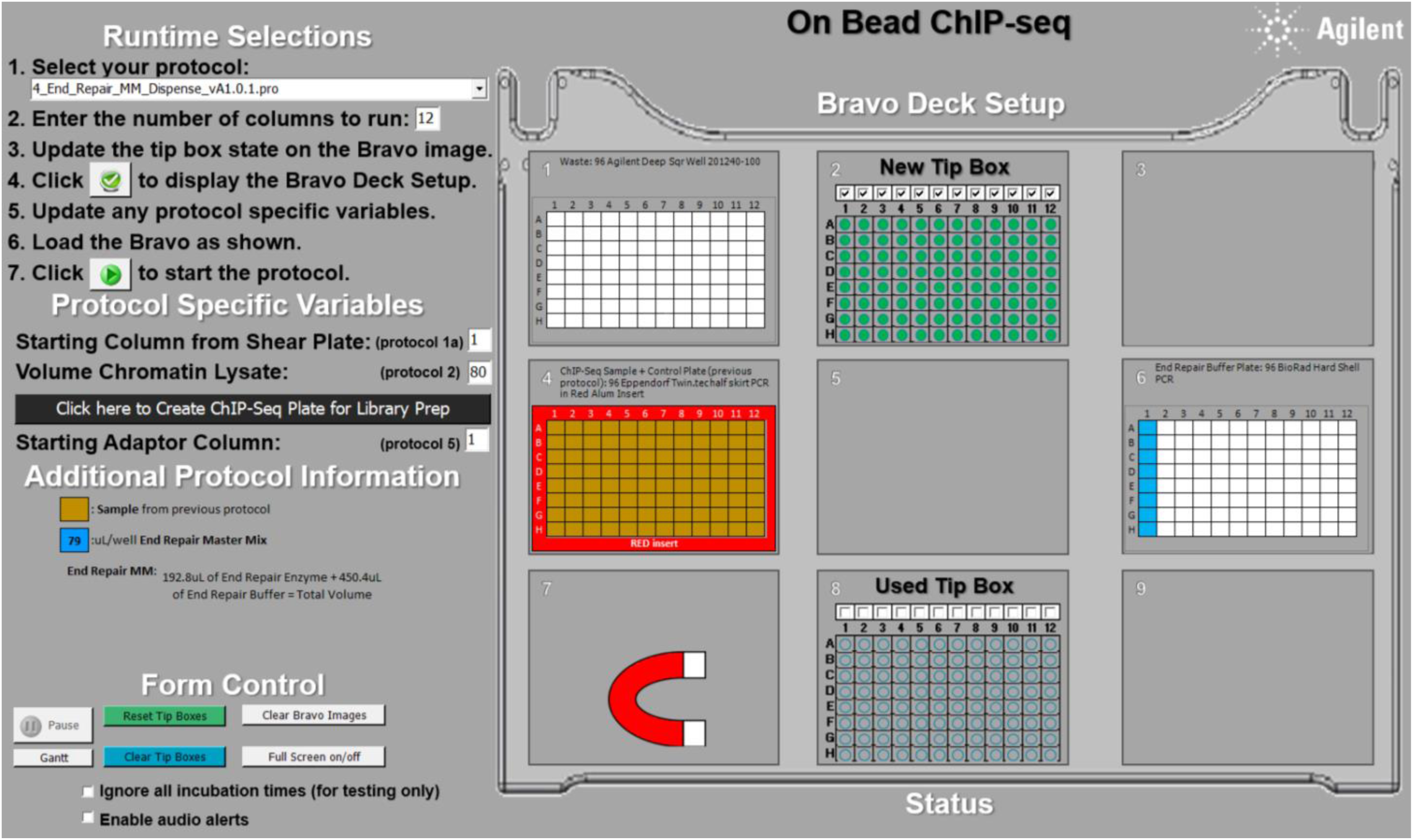 Bravo will perform the following steps:

- Add 5 uL End Prep Mix to each sample.
- Mix up and down at least 10 times (the mix cycle can be changed in the parameters).
5. Incubate for 30 minutes at 20°C, then 30 minutes at 65°C, then hold at 4°C in a PCR cycler with the lid set to 75°C (can leave at 4°C for ∼ 1 hour with no problems).
6. Freshly prepare the following **Ligation Mix** on ice. Each sample requires 15.5 μL and it is recommended to prepare 20% more reaction volumes per sample. Aliquot 1/8 of the total volume of the mastermix to each well in the first column of the **Ligation Mix Plate**. Place the **Ligation Mix Plate** on **Bravo deck 6**. (The example below shows the amount required for a full 96-well plate.) 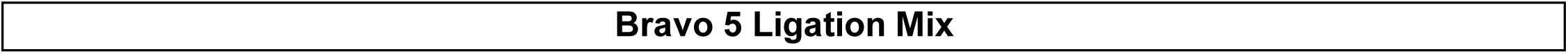
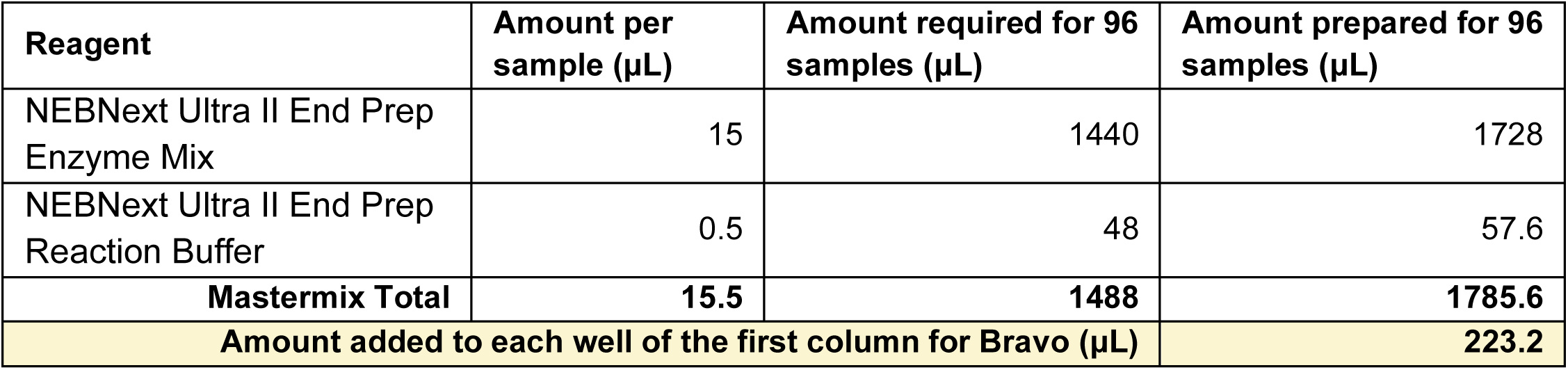
7. After incubation, place the **ChIP Plate** on **Bravo deck 4** and the **Index Plate** on **Bravo deck** The indexing adapters we usually use are from NEXTflex® Unique Dual Index Barcodes (Set B: UDI#97-192) (Catalog #NOVA-514151) (1:100 diluted 0.25 µM). Adapters can be diluted with 1× T4 DNA ligase buffer (50 mM Tris-HCl, 10 mM MgCl2, 1 mM ATP, 10 mM DTT, pH 7.5 at 25°C).
8. Select Bravo protocol **5_Adapter_Ligation_MM_vA1.0.1.pro** from the drop down manual in step 1 and click the green check symbol to display the Bravo Deck Setup. Make sure the deck is set up as shown. Once ready, click on the right-pointing triangle (▷) at step 7 to start the protocol. 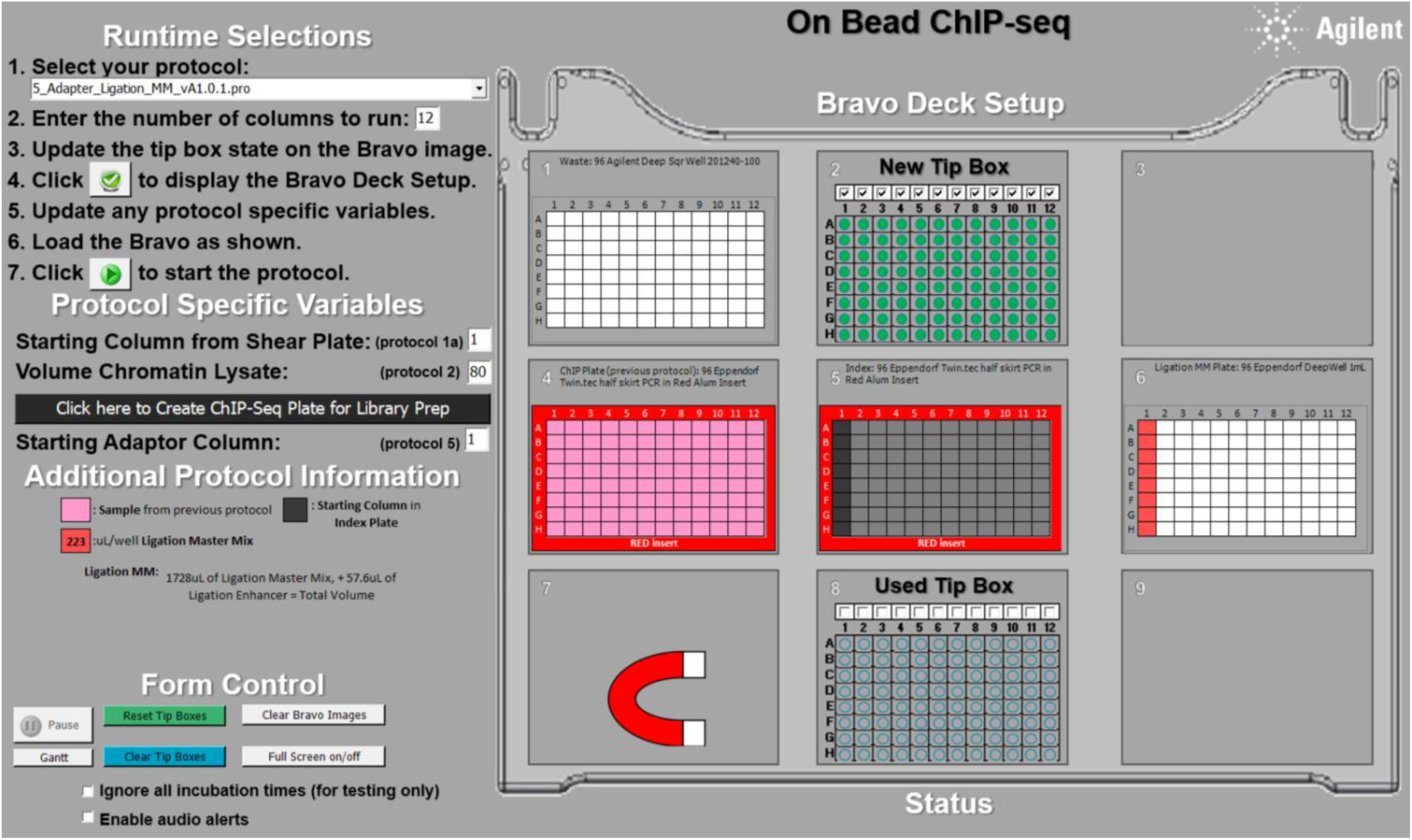 Bravo will perform the following steps:

- Add 15.5 μL of the Ligation Mix to each sample.
- Add 2.5 μL of Index Barcodes (1:100 diluted 0.25μM) to each sample.
- Mix up and down at least 10 times (the mix cycle can be changed in the parameters).
9. Incubate the plate at 20°C for 15 minutes in a PCR cycler with the heated lid set to “off.”
10. Freshly prepare the following **Ligation Stop Solution** at room temperature. Each sample requires 27.5 μL and it is recommended to prepare 1.2 reaction volumes per sample. Aliquot 1/8 of the total volume of the mastermix to each well in the first column of the **Stop Solution Plate**. Place the **Stop Solution Plate** on **Bravo deck 6**. (The example below shows the amount required for a full 96-well plate.) 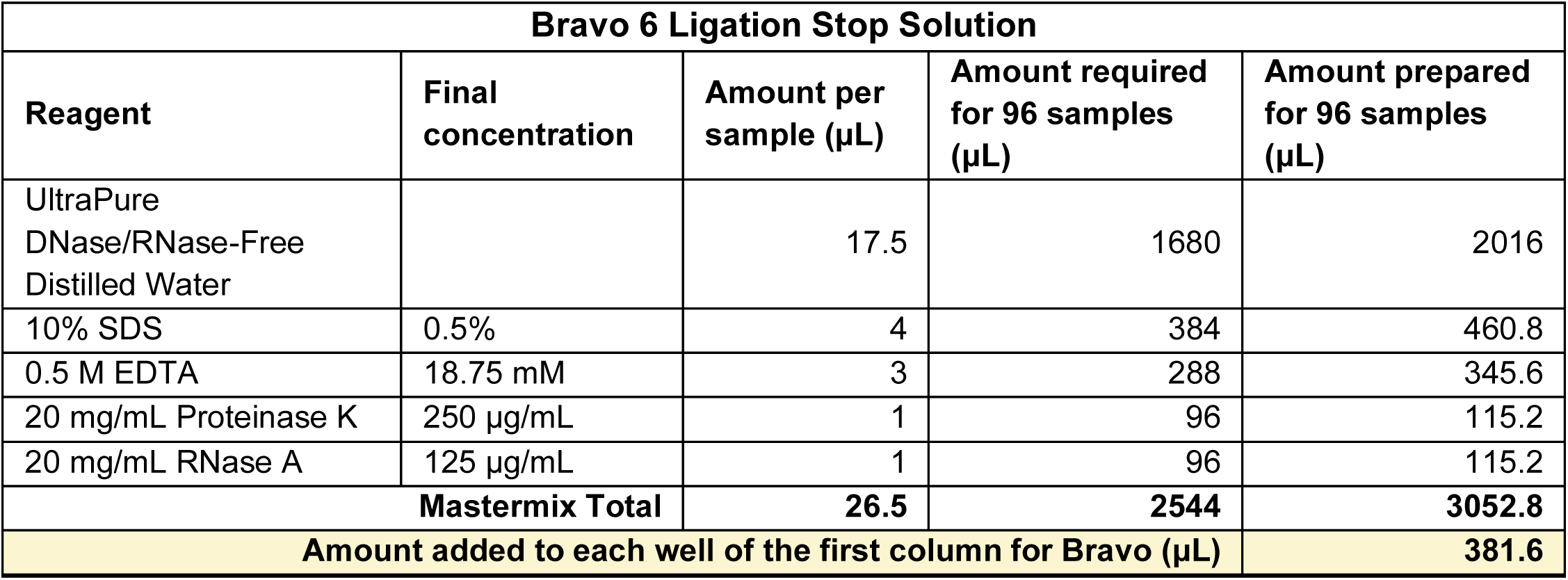
11. After incubation, place the **ChIP Plate** on **Bravo deck 4** and **5M NaCl Reservoir** on **Bravo deck 9**. Ensure the reservoirs contain at least enough liquid to form a thin layer across the bottom.
12. Select Bravo protocol **6_Ligation_Stop_Solution_Disp_vA1.0.1.pro** from the drop down manual in step 1 and click the green check symbol to display the Bravo Deck Setup. Make sure the deck is set up as shown. Once ready, click on the right-pointing triangle (▷) at step 7 to start the protocol. 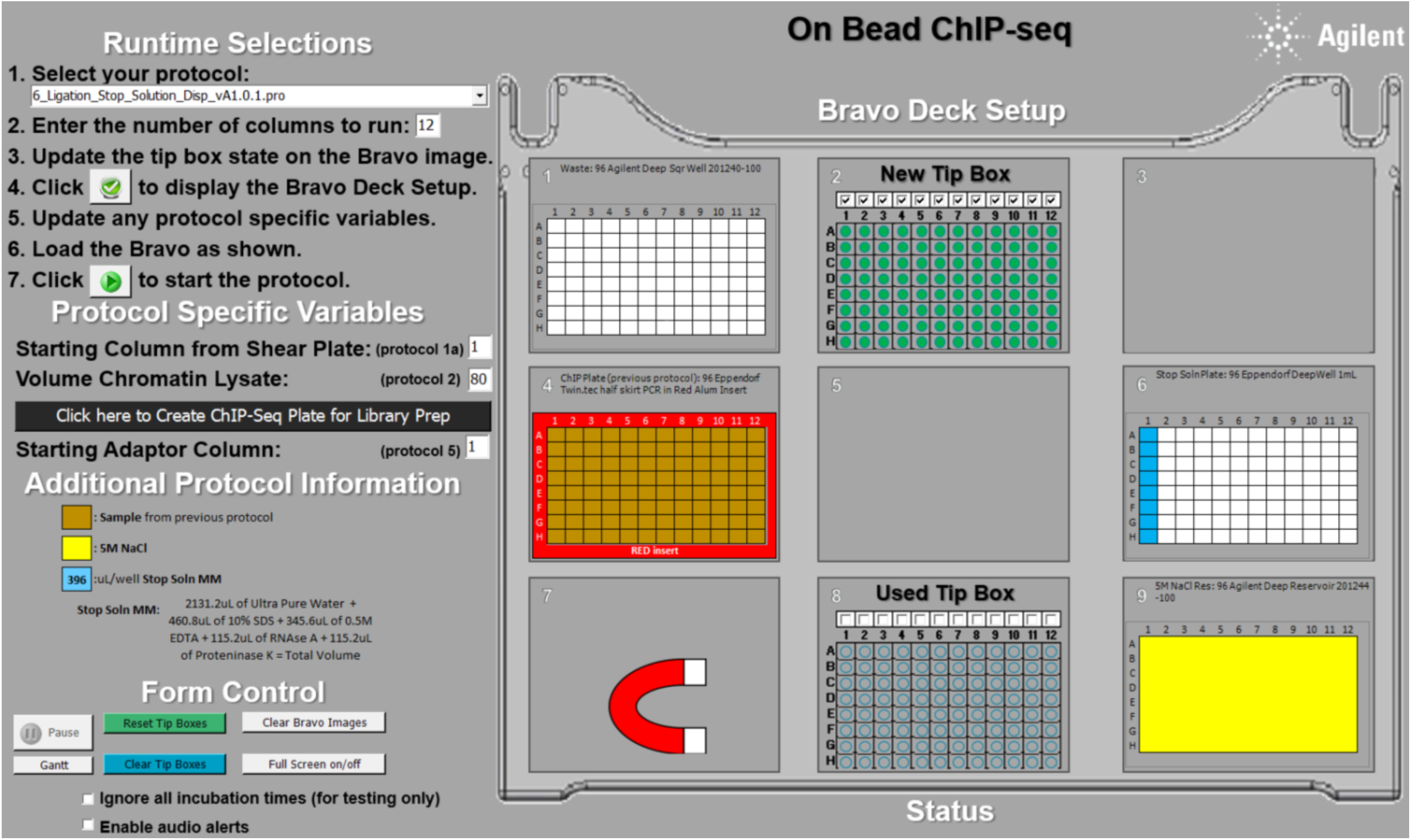 Bravo will perform the following steps:

- Add 27.5 μL of the Ligation Stop Solution to each sample.
- Add 4.5 μL of 5 M NaCl to each sample.
- Mix up and down at least 10 times (the mix cycle can be changed in the parameters).
13. **Reverse crosslinking**: Incubate at 55°C for 1 hour to digest RNA and proteins, then shift the incubation temperature to 65°C for at least 2 hours (up to 16 hours) in a PCR cycler with the heated lid set to at least 75°C to prevent evaporation and drying out of the samples. This can be left at 12°C or 4°C overnight.
14. Freshly prepare the following **Speedbeads Mastermix** at room temperature. Each sample requires 63 μL and it is recommended to prepare 20% more reaction volumes per sample. Aliquot 1/8 of the total volume of the mastermix to each well in the first column of the **Speedbeads Mastermix Plate**. Place the **Speedbeads Mastermix Plate** on **Bravo deck 5**. (The example below shows the amount required for a full 96-well plate.) 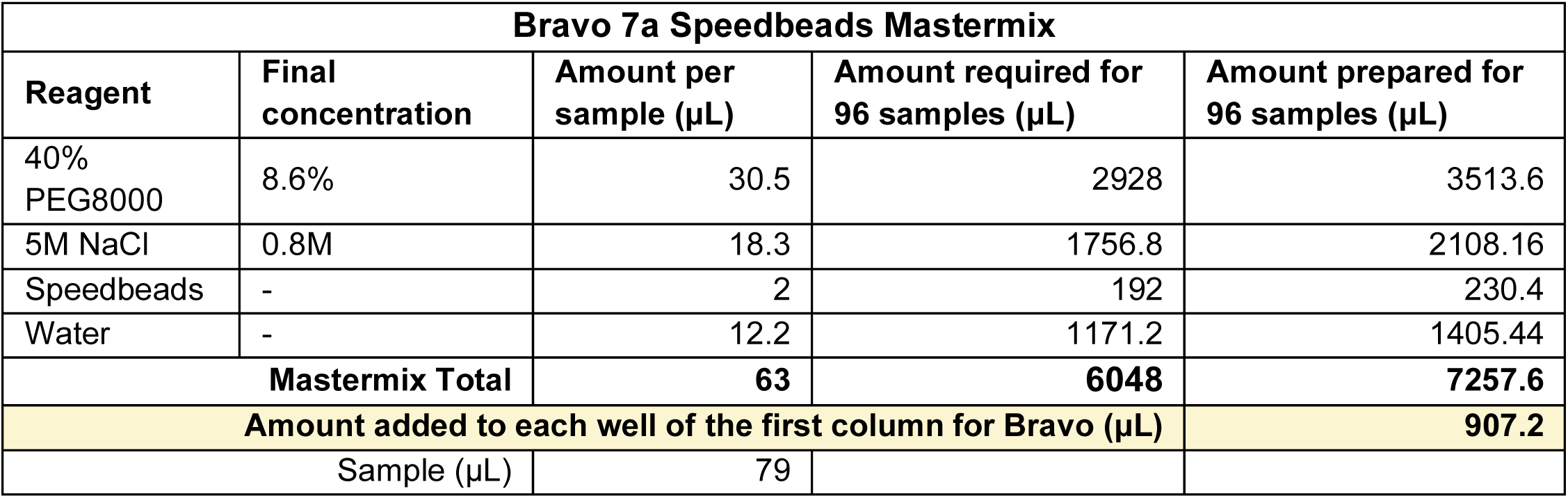
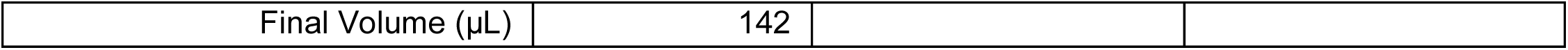
15. After incubation, place the **ChIP Plate** back to **Bravo deck 4**.
16. Select Bravo protocol **7a_Adapter_Ligation_Cleanup_vA1.0.1.pro** from the drop down manual in step 1 and click the green check symbol to display the Bravo Deck Setup. Make sure the deck is set up as shown. Once ready, click on the right-pointing triangle (▷) at step 7 to start the protocol. 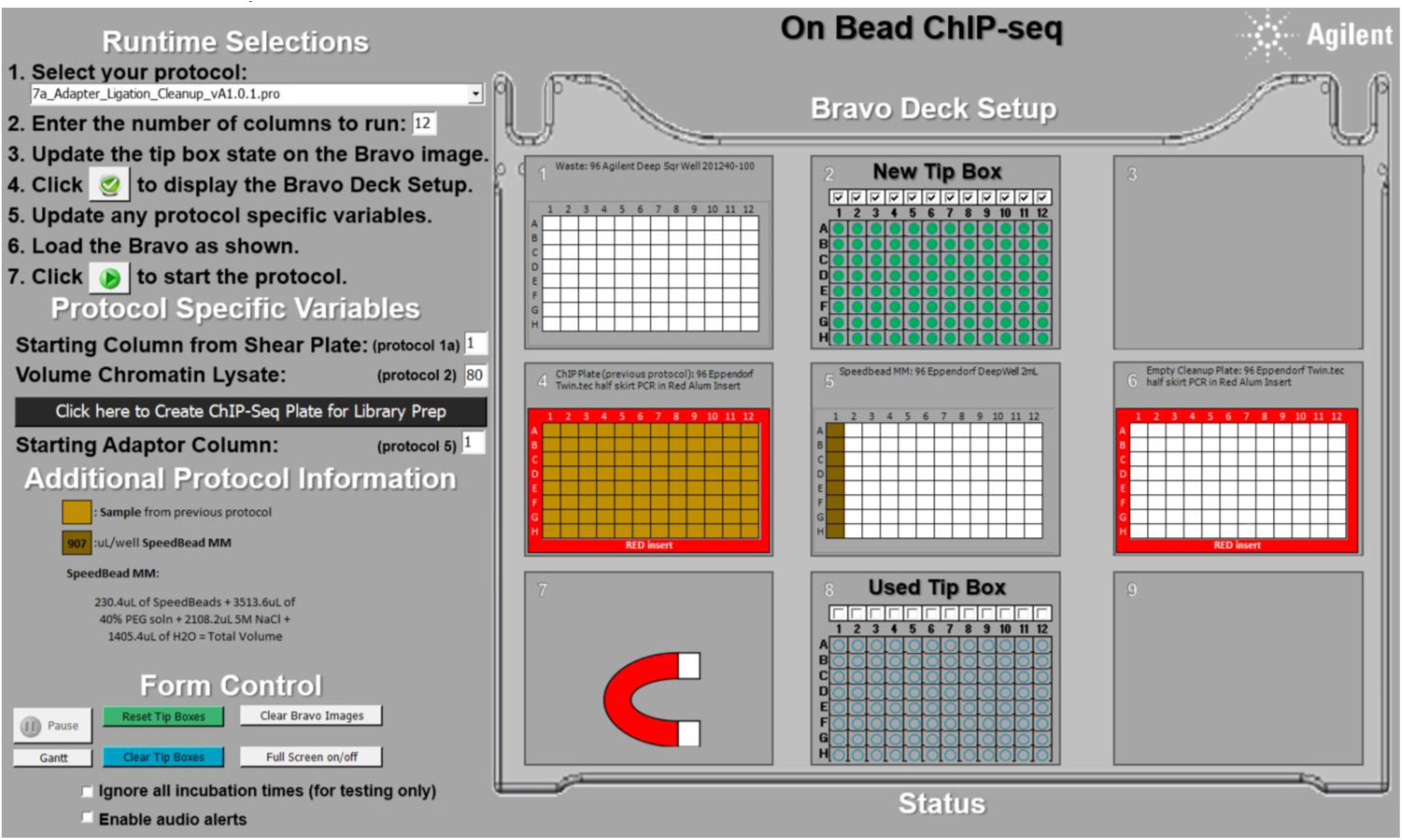 Bravo will perform the following steps:

- Transfer the supernatant from the ChIP plate with Dynabeads to the empty ChIP Cleanup Plate.
- Add 63 μL of Speedbeads mastermix to each 79 μL ChIP sample. The final concentration of PEG will be 8.6% and the final concentration of NaCl will be 0.8 M.
- Mix the samples to homogeneity by vortexing or repetitive pipetting. Incubate at room temperature for 10 minutes.
17. During incubation, place the **Waste Plate** on **Bravo deck 1**, **TT Buffer Reservoir** on **Bravo deck 3**, **empty Elution Plate** on **Bravo deck 6**, and **80% Ethanol Reservoir** on **Bravo deck 9**. Ensure the reservoirs contain at least enough liquid to form a thin layer across the bottom.
18. Immediately after protocol 7a is completed, select Bravo protocol **7b_Adapter_Ligation_Cleanup_vA1.0.1.pro** from the drop down manual in step 1 and click the green check symbol to display the Bravo Deck Setup. Make sure the deck is set up as shown. Once ready, click on the right-pointing triangle (▷) at step 7 to start the protocol. 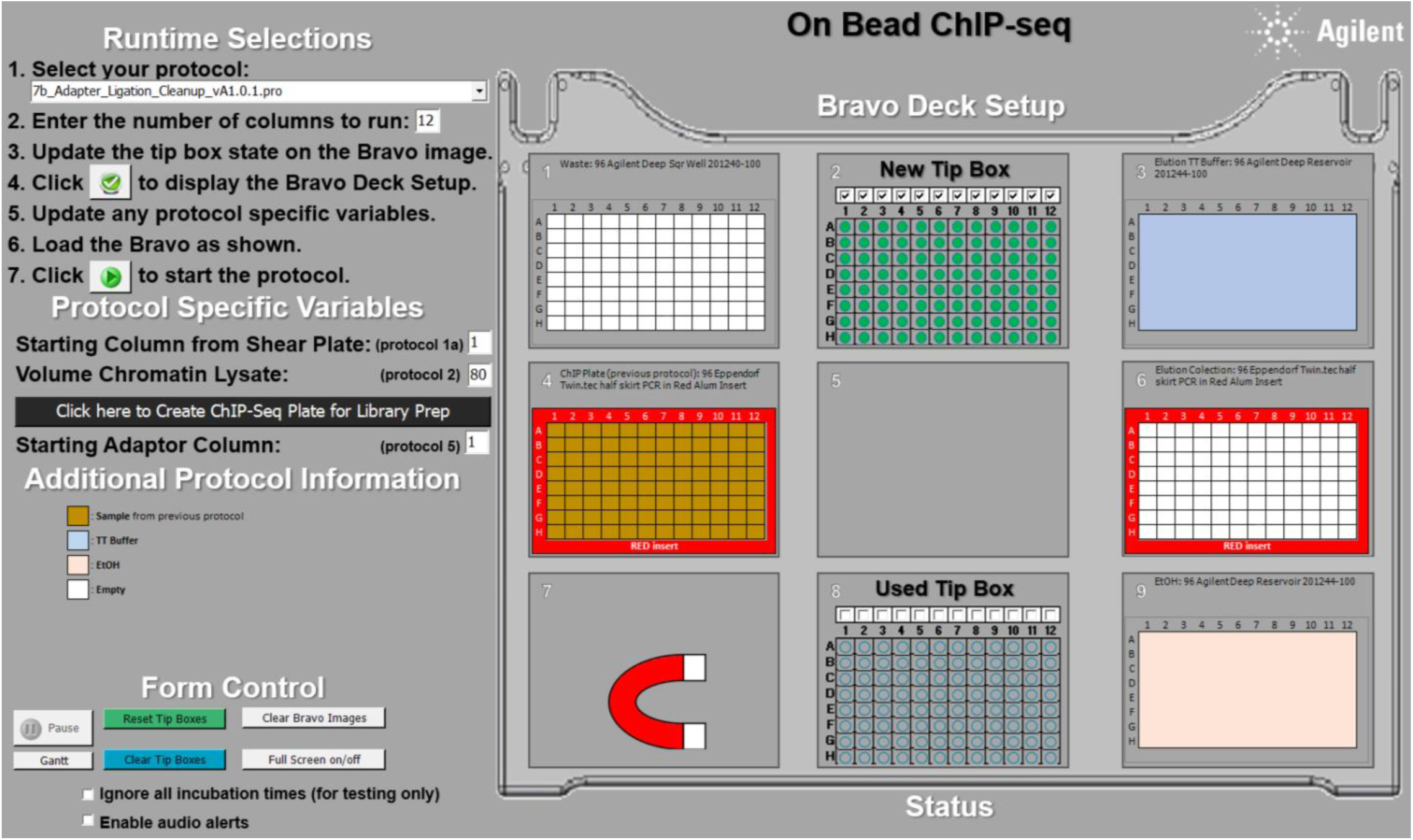 Bravo will perform the following steps:

- Apply the plate to a magnet and aspirate the cleared supernatant.
- Wash the beads by adding 180 μL of 80% ethanol at room temperature and moving the plate to either side of the magnet six times. Collect the beads on the magnet and discard the supernatant once cleared. Repeat this wash one additional time.
- After removing the second ethanol wash, air-dry the beads for 10 minutes, or until “cracks” just begin to appear in the packed beads.
- Elute DNA by adding 25 μL TT Buffer. Mix then incubate at room temperature for 5–10 minutes.
- Apply the plate to the magnet. Transfer the supernatant to a new plate and discard the Speedbeads.
19. At the end of the protocols, the **ChIP Plate** (deck 6) should contain ChIP samples eluted in 25 μL of TT buffer. Move the **ChIP Plate** to **Bravo deck 4** before proceeding to the next protocol.
20. Freshly prepare the following **PCR Mastermix**. Each sample requires 25.5 μL and it is recommended to prepare 20% more reaction volumes per sample. Aliquot appropriate amount to each well in the first column of the **PCR Mastermix Plate.** Place the **PCR Mastermix Plate** on **Bravo deck 6**. (The example below shows the amount required for a full 96-well plate.) 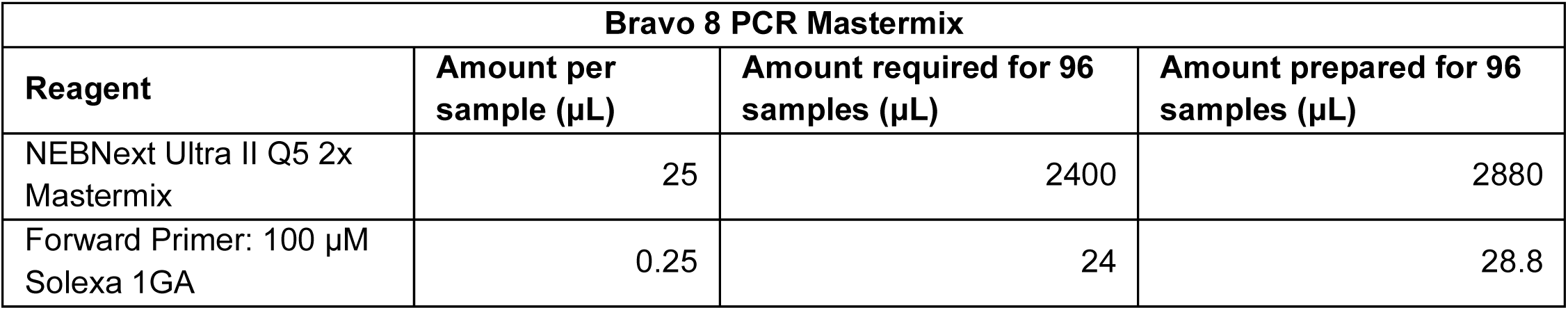
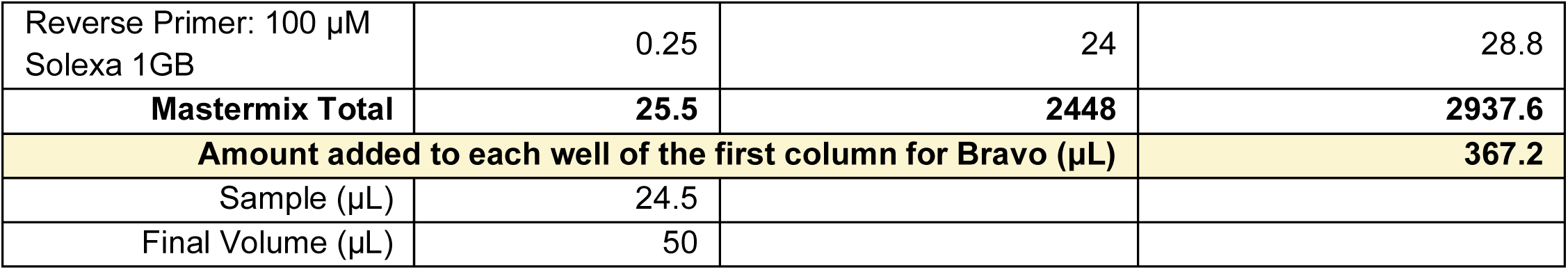
21. Select Bravo protocol **8_PCR_Enrichment_MM_Dispense_vA1.0.1.pro** from the drop down manual in step 1 and click the green check symbol to display the Bravo Deck Setup. Make sure the deck is set up as shown. Once ready, click on the right-pointing triangle (▷) at step 7 to start the protocol. 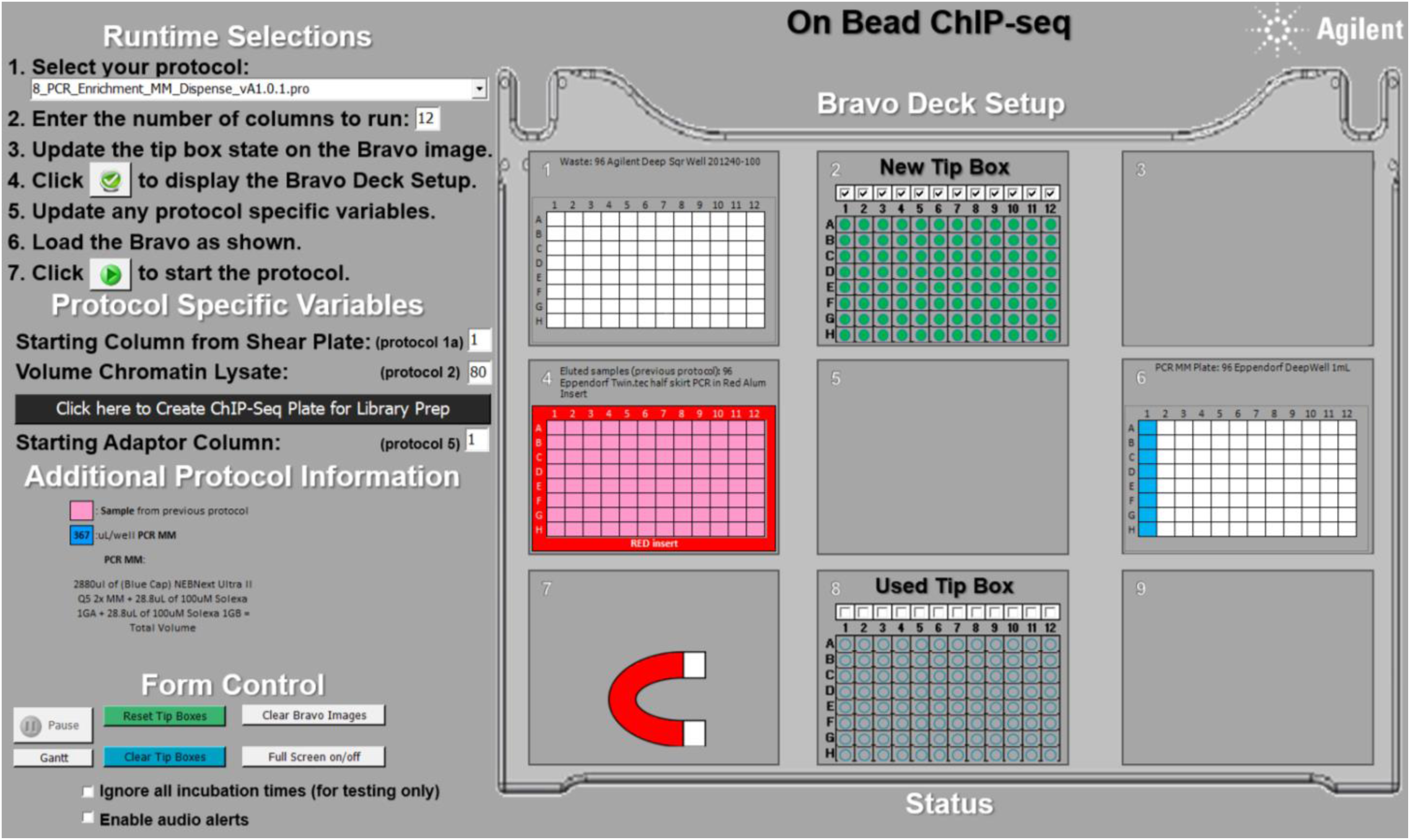 Bravo will perform the following steps:

- Add 25.5 μL of the PCR Mastermix to each sample.
- Mix up and down at least 10 times (the mix cycle can be changed in the parameters).
22. PCR amplify using a PCR cycler with a heated lid set to 105°C. 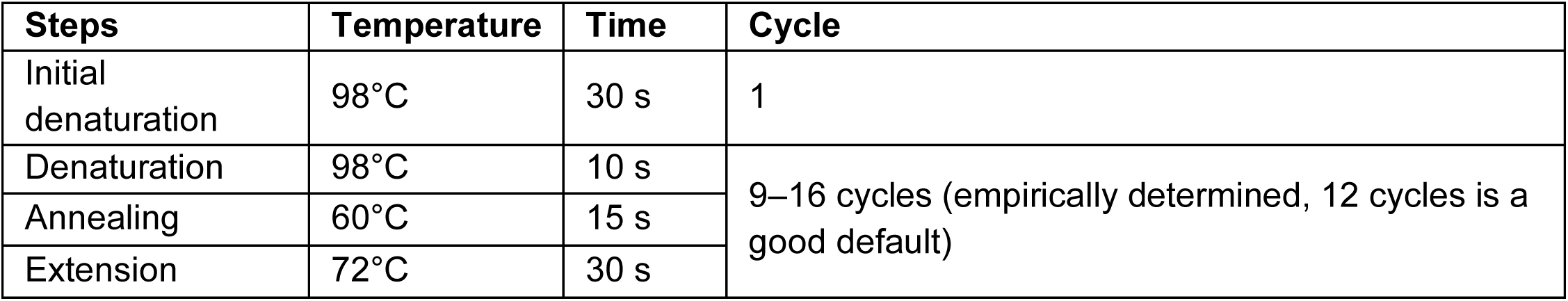
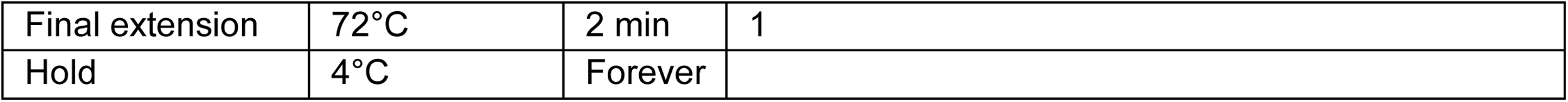
23. Freshly prepare the following **Speedbeads Mastermix** at room temperature. Each sample requires 40.5 μL and it is recommended to prepare 20% more reaction volumes per sample. Aliquot 1/8 of the total volume of the mastermix to each well in the first column of the **Speedbeads Mastermix Plate**. Place the **Speedbeads Mastermix Plate** on **Bravo deck 5**. (The example below shows the amount required for a full 96-well plate.)

- Note: The low (8.5%) PEG8000 amount size selects against short adapter dimers (∼125 bp size). 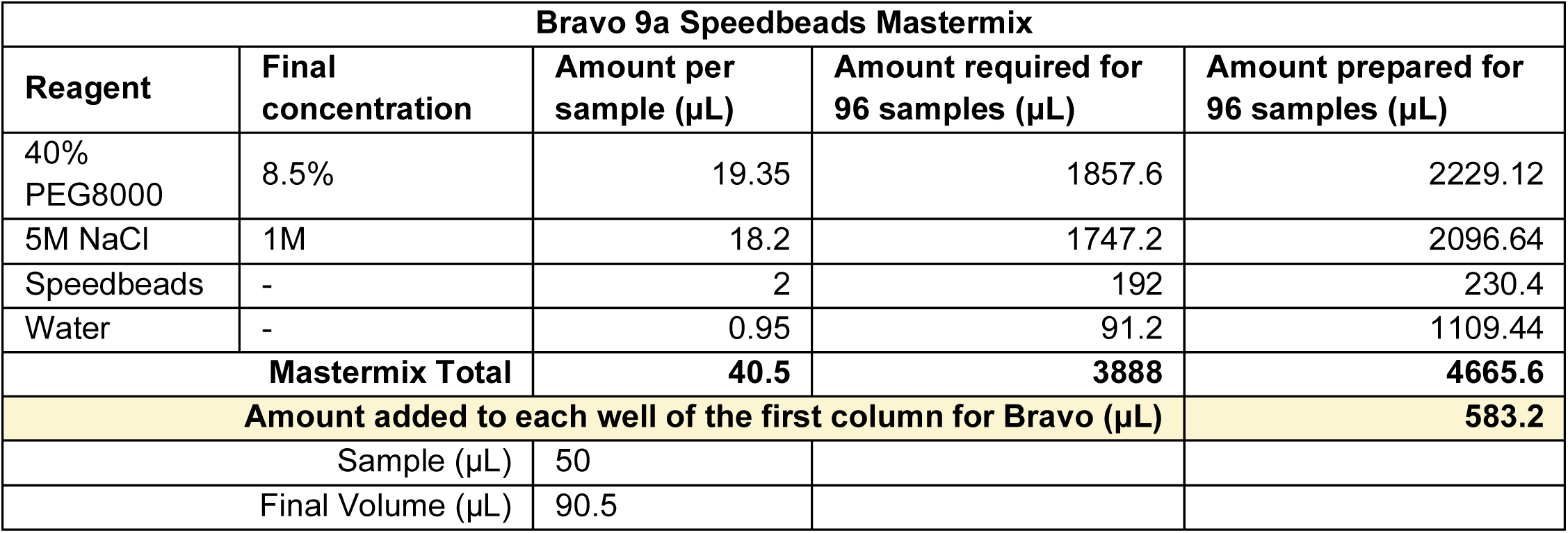
24. After PCR, place the **ChIP Plate** back to **Bravo deck 4**.
25. Select Bravo protocol **9a_PCR_Enrichment_Cleanup v2_vA1.0.1.pro** from the drop down manual in step 1 and click the green check symbol to display the Bravo Deck Setup. Make sure the deck is set up as shown. Once ready, click on the right-pointing triangle (▷) at step 7 to start the protocol. 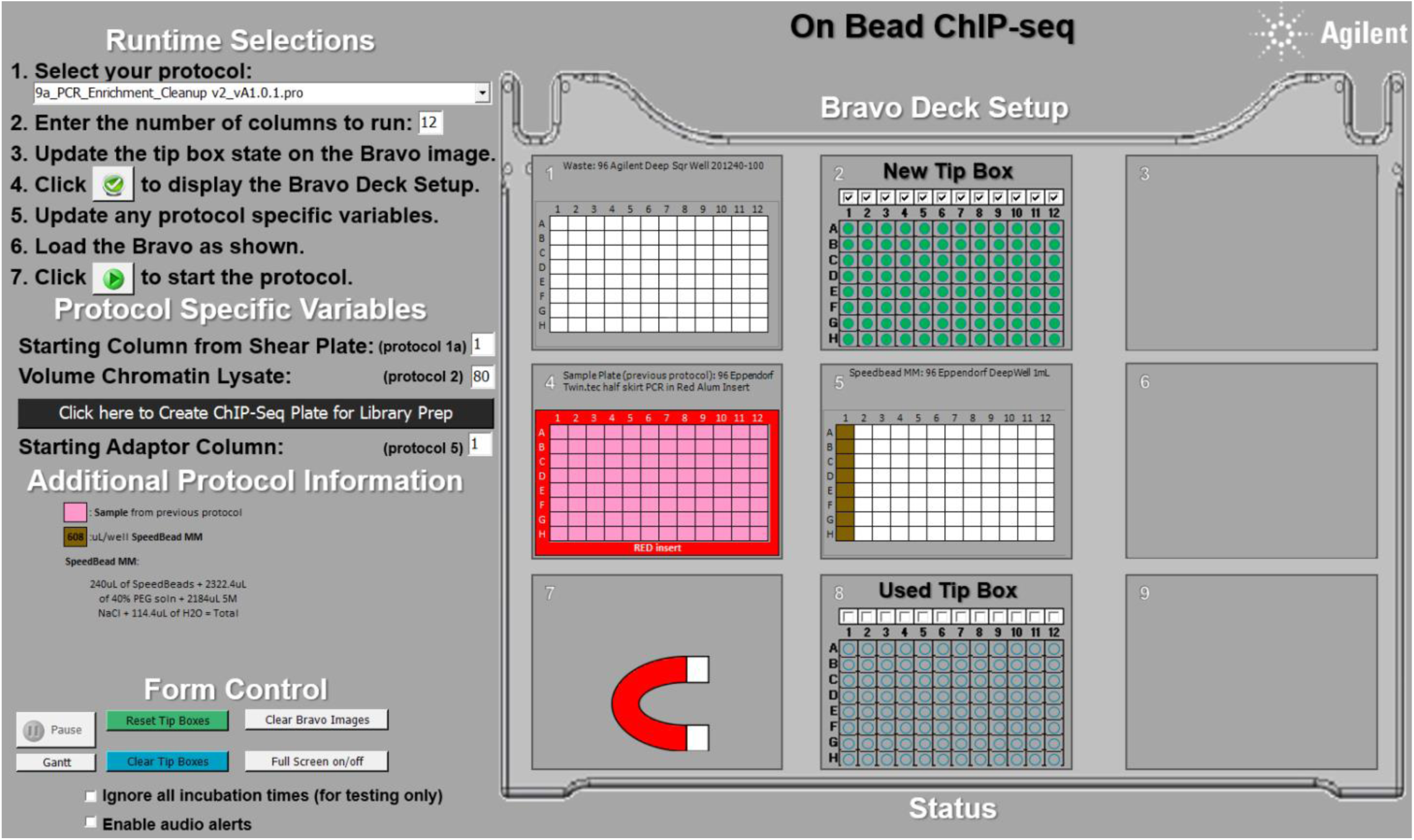 Bravo will perform the following steps:

- Add 40.5 μL of Speedbead/PEG mastermix to each 50 μL ChIP sample. The final concentration of PEG8000 will be 8.5% and the final concentration of NaCl will be 1 M.
- Mix the samples to homogeneity by vortexing or repetitive pipetting. Incubate at room temperature for 10 min.
26. During incubation, place the **Waste Plate** on **Bravo deck 1**, **TT Buffer Reservoir** on **Bravo deck 3**, **empty Elution Plate** on **Bravo deck 6**, and **80% Ethanol Reservoir** on **Bravo deck 9**. Ensure the reservoirs contain at least enough liquid to form a thin layer across the bottom.
27. Immediately after protocol 9a is completed, select Bravo protocol **9b_PCR_Enrichment_Cleanup v2_vA1.0.1.pro** from the drop down manual in step 1 and click the green check symbol to display the Bravo Deck Setup. Make sure the deck is set up as shown. Once ready, click on the right-pointing triangle (▷) at step 7 to start the protocol. 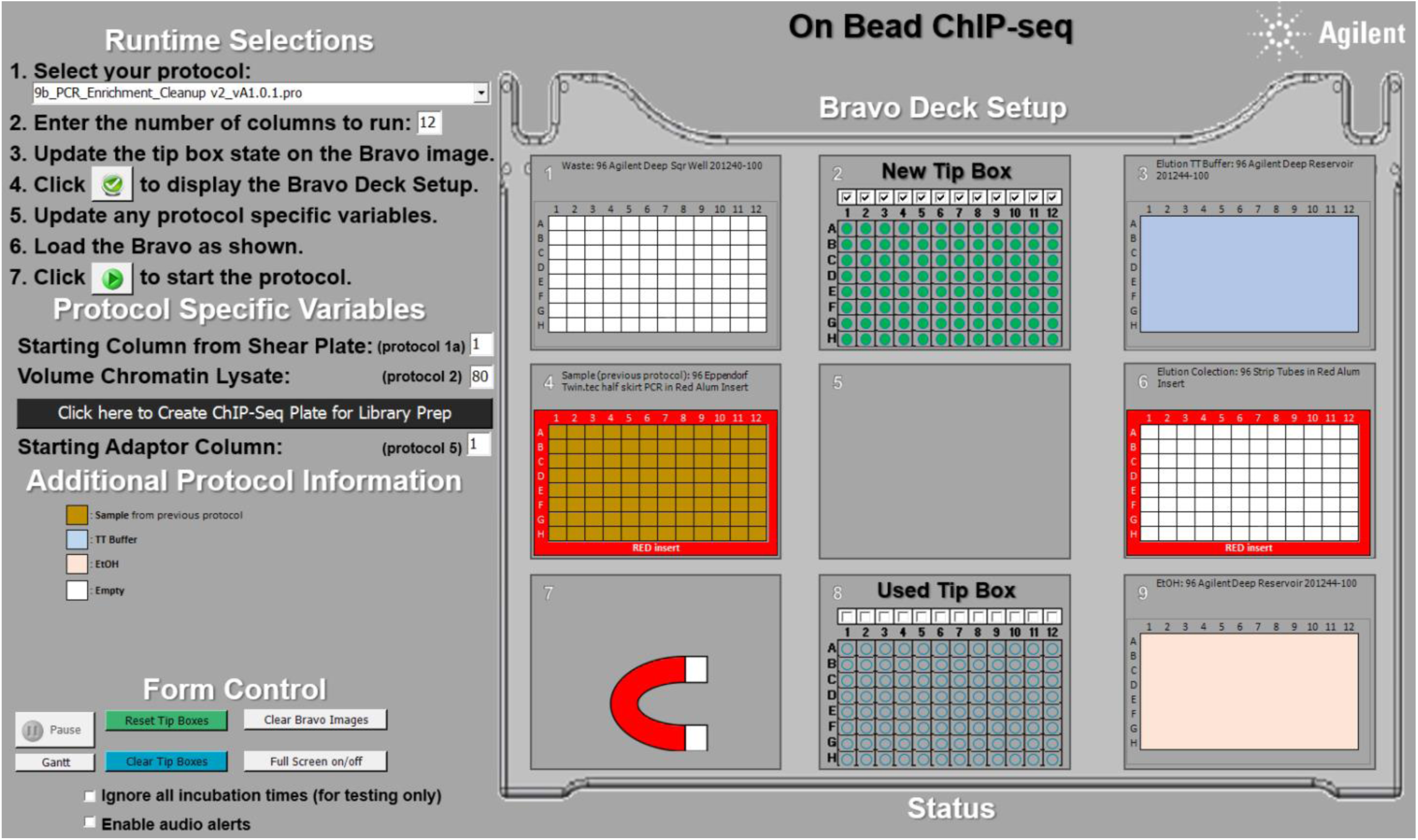 Bravo will perform the following steps:

- Apply the plate to a magnet and aspirate the cleared supernatant.
- Wash the beads by adding 180 μL of 80% ethanol at room temperature and moving the plate to either side of the magnet six times. Collect the beads on the magnet and discard the supernatant once cleared. Repeat this wash one additional time.
- After removing the second ethanol wash, air-dry the beads for ∼10 minutes, or until “cracks” just begin to appear in the packed beads.
- Elute DNA by adding 25 μL TT Buffer. Mix then incubate at room temperature for 5–10 minutes.
- Apply the plate to the magnet. Transfer the supernatant to a new plate and discard the Speedbeads.
28. At the end of the protocols, the **ChIP Elution Plate** (deck 6) should contain ChIP library samples eluted in 25 μL TT buffer.

### DNA Quantification

1. Size: Run 5 μL of DNA library + 1 μL of 6X TriTrack DNA Loading Dye on a 2% agarose gel pre-stained with GelGreen.
2. Concentration: Use 1 μL of each library with Qubit HS DNA buffer and standards.
3. Store the remaining DNA library samples at -20°C until ready for sequencing.

