## Supplementary material for "Automated chromatin profiling with spa-ChIP-seq uncovers the impacts of condition variations": Cao-etal-revised-2025-biorxiv-automation-files: 96LT_Tip_Images_for_Form.js

```
/***********************************
************************************
****** Set Default Variables  ******
************************************
***********************************/
//image locations

var myRoot_NTip =  masterFilePath + 'Forms/Supporting Files/Tips/Tips_no.png';
var myRoot_YTip =  masterFilePath + 'Forms/Supporting Files/Tips/Tips_yes.png';

//Loop variables
//Columns
var columns = 12;
var tipsOn_Col = 0;
var tipsOff_Col = 0;
var colCount = 0;

//Tip Box
var tipBoxLoc = 0;


/***********************************
************************************
******  Set Arrays from Form  ******
************************************
***********************************/

//***Columns**\\
//set arrays for tip cols
var tips_ColOn = 
[
 "box2_1",
 "box2_2",
 "box2_3",
 "box2_4",
 "box2_5",
 "box2_6",
 "box2_7",
 "box2_8",
 "box2_9",
 "box2_10",
 "box2_11",
 "box2_12"
]

var tips_ColOff =
[
 "box8_1",
 "box8_2",
 "box8_3",
 "box8_4",
 "box8_5",
 "box8_6",
 "box8_7",
 "box8_8",
 "box8_9",
 "box8_10",
 "box8_11",
 "box8_12"
]

//***Images**\\
//set arrays for images
var tips_ImgOn = 
[
 "TipsOn_1",
 "TipsOn_2",
 "TipsOn_3",
 "TipsOn_4",
 "TipsOn_5",
 "TipsOn_6",
 "TipsOn_7",
 "TipsOn_8",
 "TipsOn_9",
 "TipsOn_10",
 "TipsOn_11",
 "TipsOn_12"
]

var tips_ImgOff = 
[
 "TipsOff_1",
 "TipsOff_2",
 "TipsOff_3",
 "TipsOff_4",
 "TipsOff_5",
 "TipsOff_6",
 "TipsOff_7",
 "TipsOff_8",
 "TipsOff_9",
 "TipsOff_10",
 "TipsOff_11",
 "TipsOff_12"
]

/***********************************
************************************
******Set TipBox Images 96 LT******
************************************
***********************************/

//set images for Source Box based on the existing checkmarks
function setSourceBoxImg_96()
{
	//trying loop thru img array instead
	//reset images for all the tips in the Source box
	//set for loop variables
	var tipboxCols = 12;
	var imgOn, k;
	var tipsCOn, i;
	var column = 0;
	var tipBoxLoc = 0;
   
	//loop thru cols to reset image selections
	for (k=0; k<tipboxCols; k++)
	{   
		//calc col array position
		tipsCOn = tips_ColOn[k];
		imgOn = tips_ImgOn[k];
		if(this[tipsCOn] == 1)
		{
			this[imgOn] = myRoot_YTip;
		}
		else
		{
			this[imgOn] = myRoot_NTip;
		}
	}

}

//set images for Dest Box based on the existing checkmarks
function setDestBoxImg_96()
{
	//trying loop thru img array instead
	//reset images for all the tips in the Source box
	//set for loop variables
	var tipboxCols = 12;
	var imgOff, k;
	var tipsCOff, i;
	var column = 0;
	var tipBoxLoc = 0;
   
	//loop thru cols to reset image selections
	for (k=0; k<tipboxCols; k++)
	{   
		//calc col array position
		tipsCOff = tips_ColOff[k];
		imgOff = tips_ImgOff[k];
		if(this[tipsCOff] == 1)
		{
			this[imgOff] = myRoot_YTip;
		}
		else
		{
			this[imgOff] = myRoot_NTip;
		}
	}

}
```
