## Supplementary material for "Automated chromatin profiling with spa-ChIP-seq uncovers the impacts of condition variations": Cao-etal-revised-2025-biorxiv-automation-files: Frm_Images.js

```
/***********************************
************************************
****** Set Default Variables  ******
************************************
***********************************/
//array count from the protocol file
var row = 0;
var column = 0;

//image locations
var myRoot =  masterFilePath + 'Forms/Supporting Files';

//new setup
var cols = myRoot + '/Plates/Columns/img_';
var pools = myRoot + '/Plates/Pools/img_';
var index = myRoot + '/Plates/Index/img_Index_';
var red_insert = myRoot + '/Plates/Inserts/insert_Red.png';
var red_plate = myRoot + '/Plates/Inserts/insert_Red_Plate.png';
var silver_insert = myRoot + '/Plates/Inserts/insert_Silver.png';
var silver_plate = myRoot + '/Plates/Inserts/insert_Silver_Plate.png';
var no_insert = myRoot + '/Plates/Inserts/insert_None.png';
var blank_insert = myRoot + '/Plates/Inserts/insert_Blank.png';
var res = myRoot + '/Plates/Res/img_';
var blank_plate = myRoot + '/Plates/Columns/img_Blank.png';
var empty_plate = myRoot + '/Plates/Columns/img_Empty.png';
var blank_key = myRoot + '/Plates/Columns/img_Blank.png';
var key = myRoot + '/Plates/Key/Img_Key_';
var mm = myRoot + '/Plates/MM/Img_MM_';
var mag = myRoot + '/Plates/Img_Mag.png';

/***********************************
************************************
******  Set Arrays from Form  ******
************************************
***********************************/
//***Images**\\
//set arrays for images
var bravo_Img = 
[
 "imgBravo_1",
 "imgBravo_2",
 "imgBravo_3",
 "imgBravo_4",
 "imgBravo_5",
 "imgBravo_6",
 "imgBravo_7",
 "imgBravo_8",
 "imgBravo_9"
 ]
 
 var key_Img = 
[
 "imgKey"
]

  //inserts
 var insert_Img = 
[
 "imgInsert_1",
 "imgInsert_2",
 "imgInsert_3",
 "imgInsert_4",
 "imgInsert_5",
 "imgInsert_6",
 "imgInsert_7",
 "imgInsert_8",
 "imgInsert_9"
 ]
 
var ThermoCube =
[
"frm_ThermoCube"
]

//labware name array
var bravo_Labware =
[
 "bravo_deck_1",
 "bravo_deck_2",
 "bravo_deck_3",
 "bravo_deck_4",
 "bravo_deck_5",
 "bravo_deck_6",
 "bravo_deck_7",
 "bravo_deck_8",
 "bravo_deck_9"
]
/***********************************
************************************
********* Set Bravo Images **********
************************************
***********************************/

//clear the form
function empty_BravoDeck()
{
	this["imgKey"] = blank;
	for (var i = 0 ; i <=9; i++) 
	{
		this["imgBravo_"+i] = blank;
		this["bravo_deck_"+i] = "";
	}
	//print to log
	print("bravo img grid was reset to blank.")
	//clear the status bar
	var update_status ="";
}	
var deck_pos = 9;
var color = "Blue";
//switch to Water plate for Elution
function update_BravoDeck(deck_pos, color)
{
	this['imgInsert_'+deck_pos] = no_insert;
	this['imgBravo_'+deck_pos] = res + color + '.png';
}	
```
