## Supplementary material for "Automated chromatin profiling with spa-ChIP-seq uncovers the impacts of condition variations": Cao-etal-revised-2025-biorxiv-automation-files: Frm_Push_Btns.js

```
/***********************************
************************************
*******scripts for all buttons******
************************************
***********************************/


//Check:
function btnCheckWorkspace()
{
	// Show deck layout and additional information
	open(masterFilePath + 'Forms/Scripts/Set_SS_WS_State_Function.js');
	running = 0
	//display the workspace text
	set_initial_state(frm_protocol_selected, columns_samples);

	//re-set tip box images
	open(masterFilePath + 'Forms/Scripts/96LT_Tip_Images_for_Form.js');
	running = 0
	//run the tip box functions
	setSourceBoxImg_96();
	setDestBoxImg_96();

	//update images
	open(masterFilePath + 'Forms/Scripts/Frm_Images.js');
	running = 0
	//run img fuction
	set_BravoDeck_Img(columns_samples)

}
//Run:
function btnRunProtocol()
{
	// get current_protocol_selected
	if(typeof(protocol_selected) == "undefined")
	{
		var protocol_selected = ""
	}

	var current_protocol_selected = protocol_selected

	// Show deck layout and additional information
	open(masterFilePath + 'Forms/Scripts/Set_SS_WS_State_Function.js');

	if(current_protocol_selected == protocol_selected)
	{
		running = 1
		print("Run: " + form_protocol_selected)
		runset.appendProtocolFileToRunset (form_protocol_selected, "1", "", frmName)
	}
	else
	{
		print("Skip Run as displayed layout incorrect at time of press")
	}
}
//Reset the form to defaults:
function btnResetDefaults_96LT()
{
	//reset the form selections
	var form_protocol = "";
	var columns_samples = 5;

	//reset tip selections
	for(i=1;i<13;i++)
	{
		eval("box2_"+i+" = 0");
		eval("box8_"+i+" = 0");
	}

	//reset Bravo layout
	for(i=1;i<10;i++)
	{
		eval("bravo_deck_"+i+" = 0");
	}

	var temp_4 = "";
	var temp_6 = "";
	var temp_9 = "";
}

//Update tip boxes:
function btnUpdateTipBoxes_96LT()
{
	//re-set images
	//open the file with the functions
	var scriptPath =  masterFilePath + 'Forms/Scripts/'
	//open script to acces functions in that script
	open(scriptPath + "96LT_Tip_Images_for_Form.js");

	//run the rest function
	setSourceBoxImg_96();
	setDestBoxImg_96();
}

//Reset tip boxes:
function btnResetTipBoxes_96LT()
{
	// reset current tip state
	// full box on position 2
	// empty box on position 8

	if(typeof(running)=="undefined")
	{
		for(x = 1;x<13;x++)
		{
			eval("box2_"+x+"=1")
			eval("box8_"+x+"=0")
		}
	print("Tip box count reset")
	}
	else
	{
		if(running==0)
		{
			for(x = 1;x<13;x++)
			{
				eval("box2_"+x+"=1")
				eval("box8_"+x+"=0")
			}
		print("Tip box count reset")
		}
	}

	//re-set images
	//open the file with the functions
	var scriptPath =  masterFilePath + 'Forms/Scripts/'
	//open script to acces functions in that script
	open(scriptPath + '96LT_Tip_Images_for_Form.js');

	//run the rest function
	setSourceBoxImg_96();
	setDestBoxImg_96();
}

//Clear tip box:
function btnClearTipBoxes_96LT()
{
	// reset current tip state
	// full box on position 2
	// empty box on position 8

	if(typeof(running)=="undefined")
	{
		for(x = 1;x<13;x++)
		{
			eval("box2_"+x+"=0")
			eval("box8_"+x+"=0")
		}
	print("Tip box count reset")
	}
	else
	{
	if(running==0)
		{
			for(x = 1;x<13;x++)
			{
				eval("box2_"+x+"=0")
				eval("box8_"+x+"=0")
			}
		print("Tip box count reset")
		}
	}

	//re-set images
	//open the file with the functions
	var scriptPath = masterFilePath + 'Forms/Scripts/';
	
	//open script to acces functions in that script
	open(scriptPath + '96LT_Tip_Images_for_Form.js');
		
	//run the rest function
	setSourceBoxImg_96();
	setDestBoxImg_96();
}
```
