## Supplementary material for "Automated chromatin profiling with spa-ChIP-seq uncovers the impacts of condition variations": Cao-etal-revised-2025-biorxiv-automation-files: run_form_popup.js

```
// Run a batch file located on the root of drive C: to display a file.
// Batch file is created new each time.
// File must be assiciated with a suitable program to display it.

// create batch file to display popup file
// if pdf view in Acrobat reader, for others just use image viewer
// replace AcroRd32.exe with Acrobat.exe if version  older than 10 is installed

popfile = new File()
popfile.Open("c:\\popup.bat","w")
filetype = popup_file.split(".")[1]
if(filetype=="pdf"||filetype=="PDF")
	{
	popfile.Write("@ECHO OFF\nCD %1\nSTART AcroRd32.exe %2\nEXIT\n")
	}
else
	{
	popfile.Write("@ECHO OFF\nCD %1\n%2\nEXIT\n")
	}
popfile.Close()

// run batch file with the popup_file supplied by the button script

dir = 'C:\\"VWorks Workspace"\\"NGS Option A"\\"illumina DNA Prep with Enrichment_v.A1.0.2"\\"Forms"'\\"Supporting Files"


program = 'C:\\popup.bat'


command = program +' '+dir+' "'+popup_file+'"'


run(command,1)

// delete the batch file

popfile.Delete("c:\\popup.bat")
```
