## Supplementary material for "Automated chromatin profiling with spa-ChIP-seq uncovers the impacts of condition variations": Cao-etal-revised-2025-biorxiv-automation-files: Set_SS_WS_State_Imgs_Function.js

```
// set protocol directory
var protocol_directory = masterFilePath + 'Protocol Files/';

// Set defaults
var form_status = "";

var protocol_selected = "none"

var form_column_required = ""

var i = 0;
/***********************************
************************************
****** Set Default Variables  ******
************************************
***********************************/
//array count from the protocol file
var row = 0;
var column = 0;

//image locations
var myRoot =  masterFilePath + 'Forms/Supporting Files';

//new setup
var cols = myRoot + '/Plates/Columns/img_';
var pools = myRoot + '/Plates/Pools/img_';
var index = myRoot + '/Plates/Index/img_Index_';
var red_insert = myRoot + '/Plates/Inserts/insert_Red.png';
var red_plate = myRoot + '/Plates/Inserts/insert_Red_Plate.png';
var silver_insert = myRoot + '/Plates/Inserts/insert_Silver.png';
var silver_plate = myRoot + '/Plates/Inserts/insert_Silver_Plate.png';
var no_insert = myRoot + '/Plates/Inserts/insert_None.png';
var blank_insert = myRoot + '/Plates/Inserts/insert_Blank.png';
var res = myRoot + '/Plates/Res/img_';
var blank_plate = myRoot + '/Plates/Columns/img_Blank.png';
var empty_plate = myRoot + '/Plates/Columns/img_Empty.png';
var blank_key = myRoot + '/Plates/Columns/img_Blank.png';
var key = myRoot + '/Plates/Key/Img_Key_';
var mm = myRoot + '/Plates/MM/Img_MM_';
var mag = myRoot + '/Plates/Img_Mag.png';

/***********************************
************************************
******  Set Arrays from Form  ******
************************************
***********************************/
//***Images**\\
//set arrays for images
var bravo_Img = 
[
 "imgBravo_1",
 "imgBravo_2",
 "imgBravo_3",
 "imgBravo_4",
 "imgBravo_5",
 "imgBravo_6",
 "imgBravo_7",
 "imgBravo_8",
 "imgBravo_9"
 ]
 
 var key_Img = 
[
 "imgKey"
]

  //inserts
 var insert_Img = 
[
 "imgInsert_1",
 "imgInsert_2",
 "imgInsert_3",
 "imgInsert_4",
 "imgInsert_5",
 "imgInsert_6",
 "imgInsert_7",
 "imgInsert_8",
 "imgInsert_9"
 ]
 
var ThermoCube =
[
"frm_ThermoCube"
]

//labware name array
var bravo_Labware =
[
 "bravo_deck_1",
 "bravo_deck_2",
 "bravo_deck_3",
 "bravo_deck_4",
 "bravo_deck_5",
 "bravo_deck_6",
 "bravo_deck_7",
 "bravo_deck_8",
 "bravo_deck_9"
]
/***********************************
****Create Insert Images on Form****
******for Dev File Selelected*******
***********************************/
//new fuction to set the inserts based off the selected .dev file
function devInserts_BravoDeck(devFile)
{
	if(devFile == 'Bravo_round_magnet_RedInsert.dev')
	{
		//red inserts at 4 & 6
		for (var i = 0 ; i <=9; i++) 
		{
			if(i == 4)
			{
				this["imgInsert_"+i] = red_insert;
			}
			else 
			{
			this["imgInsert_"+i] = blank_insert;
			}
			print("Based on dev file... Bravo Insert"+i+" file path = "+this["imgInsert_"+i]);
		}
	}
	if(devFile == 'Bravo_round_magnet_wash.dev')
	{
		//silver insert at 4
		for (var i = 0 ; i <=9; i++) 
		{			
			if(i == 4)
			{
				this["imgInsert_"+i] = red_insert;
			}
			if(i == 6)
			{
				this["imgInsert_"+i] = silver_insert;
			}
			else
			{
				this["imgInsert_"+i] = blank_insert;
			}
			print("Based on dev file... Bravo Insert"+i+" file path = "+this["imgInsert_"+i]);
		}
	}
	if(devFile == 'Bravo_round_magnet.dev')
	{
		//no inserts 
		for (var i = 0 ; i <=9; i++) 
		{
			this["imgInsert_"+i] = blank_insert;
			print("Based on dev file... Bravo Insert"+i+" file path = "+this["imgInsert_"+i]);
		}
	}
}

/***********************************
************************************
********* Set Bravo Images **********
************************************
***********************************/

//clear the form
function empty_BravoDeck()
{
	this["imgKey"] = blank_key;
	this["frm_ThermoCube"] = ""
	for (var i = 1 ; i <=9; i++) 
	{
		this["imgInsert_"+i] = blank_insert;
		this["imgBravo_"+i] = blank_plate;
		this["bravo_deck_"+i] = "";
	}
	//print to log
	print("bravo img grid was reset to blank.")
	//clear the status bar
	var update_status ="";
}	


function set_workspace_labware_imgs(functionProtocolName, function_columns_samples, frm_indexCol)
{
	print("starting deck layout");
	print("functionProtocol Name : " +functionProtocolName);
	switch(functionProtocolName)
	{
		// **************************************First**************************************
		// ************************************Protocol************************************ 
		case "1a_Input_Prep_vA1.0.1.pro":
		{
			//clear deck
			empty_BravoDeck();
			
			//set deck based on dev file
			var devFile = 'Bravo_round_magnet_RedInsert.dev';
			devInserts_BravoDeck(devFile);
			
			var protocol_selected = functionProtocolName
			this["frm_ThermoCube"] = "";
			
			//img key
			this["imgKey"] = key +'01a_' + columns_samples + '.png';
			print("imgKey file path= "+this["imgKey"]);
	
			for(i=1; i<=9; i++)
			{
				//set the img based on the num of columns
				if(i == 1)
				{
					//img
					this["imgInsert_"+i] = no_insert;
					this["imgBravo_"+i] = cols + 'Empty.png';
					print("Bravo "+i+" file path = "+this["imgBravo_"+i]);
					//txt
					if(plate_type_waste == "96 Deep Well PP Sqr Well V Btm")
					{
						this["bravo_deck_"+i] = "Waste: Empty tip box WITHOUT colored top";
					}
					else
					{
						this["bravo_deck_"+i] = 'Waste: ' + plate_type_waste;
					}
				}
				if(i == 2)
				{
					//tip box pos
				}
					
				if(i == 3)
				{
					//img
					this["imgInsert_"+i] = no_insert;
					print("imgInsert_"+i+" file path = "+this["imgInsert_"+i]);
					this["imgBravo_"+i] = res + 'LightBlue.png';
					print("Bravo "+i+" file path = "+this["imgBravo_"+i]);
					//txt
					this["bravo_deck_"+i] = 'Water: ' + plate_type_res
				}		
				if(i == 4)
				{
					//img insert
					this["imgInsert_"+i] = red_insert;
					print("imgInsert_"+i+" file path = "+this["imgInsert_"+i]);

					//img Bravo
					this["imgBravo_"+i] = empty_plate;
					print("Bravo "+i+" file path = "+this["imgBravo_"+i]);
					//txt
					if(devFile = "Bravo_round_magnet_RedInsert.dev")
					{
						this["bravo_deck_"+i] = 'Empty Plate: ' + plate_type_cleanup_pcr + ' in Red Alum Insert';
					}
					else
					{
						this["bravo_deck_"+i] = 'Empty Plate: ' + plate_type_cleanup_pcr;
					}
				}
				if(i == 5)
				{
					//img
					this["imgInsert_"+i] = no_insert;
					print("imgInsert_"+i+" file path = "+this["imgInsert_"+i]);
					this["imgBravo_"+i] = cols + 'Pink_' + columns_samples + '.png';
					print("Bravo "+i+" file path = "+this["imgBravo_"+i]);
					//txt
					this["bravo_deck_"+i] = 'PIXUL Shearing Plate: ' + plate_type_shear;
				}
				if(i == 6)
				{
					//img insert
					this["imgInsert_"+i] = no_insert;
					print("imgInsert_"+i+" file path = "+this["imgInsert_"+i]);
					this["imgBravo_"+i] = cols + 'BrightBlue_1.png';
					print("Bravo "+i+" file path = "+this["imgBravo_"+i]);
					//txt
					if(columns_samples > 3)
					{
						this["bravo_deck_"+i] = 'Rxn Mix Plate: ' + plate_type_mm_dw;
					}
					else
					{
						this["bravo_deck_"+i] = 'Rxn Mix Plate: ' + plate_type_mm_pcr;
					}
				}
				if(i == 7)
				{
					//img
					this["imgBravo_"+i] = mag;
					print("Bravo "+i+" file path = "+this["imgBravo_"+i]);
					//txt
					this["bravo_deck_"+i] = "";
				}
				if(i == 8)
				{
					//tip box pos
				}
				if(i == 9)
				{
					//img insert
					this["imgInsert_"+i] = no_insert;
					print("imgInsert_"+i+" file path = "+this["imgInsert_"+i]);
					//img bravo deck
					this["imgBravo_"+i] = res + 'Yellow.png';
					print("Bravo "+i+" file path = "+this["imgBravo_"+i]);
					//txt
					this["bravo_deck_"+i] = '5M NaCl Res: ' + plate_type_res;
				}
			}
		break;
		}
		// **************************************Next**************************************
		// ************************************Protocol************************************ 
		case "1b_Input_PEG_vA1.0.1.pro":
		{
			//clear deck
			empty_BravoDeck();
			
			//set deck based on dev file
			var devFile = 'Bravo_round_magnet_RedInsert.dev';
			devInserts_BravoDeck(devFile);
			
			var protocol_selected = functionProtocolName
			this["frm_ThermoCube"] = "";
			
			//img key
			this["imgKey"] = key +'01b_' + columns_samples + '.png';
			print("imgKey file path= "+this["imgKey"]);
	
			for(i=1; i<=9; i++)
			{
				//set the img based on the num of columns
				if(i == 1)
				{
					//img
					this["imgInsert_"+i] = no_insert;
					this["imgBravo_"+i] = cols + 'Empty.png';
					print("Bravo "+i+" file path = "+this["imgBravo_"+i]);
					//txt
					if(plate_type_waste == "96 Deep Well PP Sqr Well V Btm")
					{
						this["bravo_deck_"+i] = "Waste: Empty tip box WITHOUT colored top";
					}
					else
					{
						this["bravo_deck_"+i] = 'Waste: ' + plate_type_waste;
					}
				}
				if(i == 2)
				{
					//tip box pos
				}
					
				if(i == 3)
				{
					//img
					this["imgInsert_"+i] = blank_insert;
					print("imgInsert_"+i+" file path = "+this["imgInsert_"+i]);
					this["imgBravo_"+i] = blank_plate;
					print("Bravo "+i+" file path = "+this["imgBravo_"+i]);
					//txt
					this["bravo_deck_"+i] = "";
				}		
				if(i == 4)
				{
					//img insert
					this["imgInsert_"+i] = red_insert;
					print("imgInsert_"+i+" file path = "+this["imgInsert_"+i]);

					//img Bravo
					this["imgBravo_"+i] = cols + 'Pink_' + columns_samples + '.png';
					print("Bravo "+i+" file path = "+this["imgBravo_"+i]);
					//txt
					if(devFile = "Bravo_round_magnet_RedInsert.dev")
					{
						this["bravo_deck_"+i] = 'Input Samples (previous protocol): ' + plate_type_cleanup_pcr + ' in Red Alum Insert';
					}
					else
					{
						this["bravo_deck_"+i] = 'Input Samples (previous protocol): ' + plate_type_cleanup_pcr
					}
				}
				if(i == 5)
				{
					//img insert
					this["imgInsert_"+i] = no_insert;
					print("imgInsert_"+i+" file path = "+this["imgInsert_"+i]);
					this["imgBravo_"+i] = cols + 'Brown_1.png';
					print("Bravo "+i+" file path = "+this["imgBravo_"+i]);
					//txt
					if(columns_samples < 9)
					{
						this["bravo_deck_"+i] = 'Beads, PEG, NaCl Plate: ' + plate_type_mm_dw;
					}
					if(columns_samples > 8)
					{
						this["bravo_deck_"+i] = 'Beads, PEG, NaCl Plate: ' + plate_type_mm_dw_2mL;
					}
					
				}
				if(i == 6)
				{
					//img
					this["imgInsert_"+i] = blank_insert;
					print("imgInsert_"+i+" file path = "+this["imgInsert_"+i]);
					this["imgBravo_"+i] = blank_plate;
					print("Bravo "+i+" file path = "+this["imgBravo_"+i]);
					//txt
					this["bravo_deck_"+i] = "";
				}
				if(i == 7)
				{
					//img
					this["imgBravo_"+i] = mag;
					print("Bravo "+i+" file path = "+this["imgBravo_"+i]);
					//txt
					this["bravo_deck_"+i] = "";
				}
				if(i == 8)
				{
					//tip box pos
				}
				if(i == 9)
				{
					//img
					this["imgInsert_"+i] = blank_insert;
					print("imgInsert_"+i+" file path = "+this["imgInsert_"+i]);
					this["imgBravo_"+i] = blank_plate;
					print("Bravo "+i+" file path = "+this["imgBravo_"+i]);
					//txt
					this["bravo_deck_"+i] = "";
				}
			}
		break;
		}
		// **************************************Next**************************************
		// ************************************Protocol************************************ 
		case "1c_Input_Speedbead_Cleanup_vA1.0.1.pro":
		{
			//clear deck
			empty_BravoDeck();
			
			//set deck based on dev file
			var devFile = 'Bravo_round_magnet_RedInsert.dev';
			devInserts_BravoDeck(devFile);
			
			var protocol_selected = functionProtocolName
			this["frm_ThermoCube"] = "";
			//img key
			this["imgKey"] = key +'01c.png';
			print("imgKey file path= "+this["imgKey"]);
	
			for(i=1; i<=9; i++)
			{
				//set the img based on the num of columns
				if(i == 1)
				{
					//img
					this["imgInsert_"+i] = no_insert;
					this["imgBravo_"+i] = cols + 'Empty.png';
					print("Bravo "+i+" file path = "+this["imgBravo_"+i]);
					//txt
					if(plate_type_waste == "96 Deep Well PP Sqr Well V Btm")
					{
						this["bravo_deck_"+i] = "Waste: Empty tip box WITHOUT colored top";
					}
					else
					{
						this["bravo_deck_"+i] = 'Waste: ' + plate_type_waste;
					}
				}
				if(i == 2)
				{
					//tip box pos
				}
				if(i == 3)
				{
					//img
					this["imgInsert_"+i] = no_insert;
					this["imgBravo_"+i] = res + 'LightBlue.png';
					print("Bravo "+i+" file path = "+this["imgBravo_"+i]);
					//txt
					this["bravo_deck_"+i] = 'Elution TT Buffer: ' + plate_type_res;
				}
				if(i == 4)
				{
					//img
					this["imgInsert_"+i] = red_insert;
					this["imgBravo_"+i] = cols + 'Brown_' + columns_samples + '.png';
					print("Bravo "+i+" file path = "+this["imgBravo_"+i]);
					//txt
					if(devFile = "Bravo_round_magnet_RedInsert.dev")
					{
						this["bravo_deck_"+i] = 'Input Samples (previous protocol): ' + plate_type_cleanup_pcr + " in Red Alum Insert";
					}
					else
					{
						this["bravo_deck_"+i] = 'Input Samples (previous protocol): ' + plate_type_cleanup_pcr
					}
				}
				if(i == 5)
				{
					//img
					this["imgInsert_"+i] = blank_insert;
					this["imgBravo_"+i] = blank_plate;
					print("Bravo "+i+" file path = "+this["imgBravo_"+i]);
					//txt
					this["bravo_deck_"+i] = "";
				}
				if(i == 6)
				{
					//img
					this["imgInsert_"+i] = red_insert;
					this["imgBravo_"+i] = empty_plate;
					print("Bravo "+i+" file path = "+this["imgBravo_"+i]);
					//txt
					this["bravo_deck_"+i] = 'Elution Colection: ' + plate_type_cleanup_pcr_red;
				}
				if(i == 7)
				{
					//img
					this["imgBravo_"+i] = mag;
					print("Bravo "+i+" file path = "+this["imgBravo_"+i]);
					//txt
					this["bravo_deck_"+i] = "";
				}
				if(i == 8)
				{
					//tip box pos
				}
				if(i == 9)
				{
					//img
					this["imgInsert_"+i] = no_insert;
					this["imgBravo_"+i] = res + 'Orange.png';
					print("Bravo "+i+" file path = "+this["imgBravo_"+i]);
					//txt
					this["bravo_deck_"+i] = 'EtOH: ' + plate_type_res;
				}
			}
		break;
		}
		// **************************************Next**************************************
		// ************************************Protocol************************************ 
		case "2_Chromatin_Beads_Antibody_On_Mag_vA1.0.1.pro":
		{
			//clear deck
			empty_BravoDeck();
			
			//set deck based on dev file
			var devFile = 'Bravo_round_magnet_RedInsert.dev';
			devInserts_BravoDeck(devFile);
			
			var protocol_selected = functionProtocolName
			this["frm_ThermoCube"] = "";
			
			//img key
			this["imgKey"] = key +'02.png';
			print("imgKey file path= "+this["imgKey"]);
	
			for(i=1; i<=9; i++)
			{
				//set the img based on the num of columns
				if(i == 1)
				{
					//img
					this["imgInsert_"+i] = no_insert;
					this["imgBravo_"+i] = cols + 'Empty.png';
					print("Bravo "+i+" file path = "+this["imgBravo_"+i]);
					//txt
					if(plate_type_waste == "96 Deep Well PP Sqr Well V Btm")
					{
						this["bravo_deck_"+i] = "Waste: Empty tip box WITHOUT colored top";
					}
					else
					{
						this["bravo_deck_"+i] = 'Waste: ' + plate_type_waste;
					}
				}
				if(i == 2)
				{
					//tip box pos
				}
					
				if(i == 3)
				{
					//img
					this["imgInsert_"+i] = blank_insert;
					print("imgInsert_"+i+" file path = "+this["imgInsert_"+i]);
					this["imgBravo_"+i] = blank_plate;
					print("Bravo "+i+" file path = "+this["imgBravo_"+i]);
					//txt
					this["bravo_deck_"+i] = "";
				}		
				if(i == 4)
				{
					//img insert
					this["imgInsert_"+i] = red_insert;
					print("imgInsert_"+i+" file path = "+this["imgInsert_"+i]);

					//img Bravo
					this["imgBravo_"+i] = red_plate;
					print("Bravo "+i+" file path = "+this["imgBravo_"+i]);
					//txt
					this["bravo_deck_"+i] = "Red Alum Insert";
				}
				if(i == 5)
				{
					//img insert
					this["imgInsert_"+i] = no_insert;
					print("imgInsert_"+i+" file path = "+this["imgInsert_"+i]);
					this["imgBravo_"+i] = cols + 'Brown_1.png';
					print("Bravo "+i+" file path = "+this["imgBravo_"+i]);
					//txt
					if(columns_samples > 1)
					{
						this["bravo_deck_"+i] = 'Washed Beads in Lysis Buffer: ' + plate_type_mm_dw;
					}
					else
					{
						this["bravo_deck_"+i] = 'Washed Beads in Lysis Buffer: ' + plate_type_mm_pcr;
					}
				}
				if(i == 6)
				{
					//img
					this["imgInsert_"+i] = red_insert;
					print("imgInsert_"+i+" file path = "+this["imgInsert_"+i]);
					this["imgBravo_"+i] = empty_plate;
					print("Bravo "+i+" file path = "+this["imgBravo_"+i]);
					//txt
					this["bravo_deck_"+i] = "Empty: " + plate_type_tubes;
				}
				if(i == 7)
				{
					//img
					this["imgBravo_"+i] = mag;
					print("Bravo "+i+" file path = "+this["imgBravo_"+i]);
					//txt
					this["bravo_deck_"+i] = "";
				}
				if(i == 8)
				{
					//tip box pos
				}
				if(i == 9)
				{
					//img
					this["imgInsert_"+i] = no_insert;
					print("imgInsert_"+i+" file path = "+this["imgInsert_"+i]);
					this["imgBravo_"+i] = cols + 'Pink_' + columns_samples + '.png';
					print("Bravo "+i+" file path = "+this["imgBravo_"+i]);
					//txt
					this["bravo_deck_"+i] = "Chromatin Lysate: " + plate_type_shear;
				}
			}
		break;
		}
		
		// **************************************Next**************************************
		// ************************************Protocol************************************  
		case "3_ChIP_Wash_Elution_Off_Mag_vA1.0.1.pro":
		{
			//clear deck
			empty_BravoDeck();
			
			//set deck based on dev file
			var devFile = 'Bravo_round_magnet_RedInsert.dev';
			devInserts_BravoDeck(devFile);
			
			var protocol_selected = functionProtocolName
			this["frm_ThermoCube"] = "";
			//img key
			this["imgKey"] = key +'03.png';
			print("imgKey file path= "+this["imgKey"]);
	
			for(i=0; i<=9; i++)
			{
				//set the img based on the num of columns
				if(i == 1)
				{
					//img
					this["imgInsert_"+i] = no_insert;
					this["imgBravo_"+i] = cols + 'Empty.png';
					print("Bravo "+i+" file path = "+this["imgBravo_"+i]);
					//txt
					if(plate_type_waste == "96 Deep Well PP Sqr Well V Btm")
					{
						this["bravo_deck_"+i] = "Waste: Empty tip box WITHOUT colored top";
					}
					else
					{
						this["bravo_deck_"+i] = 'Waste: ' + plate_type_waste;
					}
				}
				if(i == 2)
				{
					//tip box pos
				}
				if(i == 3)
				{
					//img
					this["imgInsert_"+i] = no_insert;
					this["imgBravo_"+i] = res + 'LightBlue.png';
					print("Bravo "+i+" file path = "+this["imgBravo_"+i]);
					//txt
					this["bravo_deck_"+i] = 'Elution Buffer: ' + plate_type_res;
				}
				if(i == 4)
				{
					//img
					this["imgInsert_"+i] = red_insert;
					this["imgBravo_"+i] = empty_plate;
					print("Bravo "+i+" file path = "+this["imgBravo_"+i]);
					//txt
					if(devFile = "Bravo_round_magnet_RedInsert.dev")
					{
						this["bravo_deck_"+i] = 'Empty: ' + plate_type_cleanup_pcr + ' in Red Alum Insert';
					}
					else
					{
						this["bravo_deck_"+i] = 'Empty: ' + plate_type_cleanup_pcr;
					}
				}
				if(i == 5)
				{
					//img
					this["imgInsert_"+i] = no_insert;
					this["imgBravo_"+i] = empty_plate;
					print("Bravo "+i+" file path = "+this["imgBravo_"+i]);
					//txt
					this["bravo_deck_"+i] = 'Flowthru Colection: ' + plate_type_elute;
				}
				if(i == 6)
				{
					//img
					this["imgInsert_"+i] = red_insert;
					this["imgBravo_"+i] = cols + 'Brown_' + columns_samples + '.png';
					print("Bravo "+i+" file path = "+this["imgBravo_"+i]);
					//txt
					this["bravo_deck_"+i] = 'Overnight Sample: ' + plate_type_tubes;
				}
				if(i == 7)
				{
					//img
					this["imgBravo_"+i] = mag;
					print("Bravo "+i+" file path = "+this["imgBravo_"+i]);
					//txt
					this["bravo_deck_"+i] = "";
				}
				if(i == 8)
				{
					//tip box pos
				}
				if(i == 9)
				{
					//img
					this["imgInsert_"+i] = no_insert;
					this["imgBravo_"+i] = res + 'Blue.png';
					print("Bravo "+i+" file path = "+this["imgBravo_"+i]);
					//txt
					this["bravo_deck_"+i] = 'Wash Buffer 1: ' + plate_type_res;
				}
			}
		break;
		}

		// **************************************Next**************************************
		// ************************************Protocol************************************
		case "4_End_Repair_MM_Dispense_vA1.0.1.pro":
		{
			//clear deck
			empty_BravoDeck();
			
			//set deck based on dev file
			var devFile = 'Bravo_round_magnet_RedInsert.dev';
			devInserts_BravoDeck(devFile);
			
			var protocol_selected = functionProtocolName
			this["frm_ThermoCube"] = "";
			//img key
			this["imgKey"] = key +'04_' + columns_samples + '.png';
			print("imgKey file path= "+this["imgKey"]);
	
			for(i=1; i<=9; i++)
			{
				//set the img based on the num of columns
				if(i == 1)
				{
					//img
					this["imgInsert_"+i] = no_insert;
					this["imgBravo_"+i] = cols + 'Empty.png';
					print("Bravo "+i+" file path = "+this["imgBravo_"+i]);
					//txt
					if(plate_type_waste == "96 Deep Well PP Sqr Well V Btm")
					{
						this["bravo_deck_"+i] = "Waste: Empty tip box WITHOUT colored top";
					}
					else
					{
						this["bravo_deck_"+i] = 'Waste: ' + plate_type_waste;
					}
				}
				if(i == 2)
				{
					//tip box pos
				}
					
				if(i == 3)
				{
					//img
					this["imgInsert_"+i] = blank_insert;
					print("imgInsert_"+i+" file path = "+this["imgInsert_"+i]);
					this["imgBravo_"+i] = blank_plate;
					print("Bravo "+i+" file path = "+this["imgBravo_"+i]);
					//txt
					this["bravo_deck_"+i] = "";
				}		
				if(i == 4)
				{
					//img insert
					this["imgInsert_"+i] = red_insert;
					print("imgInsert_"+i+" file path = "+this["imgInsert_"+i]);
					//img Bravo
					this["imgBravo_"+i] = cols + 'Brown_' + columns_samples + '.png';
					print("Bravo "+i+" file path = "+this["imgBravo_"+i]);
					//txt
					if(devFile = "Bravo_round_magnet_RedInsert.dev")
					{
						this["bravo_deck_"+i] = 'ChIP-Seq Sample + Control Plate (previous protocol): ' + plate_type_cleanup_pcr + ' in Red Alum Insert';
					}
					else
					{
						this["bravo_deck_"+i] = 'ChIP-Seq Sample + Control Plate (previous protocol): ' + plate_type_cleanup_pcr;
					}
				}
				if(i == 5)
				{
					//img
					this["imgInsert_"+i] = blank_insert;
					print("imgInsert_"+i+" file path = "+this["imgInsert_"+i]);
					this["imgBravo_"+i] = blank_plate;
					print("Bravo "+i+" file path = "+this["imgBravo_"+i]);
					//txt
					this["bravo_deck_"+i] = "";
				}
				if(i == 6)
				{
					//img insert
					this["imgInsert_"+i] = no_insert;
					print("imgInsert_"+i+" file path = "+this["imgInsert_"+i]);
					this["imgBravo_"+i] = cols + 'BrightBlue_1.png';
					print("Bravo "+i+" file path = "+this["imgBravo_"+i]);
					//txt
					this["bravo_deck_"+i] = 'End Repair Buffer Plate: ' + plate_type_mm_pcr;
				}
				if(i == 7)
				{
					//img
					this["imgBravo_"+i] = mag;
					print("Bravo "+i+" file path = "+this["imgBravo_"+i]);
					//txt
					this["bravo_deck_"+i] = "";
				}
				if(i == 8)
				{
					//tip box pos
				}
				if(i == 9)
				{
					//img insert
					this["imgInsert_"+i] = blank_insert;
					print("imgInsert_"+i+" file path = "+this["imgInsert_"+i]);
					//img bravo
					this["imgBravo_"+i] = blank_plate;
					print("Bravo "+i+" file path = "+this["imgBravo_"+i]);
					//txt
					this["bravo_deck_"+i] = "";
				}
			}
		break;
		}

		// **************************************Next**************************************
		// ************************************Protocol************************************
		case "5_Adapter_Ligation_MM_vA1.0.1.pro":
		{
			//clear deck
			empty_BravoDeck();
			
			//set deck based on dev file
			var devFile = 'Bravo_round_magnet_RedInsert.dev';
			devInserts_BravoDeck(devFile);
			
			var protocol_selected = functionProtocolName
			this["frm_ThermoCube"] = "";
			//img key
			this["imgKey"] = key +'05_' + columns_samples + '.png';
			print("imgKey file path= "+this["imgKey"]);
	
			for(i=1; i<=9; i++)
			{
				//set the img based on the num of columns
				if(i == 1)
				{
					//img
					this["imgInsert_"+i] = no_insert;
					this["imgBravo_"+i] = cols + 'Empty.png';
					print("Bravo "+i+" file path = "+this["imgBravo_"+i]);
					//txt
					if(plate_type_waste == "96 Deep Well PP Sqr Well V Btm")
					{
						this["bravo_deck_"+i] = "Waste: Empty tip box WITHOUT colored top";
					}
					else
					{
						this["bravo_deck_"+i] = 'Waste: ' + plate_type_waste;
					}
				}
				if(i == 2)
				{
					//tip box pos
				}
					
				if(i == 3)
				{
					//img
					this["imgInsert_"+i] = blank_insert;
					print("imgInsert_"+i+" file path = "+this["imgInsert_"+i]);
					this["imgBravo_"+i] = blank_plate;
					print("Bravo "+i+" file path = "+this["imgBravo_"+i]);
					//txt
					this["bravo_deck_"+i] = "";
				}		
				if(i == 4)
				{
					//img insert
					this["imgInsert_"+i] = red_insert;
					print("imgInsert_"+i+" file path = "+this["imgInsert_"+i]);
					//img Bravo
					this["imgBravo_"+i] = cols + 'Pink_' + columns_samples + '.png';
					print("Bravo "+i+" file path = "+this["imgBravo_"+i]);
					//txt
					if(devFile = "Bravo_round_magnet_RedInsert.dev")
					{
						this["bravo_deck_"+i] = 'ChIP Plate (previous protocol): ' + plate_type_cleanup_pcr + ' in Red Alum Insert';
					}
					else
					{
						this["bravo_deck_"+i] = 'ChIP Plate (previous protocol): ' + plate_type_cleanup_pcr;
					}			
				}
				if(i == 5)
				{
					//img
					this["imgInsert_"+i] = red_insert;
					this["imgBravo_"+i] = index + frm_indexCol + '.png';
					print("Bravo "+i+" file path = "+this["imgBravo_"+i]);
					//txt
					this["bravo_deck_"+i] = 'Index: ' + plate_type_index;
				}
				if(i == 6)
				{
					//img insert
					this["imgInsert_"+i] = no_insert;
					print("imgInsert_"+i+" file path = "+this["imgInsert_"+i]);
					this["imgBravo_"+i] = cols + 'Rose_1.png';
					print("Bravo "+i+" file path = "+this["imgBravo_"+i]);
					//txt
					if(columns_samples > 3)
					{
						this["bravo_deck_"+i] = 'Ligation MM Plate: ' + plate_type_mm_dw;
					}
					else
					{
						this["bravo_deck_"+i] = 'Ligation MM Plate: ' + plate_type_mm_pcr;
					}
				}
				if(i == 7)
				{
					//img
					this["imgBravo_"+i] = mag;
					print("Bravo "+i+" file path = "+this["imgBravo_"+i]);
					//txt
					this["bravo_deck_"+i] = "";
				}
				if(i == 8)
				{
					//tip box pos
				}
				if(i == 9)
				{
					//img insert
					this["imgInsert_"+i] = blank_insert;
					print("imgInsert_"+i+" file path = "+this["imgInsert_"+i]);
					//img bravo
					this["imgBravo_"+i] = blank_plate;
					print("Bravo "+i+" file path = "+this["imgBravo_"+i]);
					//txt
					this["bravo_deck_"+i] = "";
				}
			}
		break;
		}

		// **************************************Next**************************************
		// ************************************Protocol************************************
	
		case "6_Ligation_Stop_Solution_Disp_vA1.0.1.pro":
		{
			//clear deck
			empty_BravoDeck();
			
			//set deck based on dev file
			var devFile = 'Bravo_round_magnet_RedInsert.dev';
			devInserts_BravoDeck(devFile);
			
			var protocol_selected = functionProtocolName
			this["frm_ThermoCube"] = "";
			
			//img key
			this["imgKey"] = key +'06_' + columns_samples + '.png';
			print("imgKey file path= "+this["imgKey"]);
	
			for(i=1; i<=9; i++)
			{
				//set the img based on the num of columns
				if(i == 1)
				{
					//img
					this["imgInsert_"+i] = no_insert;
					this["imgBravo_"+i] = cols + 'Empty.png';
					print("Bravo "+i+" file path = "+this["imgBravo_"+i]);
					//txt
					if(plate_type_waste == "96 Deep Well PP Sqr Well V Btm")
					{
						this["bravo_deck_"+i] = "Waste: Empty tip box WITHOUT colored top";
					}
					else
					{
						this["bravo_deck_"+i] = 'Waste: ' + plate_type_waste;
					}
				}
				if(i == 2)
				{
					//tip box pos
				}
					
				if(i == 3)
				{
					//img
					this["imgInsert_"+i] = blank_insert;
					print("imgInsert_"+i+" file path = "+this["imgInsert_"+i]);
					this["imgBravo_"+i] = blank_plate;
					print("Bravo "+i+" file path = "+this["imgBravo_"+i]);
					//txt
					this["bravo_deck_"+i] = "";
				}		
				if(i == 4)
				{
					//img insert
					this["imgInsert_"+i] = red_insert;
					print("imgInsert_"+i+" file path = "+this["imgInsert_"+i]);

					//img Bravo
					this["imgBravo_"+i] = cols + 'Brown_' + columns_samples + '.png';
					print("Bravo "+i+" file path = "+this["imgBravo_"+i]);
					//txt
					if(devFile = "Bravo_round_magnet_RedInsert.dev")
					{
						this["bravo_deck_"+i] = 'ChIP Plate (previous protocol): ' + plate_type_cleanup_pcr + ' in Red Alum Insert';
					}
					else
					{
						this["bravo_deck_"+i] = 'ChIP Plate (previous protocol): ' + plate_type_cleanup_pcr;
					}
				}
				if(i == 5)
				{
					//img
					this["imgInsert_"+i] = blank_insert;
					print("imgInsert_"+i+" file path = "+this["imgInsert_"+i]);
					this["imgBravo_"+i] = blank_plate;
					print("Bravo "+i+" file path = "+this["imgBravo_"+i]);
					//txt
					this["bravo_deck_"+i] = "";
				}
				if(i == 6)
				{
					//img insert
					this["imgInsert_"+i] = no_insert;
					print("imgInsert_"+i+" file path = "+this["imgInsert_"+i]);
					this["imgBravo_"+i] = cols + 'BrightBlue_1.png';
					print("Bravo "+i+" file path = "+this["imgBravo_"+i]);
					//txt
					if(columns_samples > 3)
					{
						this["bravo_deck_"+i] = 'Stop Soln Plate: ' + plate_type_mm_dw;
					}
					else
					{
						this["bravo_deck_"+i] = 'Stop Soln Plate: ' + plate_type_mm_pcr;
					}
				}
				if(i == 7)
				{
					//img
					this["imgBravo_"+i] = mag;
					print("Bravo "+i+" file path = "+this["imgBravo_"+i]);
					//txt
					this["bravo_deck_"+i] = "";
				}
				if(i == 8)
				{
					//tip box pos
				}
				if(i == 9)
				{
					//img insert
					this["imgInsert_"+i] = no_insert;
					print("imgInsert_"+i+" file path = "+this["imgInsert_"+i]);
					this["imgBravo_"+i] = res + 'Yellow.png';
					print("Bravo "+i+" file path = "+this["imgBravo_"+i]);
					//txt
					this["bravo_deck_"+i] = '5M NaCl Res: ' + plate_type_res;
				}
			}
		break;
		}

		// **************************************Next**************************************
		// ************************************Protocol************************************
		case "7a_Adapter_Ligation_Cleanup_vA1.0.1.pro":
		{
			//clear deck
			empty_BravoDeck();
			
			//set deck based on dev file
			var devFile = 'Bravo_round_magnet_RedInsert.dev';
			devInserts_BravoDeck(devFile);
			
			var protocol_selected = functionProtocolName
			this["frm_ThermoCube"] = "";
			
			//img key
			this["imgKey"] = key +'07a_' + columns_samples + '.png';
			print("imgKey file path= "+this["imgKey"]);
	
			for(i=1; i<=9; i++)
			{
				//set the img based on the num of columns
				if(i == 1)
				{
					//img
					this["imgInsert_"+i] = no_insert;
					this["imgBravo_"+i] = cols + 'Empty.png';
					print("Bravo "+i+" file path = "+this["imgBravo_"+i]);
					//txt
					if(plate_type_waste == "96 Deep Well PP Sqr Well V Btm")
					{
						this["bravo_deck_"+i] = "Waste: Empty tip box WITHOUT colored top";
					}
					else
					{
						this["bravo_deck_"+i] = 'Waste: ' + plate_type_waste;
					}
				}
				if(i == 2)
				{
					//tip box pos
				}
					
				if(i == 3)
				{
					//img
					this["imgInsert_"+i] = blank_insert;
					print("imgInsert_"+i+" file path = "+this["imgInsert_"+i]);
					this["imgBravo_"+i] = blank_plate;
					print("Bravo "+i+" file path = "+this["imgBravo_"+i]);
					//txt
					this["bravo_deck_"+i] = "";
				}		
				if(i == 4)
				{
					//img insert
					this["imgInsert_"+i] = red_insert;
					print("imgInsert_"+i+" file path = "+this["imgInsert_"+i]);

					//img Bravo
					this["imgBravo_"+i] = cols + 'Brown_' + columns_samples + '.png';
					print("Bravo "+i+" file path = "+this["imgBravo_"+i]);
					//txt
					if(devFile = "Bravo_round_magnet_RedInsert.dev")
					{
						this["bravo_deck_"+i] = 'ChIP Plate (previous protocol): ' + plate_type_cleanup_pcr + ' in Red Alum Insert';
					}
					else
					{
						this["bravo_deck_"+i] = 'ChIP Plate (previous protocol): ' + plate_type_cleanup_pcr;
					}
				}
				if(i == 5)
				{
					//img insert
					this["imgInsert_"+i] = no_insert;
					print("imgInsert_"+i+" file path = "+this["imgInsert_"+i]);
					this["imgBravo_"+i] = cols + 'DarkBrown_1.png';
					print("Bravo "+i+" file path = "+this["imgBravo_"+i]);
					//txt
					if(columns_samples < 12)
					{
						this["bravo_deck_"+i] = 'Speedbead MM: ' + plate_type_mm_dw;
					}
					if(columns_samples == 12)
					{
						this["bravo_deck_"+i] = 'Speedbead MM: ' + plate_type_mm_dw_2mL;
					}
				}
				if(i == 6)
				{
					//img
					this["imgInsert_"+i] = red_insert;
					print("imgInsert_"+i+" file path = "+this["imgInsert_"+i]);
					this["imgBravo_"+i] = empty_plate;
					print("Bravo "+i+" file path = "+this["imgBravo_"+i]);
					//txt
					this["bravo_deck_"+i] = "Empty Cleanup Plate: " + plate_type_cleanup_pcr_red;
				}
				if(i == 7)
				{
					//img
					this["imgBravo_"+i] = mag;
					print("Bravo "+i+" file path = "+this["imgBravo_"+i]);
					//txt
					this["bravo_deck_"+i] = "";
				}
				if(i == 8)
				{
					//tip box pos
				}
				if(i == 9)
				{
					//img
					this["imgInsert_"+i] = blank_insert;
					print("imgInsert_"+i+" file path = "+this["imgInsert_"+i]);
					this["imgBravo_"+i] = blank_plate;
					print("Bravo "+i+" file path = "+this["imgBravo_"+i]);
					//txt
					this["bravo_deck_"+i] = "";
				}
			}
		break;
		}

		// **************************************Next**************************************
		// ************************************Protocol************************************
		case "7b_Adapter_Ligation_Cleanup_vA1.0.1.pro":
		{
			//clear deck
			empty_BravoDeck();
			
			//set deck based on dev file
			var devFile = 'Bravo_round_magnet_RedInsert.dev';
			devInserts_BravoDeck(devFile);
			
			var protocol_selected = functionProtocolName
			this["frm_ThermoCube"] = "";
			//img key
			this["imgKey"] = key +'07b.png';
			print("imgKey file path= "+this["imgKey"]);
	
			for(i=1; i<=9; i++)
			{
				//set the img based on the num of columns
				if(i == 1)
				{
					//img
					this["imgInsert_"+i] = no_insert;
					this["imgBravo_"+i] = cols + 'Empty.png';
					print("Bravo "+i+" file path = "+this["imgBravo_"+i]);
					//txt
					if(plate_type_waste == "96 Deep Well PP Sqr Well V Btm")
					{
						this["bravo_deck_"+i] = "Waste: Empty tip box WITHOUT colored top";
					}
					else
					{
						this["bravo_deck_"+i] = 'Waste: ' + plate_type_waste;
					}
				}
				if(i == 2)
				{
					//tip box pos
				}
				if(i == 3)
				{
					//img
					this["imgInsert_"+i] = no_insert;
					this["imgBravo_"+i] = res + 'LightBlue.png';
					print("Bravo "+i+" file path = "+this["imgBravo_"+i]);
					//txt
					this["bravo_deck_"+i] = 'Elution TT Buffer: ' + plate_type_res;
				}
				if(i == 4)
				{
					//img
					this["imgInsert_"+i] = red_insert;
					this["imgBravo_"+i] = cols + 'Brown_' + columns_samples + '.png';
					print("Bravo "+i+" file path = "+this["imgBravo_"+i]);
					//txt
					if(devFile = "Bravo_round_magnet_RedInsert.dev")
					{
						this["bravo_deck_"+i] = 'ChIP Plate (previous protocol): ' + plate_type_cleanup_pcr + ' in Red Alum Insert';
					}
					else
					{
						this["bravo_deck_"+i] = 'ChIP Plate (previous protocol): ' + plate_type_cleanup_pcr;
					}
				}
				if(i == 5)
				{
					//img
					this["imgInsert_"+i] = blank_insert;
					this["imgBravo_"+i] = blank_plate;
					print("Bravo "+i+" file path = "+this["imgBravo_"+i]);
					//txt
					this["bravo_deck_"+i] = "";
				}
				if(i == 6)
				{
					//img
					this["imgInsert_"+i] = red_insert;
					this["imgBravo_"+i] = empty_plate;
					print("Bravo "+i+" file path = "+this["imgBravo_"+i]);
					//txt
					this["bravo_deck_"+i] = 'Elution Colection: ' + plate_type_cleanup_pcr_red;
				}
				if(i == 7)
				{
					//img
					this["imgBravo_"+i] = mag;
					print("Bravo "+i+" file path = "+this["imgBravo_"+i]);
					//txt
					this["bravo_deck_"+i] = "";
				}
				if(i == 8)
				{
					//tip box pos
				}
				if(i == 9)
				{
					//img
					this["imgInsert_"+i] = no_insert;
					this["imgBravo_"+i] = res + 'Orange.png';
					print("Bravo "+i+" file path = "+this["imgBravo_"+i]);
					//txt
					this["bravo_deck_"+i] = 'EtOH: ' + plate_type_res;
				}
			}
		break;
		}

		// **************************************Next**************************************
		// ************************************Protocol************************************
		case "8_PCR_Enrichment_MM_Dispense_vA1.0.1.pro":
		{
			//clear deck
			empty_BravoDeck();
			
			//set deck based on dev file
			var devFile = 'Bravo_round_magnet_RedInsert.dev';
			devInserts_BravoDeck(devFile);
			
			var protocol_selected = functionProtocolName
			this["frm_ThermoCube"] = "";
			//img key
			this["imgKey"] = key +'08_' + columns_samples + '.png';
			print("imgKey file path= "+this["imgKey"]);
	
			for(i=1; i<=9; i++)
			{
				//set the img based on the num of columns
				if(i == 1)
				{
					//img
					this["imgInsert_"+i] = no_insert;
					this["imgBravo_"+i] = cols + 'Empty.png';
					print("Bravo "+i+" file path = "+this["imgBravo_"+i]);
					//txt
					if(plate_type_waste == "96 Deep Well PP Sqr Well V Btm")
					{
						this["bravo_deck_"+i] = "Waste: Empty tip box WITHOUT colored top";
					}
					else
					{
						this["bravo_deck_"+i] = 'Waste: ' + plate_type_waste;
					}
				}
				if(i == 2)
				{
					//tip box pos
				}
					
				if(i == 3)
				{
					//img
					this["imgInsert_"+i] = blank_insert;
					print("imgInsert_"+i+" file path = "+this["imgInsert_"+i]);
					this["imgBravo_"+i] = blank_plate;
					print("Bravo "+i+" file path = "+this["imgBravo_"+i]);
					//txt
					this["bravo_deck_"+i] = "";
				}		
				if(i == 4)
				{
					//img insert
					this["imgInsert_"+i] = red_insert;
					print("imgInsert_"+i+" file path = "+this["imgInsert_"+i]);
					//img Bravo
					this["imgBravo_"+i] = cols + 'Pink_' + columns_samples + '.png';
					print("Bravo "+i+" file path = "+this["imgBravo_"+i]);
					//txt
					if(devFile = "Bravo_round_magnet_RedInsert.dev")
					{
						this["bravo_deck_"+i] = 'Eluted samples (previous protocol): ' + plate_type_cleanup_pcr + ' in Red Alum Insert';
					}
					else
					{
						this["bravo_deck_"+i] = 'Eluted samples (previous protocol): ' + plate_type_cleanup_pcr;
					}
				}
				if(i == 5)
				{
					//img
					this["imgInsert_"+i] = blank_insert;
					this["imgBravo_"+i] = blank_plate;
					print("Bravo "+i+" file path = "+this["imgBravo_"+i]);
					//txt
					this["bravo_deck_"+i] = "";
				}
				if(i == 6)
				{
					//img insert
					this["imgInsert_"+i] = no_insert;
					print("imgInsert_"+i+" file path = "+this["imgInsert_"+i]);
					this["imgBravo_"+i] = cols + 'BrightBlue_1.png';
					print("Bravo "+i+" file path = "+this["imgBravo_"+i]);
					if(columns_samples > 5)
					{
						//txt
						this["bravo_deck_"+i] = 'PCR MM Plate: ' + plate_type_mm_dw;
					}
					else
					{
						//txt
						this["bravo_deck_"+i] = 'PCR MM Plate: ' + plate_type_mm_pcr;
					}
				}
				if(i == 7)
				{
					//img
					this["imgBravo_"+i] = mag;
					print("Bravo "+i+" file path = "+this["imgBravo_"+i]);
					//txt
					this["bravo_deck_"+i] = "";
				}
				if(i == 8)
				{
					//tip box pos
				}
				if(i == 9)
				{
					//img insert
					this["imgInsert_"+i] = blank_insert;
					print("imgInsert_"+i+" file path = "+this["imgInsert_"+i]);
					//img bravo
					this["imgBravo_"+i] = blank_plate;
					print("Bravo "+i+" file path = "+this["imgBravo_"+i]);
					//txt
					this["bravo_deck_"+i] = "";
				}
			}
		break;
		}

		// **************************************Next**************************************
		// ************************************Protocol************************************
		case "9a_PCR_Enrichment_Cleanup v2_vA1.0.1.pro":
		{
			//clear deck
			empty_BravoDeck();
			
			//set deck based on dev file
			var devFile = 'Bravo_round_magnet_RedInsert.dev';
			devInserts_BravoDeck(devFile);
			
			var protocol_selected = functionProtocolName
			this["frm_ThermoCube"] = "";
			
			//img key
			this["imgKey"] = key +'09a_' + columns_samples + '.png';
			print("imgKey file path= "+this["imgKey"]);
	
			for(i=1; i<=9; i++)
			{
				//set the img based on the num of columns
				if(i == 1)
				{
					//img
					this["imgInsert_"+i] = no_insert;
					this["imgBravo_"+i] = cols + 'Empty.png';
					print("Bravo "+i+" file path = "+this["imgBravo_"+i]);
					//txt
					if(plate_type_waste == "96 Deep Well PP Sqr Well V Btm")
					{
						this["bravo_deck_"+i] = "Waste: Empty tip box WITHOUT colored top";
					}
					else
					{
						this["bravo_deck_"+i] = 'Waste: ' + plate_type_waste;
					}
				}
				if(i == 2)
				{
					//tip box pos
				}
					
				if(i == 3)
				{
					//img
					this["imgInsert_"+i] = blank_insert;
					print("imgInsert_"+i+" file path = "+this["imgInsert_"+i]);
					this["imgBravo_"+i] = blank_plate;
					print("Bravo "+i+" file path = "+this["imgBravo_"+i]);
					//txt
					this["bravo_deck_"+i] = "";
				}		
				if(i == 4)
				{
					//img insert
					this["imgInsert_"+i] = red_insert;
					print("imgInsert_"+i+" file path = "+this["imgInsert_"+i]);

					//img Bravo
					this["imgBravo_"+i] = cols + 'Pink_' + columns_samples + '.png';
					print("Bravo "+i+" file path = "+this["imgBravo_"+i]);
					//txt
					if(devFile = "Bravo_round_magnet_RedInsert.dev")
					{
						this["bravo_deck_"+i] = 'Sample Plate (previous protocol): ' + plate_type_cleanup_pcr + ' in Red Alum Insert';
					}
					else
					{
					this["bravo_deck_"+i] = 'Sample Plate (previous protocol): ' + plate_type_cleanup_pcr;
					}
				}
				if(i == 5)
				{
					//img insert
					this["imgInsert_"+i] = no_insert;
					print("imgInsert_"+i+" file path = "+this["imgInsert_"+i]);
					this["imgBravo_"+i] = cols + 'DarkBrown_1.png';
					print("Bravo "+i+" file path = "+this["imgBravo_"+i]);
					//txt
					this["bravo_deck_"+i] = 'Speedbead MM: ' + plate_type_mm_dw;
				}
				if(i == 6)
				{
					//img
					this["imgInsert_"+i] = blank_insert;
					print("imgInsert_"+i+" file path = "+this["imgInsert_"+i]);
					this["imgBravo_"+i] = blank_plate;
					print("Bravo "+i+" file path = "+this["imgBravo_"+i]);
					//txt
					this["bravo_deck_"+i] = "";
				}
				if(i == 7)
				{
					//img
					this["imgBravo_"+i] = mag;
					print("Bravo "+i+" file path = "+this["imgBravo_"+i]);
					//txt
					this["bravo_deck_"+i] = "";
				}
				if(i == 8)
				{
					//tip box pos
				}
				if(i == 9)
				{
					//img
					this["imgInsert_"+i] = blank_insert;
					print("imgInsert_"+i+" file path = "+this["imgInsert_"+i]);
					this["imgBravo_"+i] = blank_plate;
					print("Bravo "+i+" file path = "+this["imgBravo_"+i]);
					//txt
					this["bravo_deck_"+i] = "";
				}
			}
		break;
		}

		// **************************************Next**************************************
		// ************************************Protocol************************************
		case "9b_PCR_Enrichment_Cleanup v2_vA1.0.1.pro":
		{
			//clear deck
			empty_BravoDeck();
			
			//set deck based on dev file
			var devFile = 'Bravo_round_magnet_RedInsert.dev';
			devInserts_BravoDeck(devFile);
			
			var protocol_selected = functionProtocolName
			this["frm_ThermoCube"] = "";
			//img key
			this["imgKey"] = key +'09b.png';
			print("imgKey file path= "+this["imgKey"]);
	
			for(i=1; i<=9; i++)
			{
				//set the img based on the num of columns
				if(i == 1)
				{
					//img
					this["imgInsert_"+i] = no_insert;
					this["imgBravo_"+i] = cols + 'Empty.png';
					print("Bravo "+i+" file path = "+this["imgBravo_"+i]);
					//txt
					if(plate_type_waste == "96 Deep Well PP Sqr Well V Btm")
					{
						this["bravo_deck_"+i] = "Waste: Empty tip box WITHOUT colored top";
					}
					else
					{
						this["bravo_deck_"+i] = 'Waste: ' + plate_type_waste;
					}
				}
				if(i == 2)
				{
					//tip box pos
				}
				if(i == 3)
				{
					//img
					this["imgInsert_"+i] = no_insert;
					this["imgBravo_"+i] = res + 'LightBlue.png';
					print("Bravo "+i+" file path = "+this["imgBravo_"+i]);
					//txt
					this["bravo_deck_"+i] = 'Elution TT Buffer: ' + plate_type_res;
				}
				if(i == 4)
				{
					//img
					this["imgInsert_"+i] = red_insert;
					this["imgBravo_"+i] = cols + 'Brown_' + columns_samples + '.png';
					print("Bravo "+i+" file path = "+this["imgBravo_"+i]);
					//txt
					if(devFile = "Bravo_round_magnet_RedInsert.dev")
					{
						this["bravo_deck_"+i] = 'Sample (previous protocol): ' + plate_type_cleanup_pcr + ' in Red Alum Insert';
					}
					else
					{
						this["bravo_deck_"+i] = 'Sample (previous protocol): ' + plate_type_cleanup_pcr;
					}						
				}
				if(i == 5)
				{
					//img
					this["imgInsert_"+i] = blank_insert;
					this["imgBravo_"+i] = blank_plate;
					print("Bravo "+i+" file path = "+this["imgBravo_"+i]);
					//txt
					this["bravo_deck_"+i] = "";
				}
				if(i == 6)
				{
					//add if statments for insert color
					//img
					this["imgInsert_"+i] = red_insert;
					this["imgBravo_"+i] = empty_plate;
					print("Bravo "+i+" file path = "+this["imgBravo_"+i]);
					//txt
					this["bravo_deck_"+i] = 'Elution Colection: ' + plate_type_final_elute;
				}
				if(i == 7)
				{
					//img
					this["imgBravo_"+i] = mag;
					print("Bravo "+i+" file path = "+this["imgBravo_"+i]);
					//txt
					this["bravo_deck_"+i] = "";
				}
				if(i == 8)
				{
					//tip box pos
				}
				if(i == 9)
				{
					//img
					this["imgInsert_"+i] = no_insert;
					this["imgBravo_"+i] = res + 'Orange.png';
					print("Bravo "+i+" file path = "+this["imgBravo_"+i]);
					//txt
					this["bravo_deck_"+i] = 'EtOH: ' + plate_type_res;
				}
			}
		break;
		}

		// **************************************Next**************************************
		// ************************************Protocol************************************
		case "":
		{
			var protocol_selected = functionProtocolName
			this["frm_ThermoCube"] = "";
			//img key
			this["imgKey"] = blank;
			print("imgKey file path= "+this["imgKey"]);
	
			for(i=0; i<=9; i++)
			{
				//img
				this["imgBravo_"+i] = blank;
				//txt
				this["bravo_deck_"+i] = "";
			}
		break;
		}
	}
}
set_initial_state(form_protocol, form_columns_samples)

// clean up form
if(form_column_required != "")
{
form_columns_samples = 1
}

if(protocol_selected == "none")
{
form_protocol_selected = ""
}
else
{
form_protocol_selected = protocol_directory+protocol_selected
}

print("finished layout update")

function set_protocol_selected(form_protocol)
{
	var protocol_selected = form_protocol;

	if(protocol_selected == "none")
	{
		form_protocol_selected = ""
	}
	else
	{
		form_protocol_selected = protocol_directory + protocol_selected
	}

	print("finished protocol selection")
}
```
