## Supplementary material for "Automated chromatin profiling with spa-ChIP-seq uncovers the impacts of condition variations": Cao-etal-revised-2025-biorxiv-automation-files: Set_SS_WS_State_Imgs_Function_PlateSetUp.js

```
// set protocol directory
var protocol_directory = masterFilePath + 'Protocol Files/';

// Set defaults
var form_status = "";

var protocol_selected = "none"

var form_column_required = ""

var i = 0;
/***********************************
************************************
****** Set Default Variables  ******
************************************
***********************************/
//array count from the protocol file
var row = 0;
var column = 0;

//image locations
var myRoot =  masterFilePath + 'Forms/Supporting Files';

//new setup
var cols = myRoot + '/Plates/Columns/img_';
var pools = myRoot + '/Plates/Pools/img_';
var index = myRoot + '/Plates/Index/img_Index_';
var red_insert = myRoot + '/Plates/Inserts/insert_Red.png';
var red_plate = myRoot + '/Plates/Inserts/insert_Red_Plate.png';
var silver_insert = myRoot + '/Plates/Inserts/insert_Silver.png';
var silver_plate = myRoot + '/Plates/Inserts/insert_Silver_Plate.png';
var no_insert = myRoot + '/Plates/Inserts/insert_None.png';
var blank_insert = myRoot + '/Plates/Inserts/insert_Blank.png';
var res = myRoot + '/Plates/Res/img_';
var blank_plate = myRoot + '/Plates/Columns/img_Blank.png';
var empty_plate = myRoot + '/Plates/Columns/img_Empty.png';
var blank_key = myRoot + '/Plates/Columns/img_Blank.png';
var key = myRoot + '/Plates/Key/Img_Key_';
var mm = myRoot + '/Plates/MM/Img_MM_';
var mag = myRoot + '/Plates/Img_Mag.png';
var arrow = myRoot + '/alert_arrow.png';
var blank = myRoot + '/alert_blank.png';

/***********************************
************************************
******  Set Arrays from Form  ******
************************************
***********************************/
//***Images**\\
//set arrays for images
var bravo_Img = 
[
 "imgBravo_1",
 "imgBravo_2",
 "imgBravo_3",
 "imgBravo_4",
 "imgBravo_5",
 "imgBravo_6",
 "imgBravo_7",
 "imgBravo_8",
 "imgBravo_9"
 ]
 
 var key_Img = 
[
 "imgKey"
]

  //inserts
 var insert_Img = 
[
 "imgInsert_1",
 "imgInsert_2",
 "imgInsert_3",
 "imgInsert_4",
 "imgInsert_5",
 "imgInsert_6",
 "imgInsert_7",
 "imgInsert_8",
 "imgInsert_9"
 ]
 
var ThermoCube =
[
"frm_ThermoCube"
]

//labware name array
var bravo_Labware =
[
 "bravo_deck_1",
 "bravo_deck_2",
 "bravo_deck_3",
 "bravo_deck_4",
 "bravo_deck_5",
 "bravo_deck_6",
 "bravo_deck_7",
 "bravo_deck_8",
 "bravo_deck_9"
]

var plateMap_Dest
[
"inputDest_1",
"inputDest_2",
"inputDest_3",
"inputDest_4",
"inputDest_5",
"inputDest_6",
"inputDest_7",
"inputDest_8",
"inputDest_9",
"inputDest_10",
"inputDest_11",
"inputDest_12"
]

var plateMap_Source
[
"inputSource_1",
"inputSource_2",
"inputSource_3",
"inputSource_4",
"inputSource_5",
"inputSource_6",
"inputSource_7",
"inputSource_8",
"inputSource_9",
"inputSource_10",
"inputSource_11",
"inputSource_12"
]

var arrow
[
"imgAlert_1",
"imgAlert_2",
"imgAlert_3",
"imgAlert_4",
"imgAlert_5",
"imgAlert_6",
"imgAlert_7",
"imgAlert_8",
"imgAlert_9",
"imgAlert_10",
"imgAlert_11",
"imgAlert_12"
]

/***********************************
****Create Insert Images on Form****
******for Dev File Selelected*******
***********************************/
//new fuction to set the inserts based off the selected .dev file
function devInserts_BravoDeck(devFile)
{
	if(devFile == 'Bravo_round_magnet_RedInsert.dev')
	{
		//red inserts at 4 & 6
		for (var i = 0 ; i <=9; i++) 
		{
			if(i == 4)
			{
				this["imgInsert_"+i] = red_insert;
			}
			else 
			{
			this["imgInsert_"+i] = blank_insert;
			}
			print("Based on dev file... Bravo Insert"+i+" file path = "+this["imgInsert_"+i]);
		}
	}
	if(devFile == 'Bravo_round_magnet_wash.dev')
	{
		//silver insert at 4
		for (var i = 0 ; i <=9; i++) 
		{			
			if(i == 4)
			{
				this["imgInsert_"+i] = red_insert;
			}
			if(i == 6)
			{
				this["imgInsert_"+i] = silver_insert;
			}
			else
			{
				this["imgInsert_"+i] = blank_insert;
			}
			print("Based on dev file... Bravo Insert"+i+" file path = "+this["imgInsert_"+i]);
		}
	}
	if(devFile == 'Bravo_round_magnet.dev')
	{
		//no inserts 
		for (var i = 0 ; i <=9; i++) 
		{
			this["imgInsert_"+i] = blank_insert;
			print("Based on dev file... Bravo Insert"+i+" file path = "+this["imgInsert_"+i]);
		}
	}
}

/***********************************
************************************
********* Set Bravo Images **********
************************************
***********************************/

//clear the form
function empty_BravoDeck()
{
	this["imgKey"] = blank_key;
	this["frm_ThermoCube"] = ""
	for (var i = 1 ; i <=9; i++) 
	{
		this["imgInsert_"+i] = blank_insert;
		this["imgBravo_"+i] = blank_plate;
		this["bravo_deck_"+i] = "";
	}
	//print to log
	print("bravo img grid was reset to blank.")
	//clear the status bar
	var update_status ="";
}	

function empty_PlateMap()
{
	for (var i = 1 ; i <=12; i++) 
	{
		this["inputSource_"+i] = "";
		this["inputDest_"+i] = "";
		this["imgAlert_"+i] = blank;
	}
}


function set_workspace_labware_imgs(columns_samples, columns_input)
{
	print("starting deck layout");
	{
		//clear deck
		empty_BravoDeck();
			
		//set deck based on dev file
		var devFile = 'Bravo_round_magnet_RedInsert.dev';
		devInserts_BravoDeck(devFile);
			
		this["frm_ThermoCube"] = "";
		//Bravo Deck
		for(i=1; i<=9; i++)
		{
			//set the img based on the num of columns
			if(i == 1)
			{
				//img
				this["imgInsert_"+i] = no_insert;
				this["imgBravo_"+i] = cols + 'Empty.png';
				print("Bravo "+i+" file path = "+this["imgBravo_"+i]);
				//txt
				this["bravo_deck_"+i] = 'Waste Plate (not used but can be left on the deck) ';
			}
			if(i == 2)
			{
				//tip box pos
			}
			if(i == 3)
			{
				//img
				this["imgInsert_"+i] = blank_insert;
				print("imgInsert_"+i+" file path = "+this["imgInsert_"+i]);
				this["imgBravo_"+i] = blank_plate;
				print("Bravo "+i+" file path = "+this["imgBravo_"+i]);
				//txt
				this["bravo_deck_"+i] = ""
			}			
			if(i == 4)
			{
				//img insert
				this["imgInsert_"+i] = red_insert;
				print("imgInsert_"+i+" file path = "+this["imgInsert_"+i]);

				//img Bravo
				this["imgBravo_"+i] = cols + 'Pink_' + columns_samples + '.png';
				print("Bravo "+i+" file path = "+this["imgBravo_"+i]);
				//txt
				if(devFile = "Bravo_round_magnet_RedInsert.dev")
				{
					this["bravo_deck_"+i] = 'ChIP-Seq Samples: ' + plate_type_cleanup_pcr + ' in Red Alum Insert';
				}
				else
				{
					this["bravo_deck_"+i] = 'ChIP-Seq Samples: ' + plate_type_cleanup_pcr;
				}
			}
			if(i == 5)
			{
				//img
				this["imgInsert_"+i] = red_insert;
				print("imgInsert_"+i+" file path = "+this["imgInsert_"+i]);
				this["imgBravo_"+i] = cols + 'Brown_' + columns_input + '.png';
				print("Bravo "+i+" file path = "+this["imgBravo_"+i]);
				//txt
				this["bravo_deck_"+i] = 'Input Controls: ' + plate_type_cleanup_pcr + '*img reflects number of columns to transfer, use plate map for locations';
			}
			if(i == 6)
			{
				//img insert
				this["imgInsert_"+i] = blank_insert;
				print("imgInsert_"+i+" file path = "+this["imgInsert_"+i]);
				this["imgBravo_"+i] = blank_plate;
				print("Bravo "+i+" file path = "+this["imgBravo_"+i]);
				//txt
				this["bravo_deck_"+i] = "";
			}
			if(i == 7)
			{
				//img
				this["imgBravo_"+i] = mag;
				print("Bravo "+i+" file path = "+this["imgBravo_"+i]);
				//txt
				this["bravo_deck_"+i] = "";
			}
			if(i == 8)
			{
				//tip box pos
			}
			if(i == 9)
			{
				//img insert
				this["imgInsert_"+i] = no_insert;
				print("imgInsert_"+i+" file path = "+this["imgInsert_"+i]);
				//img bravo deck
				this["imgBravo_"+i] = res + 'Green.png';
				print("Bravo "+i+" file path = "+this["imgBravo_"+i]);
				//txt
				this["bravo_deck_"+i] = 'Input Dilution TT Buffer: ' + plate_type_res;
			}
		}
		
		//Plate Map Destinations
		
		for(i=1; i<=12; i++)
		{
			if(columns_samples + i <= 12)
			{
				if(columns_input >= i)
				{
				this["inputDest_"+i] = columns_samples + i;
				this["imgAlert_"+i] = arrow;
				}
			}
			else
			{
				this["inputDest_"+i] = "";
				this["imgAlert_"+i] = blank;
			}
			print( "file path for imgAlert_" + i + " is: " + this["imgAlert_"+i]);
		}
		
	}
}

set_initial_state(form_protocol, form_columns_samples)

// clean up form
if(form_column_required != "")
{
form_columns_samples = 1
}

if(protocol_selected == "none")
{
form_protocol_selected = ""
}
else
{
form_protocol_selected = protocol_directory+protocol_selected
}

print("finished layout update")

function set_protocol_selected(form_protocol)
{
	var protocol_selected = form_protocol;

	if(protocol_selected == "none")
	{
		form_protocol_selected = ""
	}
	else
	{
		form_protocol_selected = protocol_directory + protocol_selected
	}

	print("finished protocol selection")
}
```
