## Supplementary material for "Automated chromatin profiling with spa-ChIP-seq uncovers the impacts of condition variations": Cao-etal-revised-2025-biorxiv-automation-files: background slide.pptx

### Slide 1
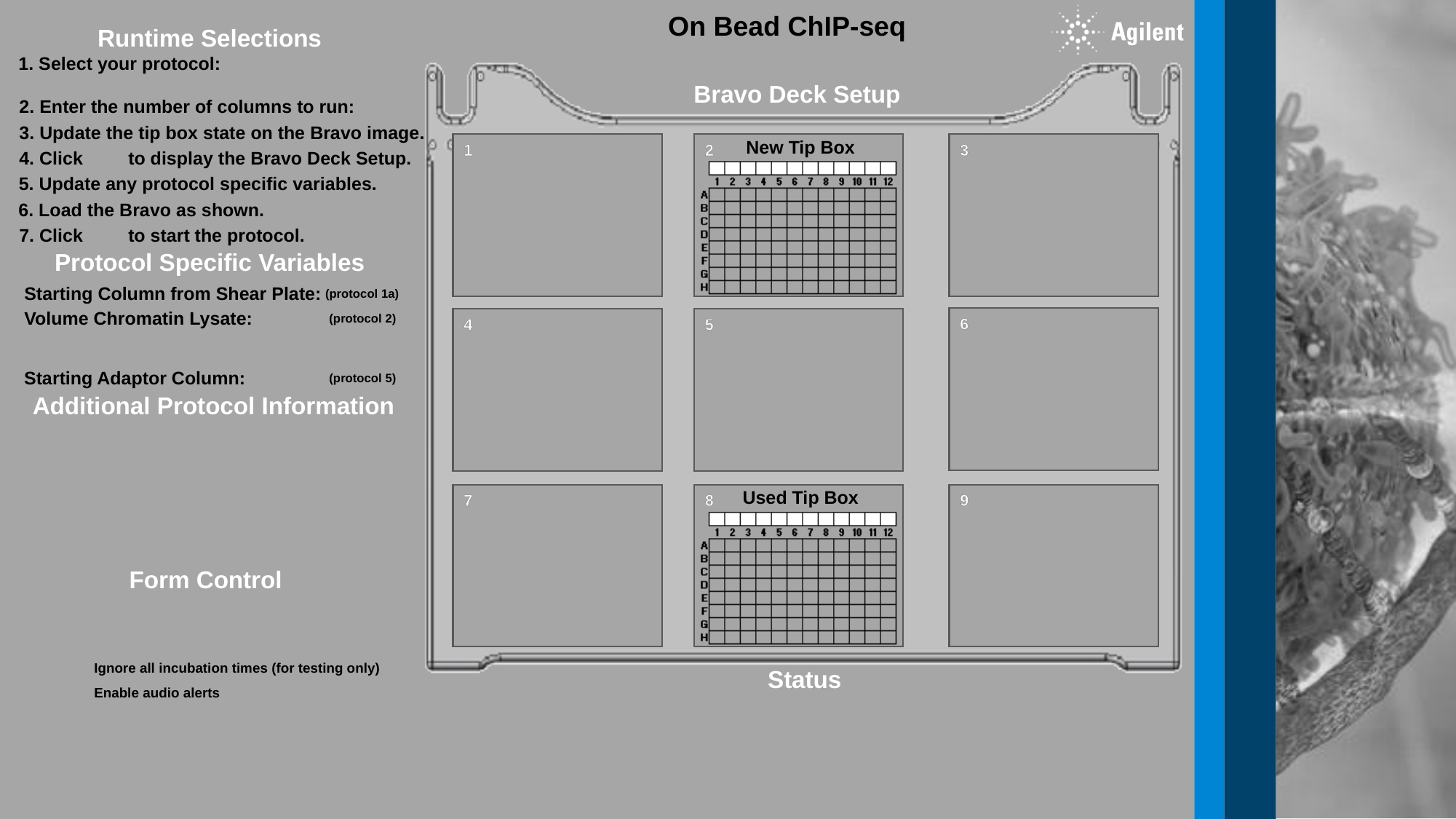

On Bead ChIP-seq
Runtime Selections
1. Select your protocol:
2. Enter the number of columns to run:
3. Update the tip box state on the Bravo image.
4. Click 	to display the Bravo Deck Setup.
5. Update any protocol specific variables.
6. Load the Bravo as shown.
7. Click 	to start the protocol.
1
3
4
7
9
Bravo Deck Setup
New Tip Box
2
Protocol Specific Variables
Starting Column from Shear Plate:
(protocol 1a)
Volume Chromatin Lysate:
(protocol 2)
Starting Adaptor Column:
(protocol 5)
6
5
Additional Protocol Information
Used Tip Box
8
Form Control
Ignore all incubation times (for testing only)
Enable audio alerts
Status

### Slide 2
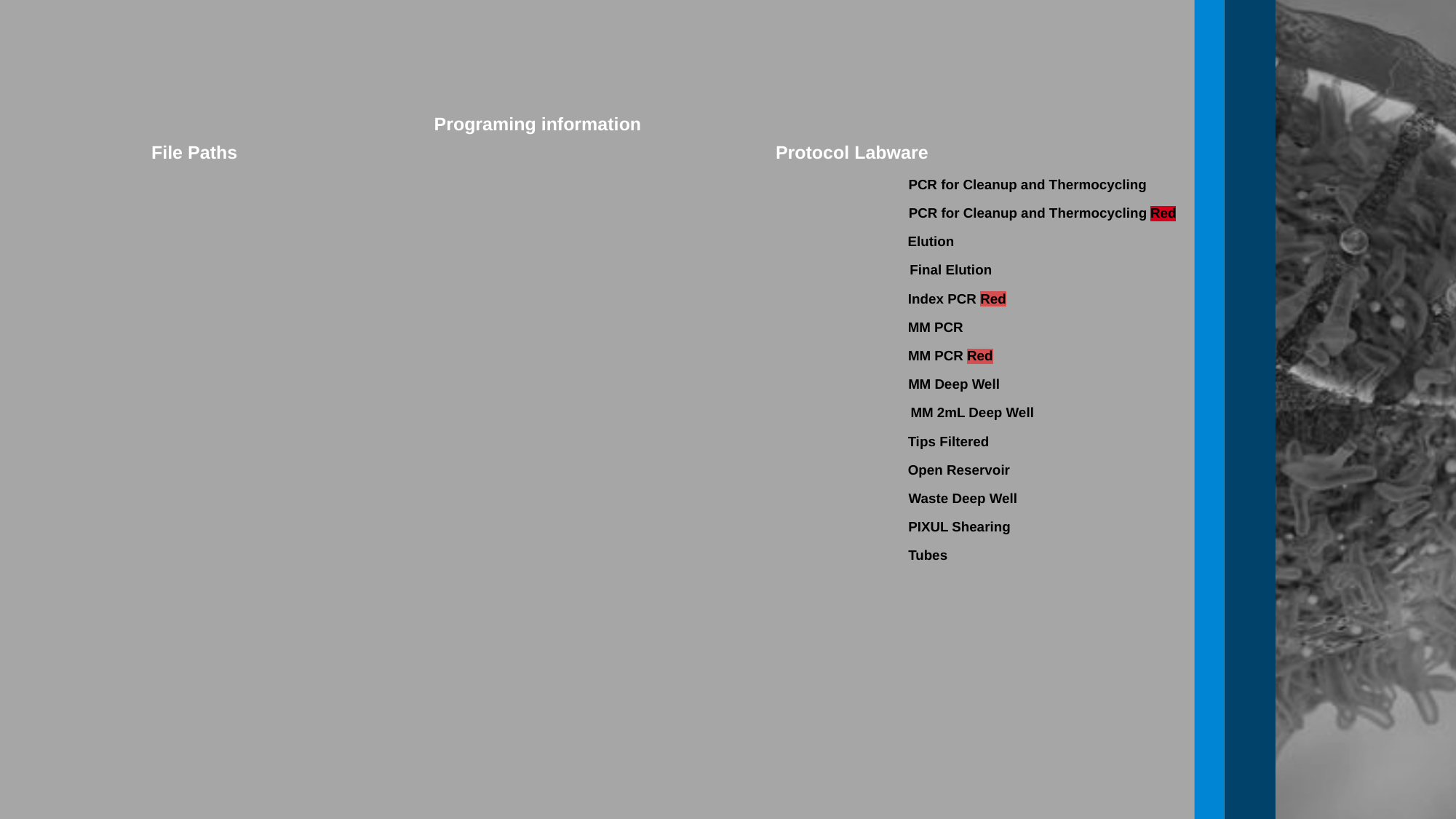

Programing information
File Paths
Protocol Labware
PCR for Cleanup and Thermocycling
PCR for Cleanup and Thermocycling Red
Elution
Final Elution
Index PCR Red
MM PCR
MM PCR Red
MM Deep Well
Tips Filtered
Open Reservoir
Waste Deep Well
PIXUL Shearing
Tubes
MM 2mL Deep Well

### Slide 3
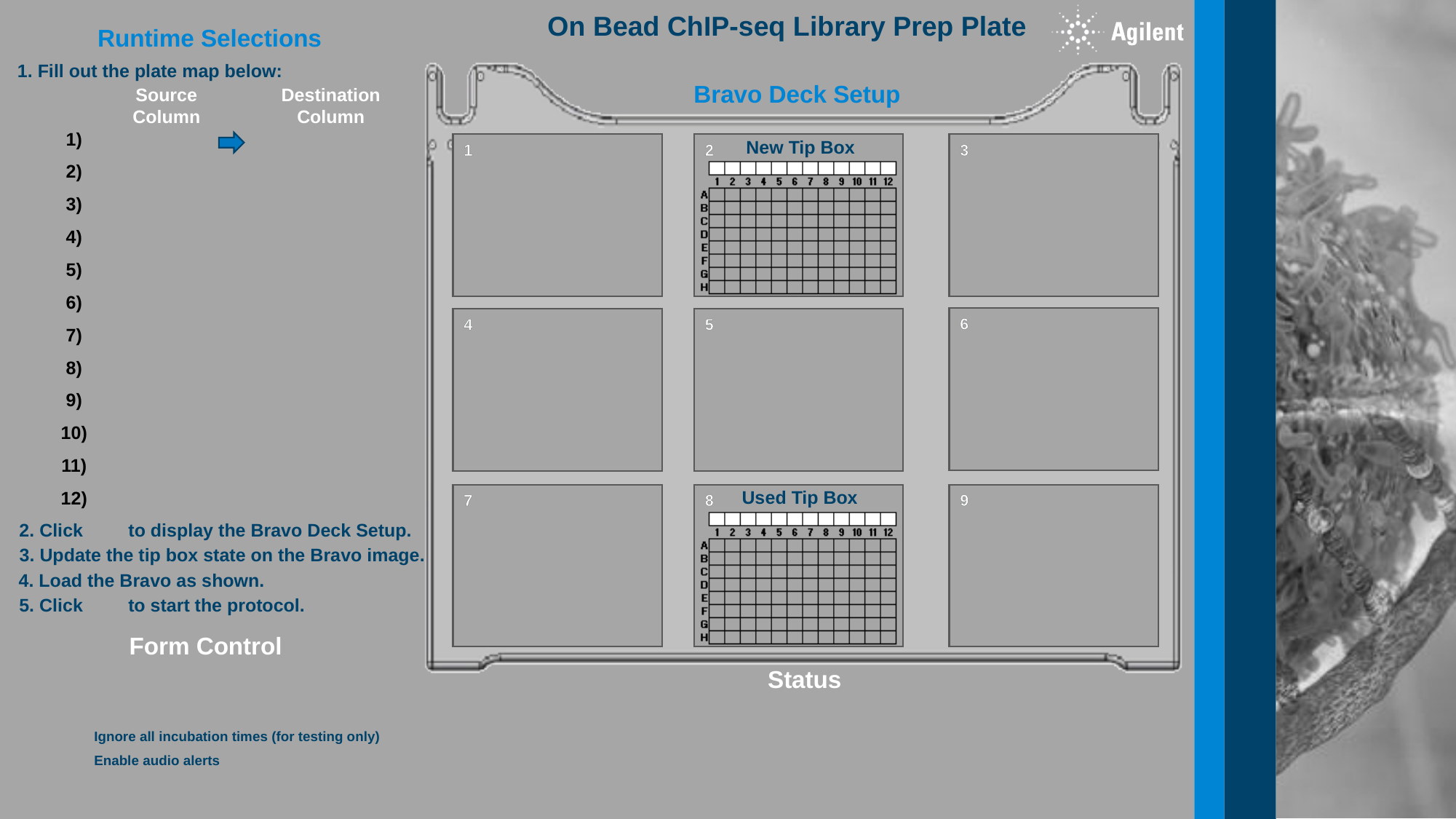

On Bead ChIP-seq Library Prep Plate
Runtime Selections
1. Fill out the plate map below:
1
3
4
7
9
Bravo Deck Setup
Source Column
Destination Column
1)
2)
3)
4)
5)
6)
7)
8)
9)
10)
11)
12)
New Tip Box
2
6
5
Used Tip Box
8
2. Click 	to display the Bravo Deck Setup.
3. Update the tip box state on the Bravo image.
4. Load the Bravo as shown.
5. Click 	to start the protocol.
Form Control
Status
Ignore all incubation times (for testing only)
Enable audio alerts

### Slide 4
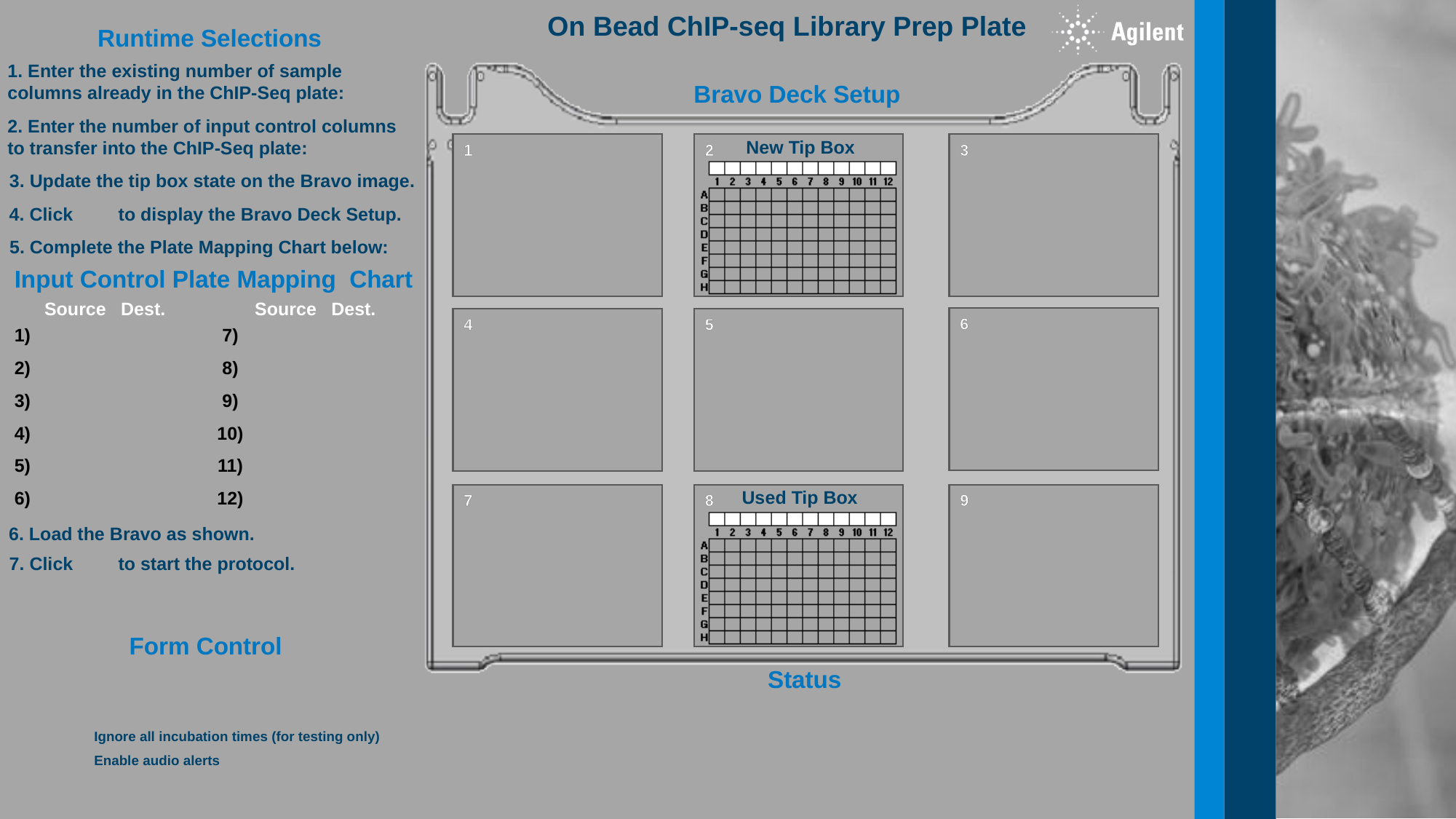

On Bead ChIP-seq Library Prep Plate
Runtime Selections
1. Enter the existing number of sample columns already in the ChIP-Seq plate:
2. Enter the number of input control columns to transfer into the ChIP-Seq plate:
3. Update the tip box state on the Bravo image.
4. Click 	to display the Bravo Deck Setup.
6. Load the Bravo as shown.
7. Click 	to start the protocol.
5. Complete the Plate Mapping Chart below:
1
3
4
7
9
Bravo Deck Setup
New Tip Box
2
Input Control Plate Mapping Chart
Source
Dest.
1)
2)
3)
4)
5)
6)
Source
Dest.
7)
8)
9)
10)
11)
12)
6
5
Used Tip Box
8
Form Control
Status
Ignore all incubation times (for testing only)
Enable audio alerts

### Slide 5
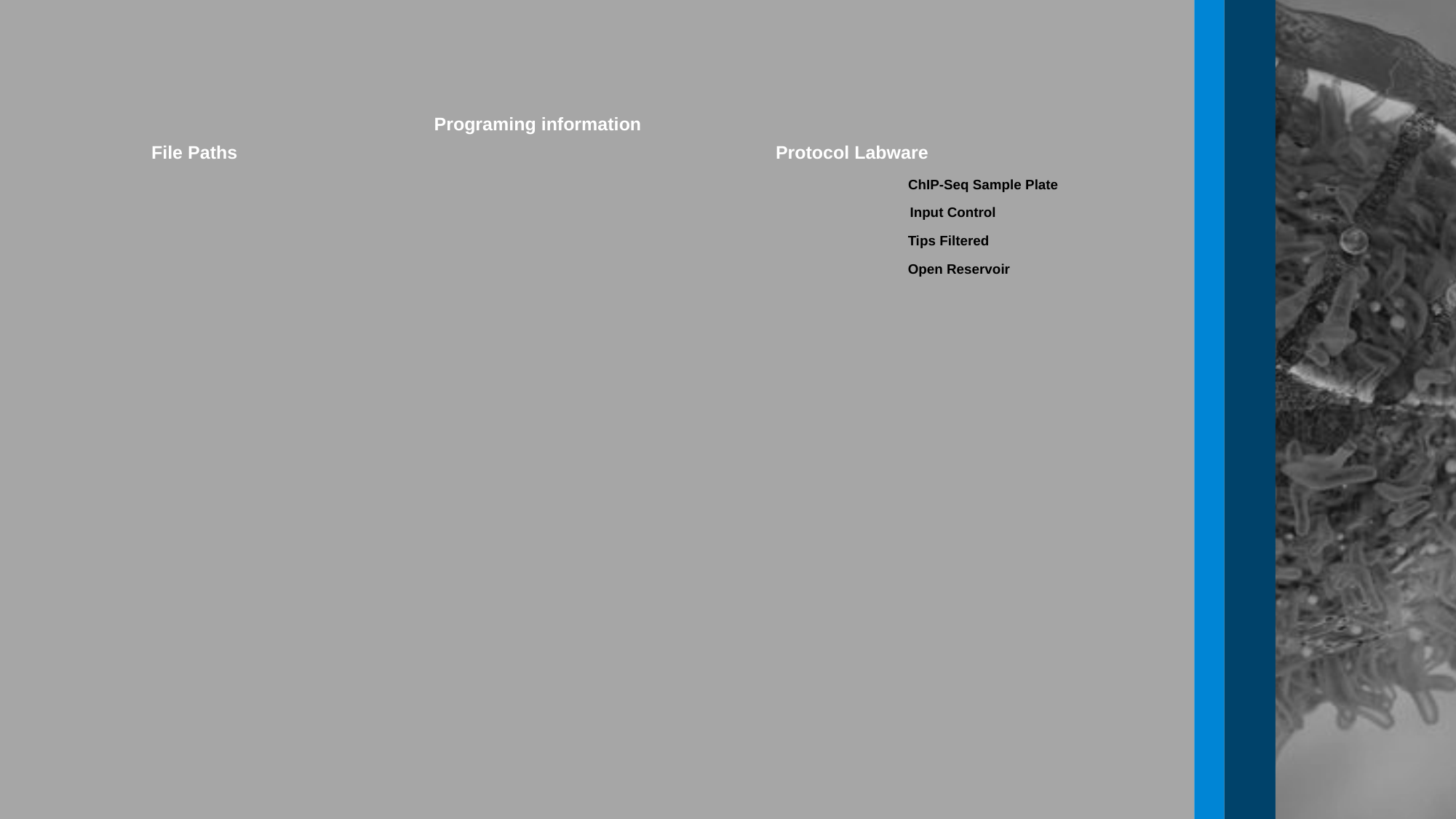

Programing information
File Paths
Protocol Labware
ChIP-Seq Sample Plate
Input Control
Tips Filtered
Open Reservoir

### Slide 6
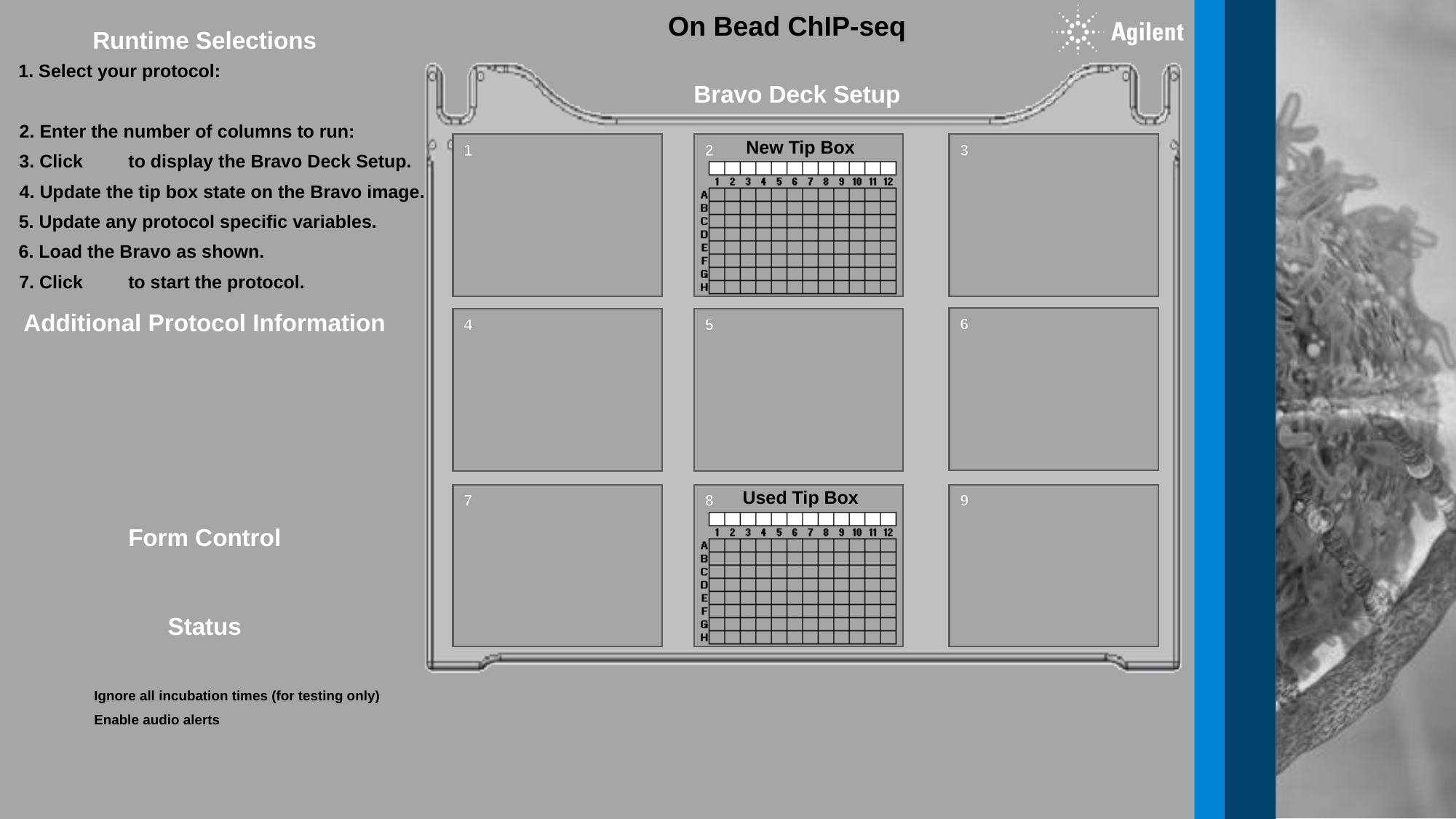

On Bead ChIP-seq
Runtime Selections
1. Select your protocol:
1
3
4
7
9
Bravo Deck Setup
2. Enter the number of columns to run:
New Tip Box
2
3. Click 	to display the Bravo Deck Setup.
4. Update the tip box state on the Bravo image.
5. Update any protocol specific variables.
6. Load the Bravo as shown.
7. Click 	to start the protocol.
Additional Protocol Information
6
5
Used Tip Box
8
Form Control
Status
Ignore all incubation times (for testing only)
Enable audio alerts
