## Supplementary material for "Automated chromatin profiling with spa-ChIP-seq uncovers the impacts of condition variations": Cao-etal-revised-2025-biorxiv-automation-files: Readme.docx

**spa-ChIP-seq protocol readme:**

The spa-ChIP-seq protocol was created using Agilent Bravo device, however, it should be compatible with any liquid handler that allows for the following requirements: (1) the ability to pipette a range of small volumes, from 1 µL to 180 µL; (2) allow for multiple liquid types from glycerol to water to ethanol, which requires different pipetting speeds for accuracy; (3) as some of the steps require removal of liquid without disturbing the pellets, spa-ChIP-seq also requires the ability to aspirate at slow speed; (4) lastly, while not required, the ability to perform mixing and temperature-controlled operations on deck reduces the operational time. Our contact information is: Yuwei Cao and Alon Goren.

Before initiating any protocols perform the following:

*Note: Any .reg files added will overwrite existing entries. If those files don’t exist on the system then protocols will not preform as intended.

1) Import labware
   a. Double click registry entries located here: C:\VWorks Workspace\NGS Option A\On-Bead_ChIP-seq_v.A1.0.2\SETUP\Labware
   b. Add the labware to the standard plate pad group
   c. Detailed instructions here: C:\VWorks Workspace\NGS Option A\On-Bead_ChIP-seq_v.A1.0.2\SETUP\How To\Adding a New Piece of Labware to VWorks.pdf
2) Import liquid classes
   a. Double click registry entries located here: C:\VWorks Workspace\NGS Option A\On-Bead_ChIP-seq_v.A1.0.2\SETUP\Liquid Classes
   b. Add the labware to the standard plate pad group
   c. Detailed instructions here: C:\VWorks Workspace\NGS Option A\On-Bead_ChIP-seq_v.A1.0.2\SETUP\How To\ Adding a New Liquid Class to VWorks.pdf
3) Add the “Sounds” Folder to the VWorks Workspace Folder
   a. Copy the “Sounds” Folder from here: C:\VWorks Workspace\NGS Option A\On-Bead_ChIP-seq_v.A1.0.2\SETUP\Sounds
   b. Paste the folder here: C:\VWorks Workspace\Sounds
4) Ensure the device file is linked to the proper Bravo profile and that it has the correct name.
   a. Find the device file here: C:\VWorks Workspace\NGS Option A\On-Bead_ChIP-seq_v.A1.0.2\Device Files
   b. The only device file used for these protocols is: Bravo_round_magnet_RedInsert.dev
   c. The Bravo profile associated with this device file is: Bravo-Red Insert Mag and Shaker.reg
   d. The Bravo should be named: Bravo – 1
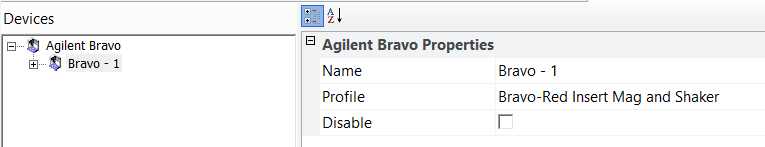

   e. Detailed instructions for creating this profile are located here: C:\VWorks Workspace\NGS Option A\On-Bead_ChIP-seq_v.A1.0.2\SETUP\How To\ NGS Bravo Setup Consolidated Profile Document.pdf
   f. An example that can be used in simulation is located here: C:\VWorks Workspace\NGS Option A\On-Bead_ChIP-seq_v.A1.0.2\SETUP\Profiles
      i. Double click on the file to add it to the registry
