## Supplementary material for "Automated chromatin profiling with spa-ChIP-seq uncovers the impacts of condition variations": Cao-etal-revised-2025-biorxiv-automation-files: Readme.pdf

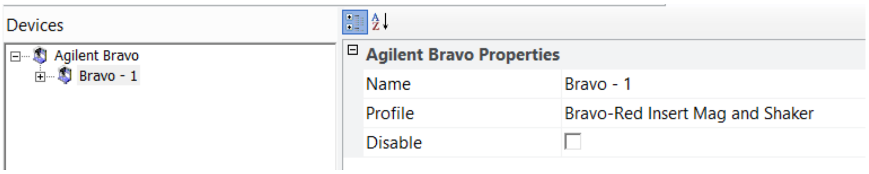


   e. Detailed instructions for creating this profile are located here: C:\VWorks Workspace\NGS Option A\On-Bead_ChIP-seq_v.A1.0.2\SETUP\How To\ NGS Bravo Setup Consolidated Profile Document.pdf

f. An example that can be used in simulation is located here: C:\VWorks Workspace\NGS Option A\On-Bead_ChIP-seq_v.A1.0.2\SETUP\Profiles
    i. Double click on the file to add it to the registry
