## Supplementary material for "Automated chromatin profiling with spa-ChIP-seq uncovers the impacts of condition variations": Cao-etal-revised-2025-biorxiv-automation-files: file_operations.js

```
// Global Debug variable
debug_file_operations = false;


// - - - - - - - - - - - - - - - - - - - - - - - - - - - -
// Functions available in this file:
// - - - - - - - - - - - - - - - - - - - - - - - - - - - -
//
//  function debug_file_ops( BOOLEAN state );
//  function file_read( STRING filepath )
//     RETURNS STRING;
//  function file_write( STRING filepath, STRING contents );
//  function file_overwrite( STRING filepath, STRING contents );
//  function file_creation( STRING folder_path, STRING file_name, STRING file_ext )
//     RETURNS STRING;
//  function get_date_time()
//     RETURNS STRING;
// - - - - - - - - - - - - - - - - - - - - - - - - - - - -


// * * * * * * * Function Definition * * * * * * * * * * *
// This function changes the debug variable to given
//  boolean value.
//
// Parameters:
//   BOOLEAN state - true or false
//
// Returns:
//   None.
// * * * * * * * * * * * * * * * * * * * * * * * * * * * *
function debug_file_ops( state )
{
    debug_file_operations = state;
    return;
} // end function debug_file_ops.


// * * * * * * * Function Definition * * * * * * * * * * *
// This function opens the given file, reads the contents,
//  and closes the file.
//
// Parameters:
//   STRING filepath - complete path to file to open
//
// Returns:
//   STRING of contents of file.
// * * * * * * * * * * * * * * * * * * * * * * * * * * * *
function file_read( filepath )
{
    // Declare file object.
    var file_obj = new File();
    var file_contents = "";

    // Opens, Reads, Closes file.
    file_obj.Open( filepath );
    file_contents = file_obj.Read();
    file_obj.Close();

    if ( debug_file_operations ) {
        print("File Read: Opened ["+filepath+"] contains \n"+
              file_contents);
    }

    return file_contents;
} // end function file_read.


// * * * * * * * Function Definition * * * * * * * * * * *
// This function opens the given file, writes the given
//  contents (appends to file), and closes the file.
//
// Parameters:
//   STRING filepath - complete path to file to open
//   STRING contents - contents to write to file
//
// Returns:
//   None.
// * * * * * * * * * * * * * * * * * * * * * * * * * * * *
function file_write( filepath, contents )
{
    // Declare file object
    var file_obj = new File();

    // Opens, Writes, and Closes file.
    file_obj.Open( filepath );
    file_obj.Write( contents+"\n" );
    file_obj.Close();

    if ( debug_file_operations ) {
        print("File Write: Opened ["+filepath+"] wrote \n"+
              contents);
    }
    return;
} // end function file_write.


// * * * * * * * Function Definition * * * * * * * * * * *
// This function opens the given file, writes the given
//  contents (overwrites file), and closes the file.
//
// Parameters:
//   STRING filepath - complete path to file to open
//   STRING contents - contents to write to file
//
// Returns:
//   None.
// * * * * * * * * * * * * * * * * * * * * * * * * * * * *
function file_overwrite( filepath, contents )
{
    // Declare file object
    var file_obj = new File();

    // Opens, Writes, and Closes file.
    file_obj.Open( filepath, 'w' );
    file_obj.Write( contents+"\n" );
    file_obj.Close();

    if ( debug_file_operations ) {
        print("File Overwrite: Opened ["+filepath+"] overwrote with \n"+
              contents);
    }
    return;
} // end function file_overwrite.


// * * * * * * * Function Definition * * * * * * * * * * *
// This function creates a file with the given parameters
// with a time-stamp appended to the filename.
//
// Parameters:
//   STRING folder_path - path to the folder
//                        (Ex: "C:/Logs/" or "C:\\Logs\\")
//   STRING file_name   - base name of file
//                        (Ex: "Replication" or "ProtocolA")
//   STRING file_ext    - the extension of the file to create
//                        (Ex: "txt" or "csv")
//
// Returns:
//   STRING of newly created file including folder path.
// * * * * * * * * * * * * * * * * * * * * * * * * * * * *
function file_creation( folder_path, file_name, file_ext )
{
    // Get the current date and time.
    var date_time_str = get_date_time();

    // Create file_path.
    var file_path = folder_path + file_name + "_"
                    + date_time_str + "." + file_ext;

    // Create the file.
    file_read( file_path );

    return file_path;
} // end function file_creation


// * * * * * * * Function Definition * * * * * * * * * * *
// This function creates a time-stamp string of the
// current date and time (with the colons replace by
// underscores in the actual time).
//
// Parameters:
//   None.
//
// Returns:
//   STRING of date and time.
// * * * * * * * * * * * * * * * * * * * * * * * * * * * *
function get_date_time()
{
    // Get Date, convert to string & remove "GMT".
    var today_str = ((new Date()).toString()).split("GMT");

    // Replace the ":" with "_" underscores.
    rExp = /:/g;
    today_str = today_str[0].replace(rExp,"_");

    // Determine length of 'today_str' to remove Weekday.
    var str_length = today_str.length;
    var date_str   = today_str.substring(4,str_length-1);

    return date_str;
} // end function get_date_time.


// end of file.
```
