## Supplementary material for "Automated chromatin profiling with spa-ChIP-seq uncovers the impacts of condition variations": Cao-etal-revised-2025-biorxiv-automation-files: protocol_set_startup.js

```
// ***********************************
// * Form javascript startup file    *
// *                                 *
// * ver 3.0 (20 April 2012)         *
// ***********************************

// declare functions

// function to display status on form

function update_status(text)
{
	form_status = text
	last_status = text
}

// function to play a wave file as a sound alert

function alert()
{
	if(form_alerts_on)
		{
		run("C:\\VWorks Workspace\\Sounds\\alert.exe")
		}

}

// function to run at end of protocol

function finish()
{
	logfile2 = new File()
	logfile2.Open(logdir + logname)
	logfile2.Write("Protocol finished successfully\n")
	logfile2.Close()

	running = 0
	alert()
}

// function to set the head mode based on the number of samples

function setHeadForRun(columns_samples)
{

	head_mode = "3,3,8," + columns_samples
  
	print("The head mode configuration is " + head_mode + "\n")

}

// function to create the incubate status text for display on form

function form_status_incubation()
{
	time = new Date()
	var hours = time.getHours().toString()
	var mins = time.getMinutes().toString()
	if(mins.length<2)
		{
		mins = "0"+mins
		}
	var secs = time.getSeconds().toString()
	if(secs.length<2)
		{
		secs="0"+secs
		}
	var incubate_text = "Incubation for "+parseInt(task.Incubationtime/60)+" mins "+parseInt(task.Incubationtime%60)+" secs started at "+hours+":"+mins+":"+secs
	return incubate_text
}

// function to set incubation time and update form

function incubate(inc_in_mins)
{
	if(form_incubations_off)
		{

		task.Incubationtime=0

		}

	else

		{

		task.Incubationtime=inc_in_mins*60
		}


	update_status(form_status_incubation())
}

// function to create the rerservation status text for display on form

function form_status_reservation()
{
	time = new Date()
	var hours = time.getHours().toString()
	var mins = time.getMinutes().toString()
	if(mins.length<2)
		{
		mins = "0"+mins
		}
	var secs = time.getSeconds().toString()
	if(secs.length<2)
		{
		secs="0"+secs
		}
	var incubate_text = "Incubation for "+parseInt(task.Reservationtime/60)+" mins "+parseInt(task.Reservationtime%60)+" secs started at "+hours+":"+mins+":"+secs
	return incubate_text
}


// function to set reserve location incubation time and update form

function reserve(res_in_mins)
{
	if(form_incubations_off)
		{

		task.Reservationtime=0

		}

	else

		{

		task.Reservationtime=res_in_mins*60
		}


	update_status(form_status_reservation())
}

// function to display user message
// will replace any <CR> with \n (newline characher)

function message(body_text)
{
		if(task.isSimulatorRunning())
			{
			task.Userdataentryintovariable = 1
			}
		else
			{
			task.Userdataentryintovariable = 0
			}
		alert()
		task.Title = "User action required"
		body_text = body_text.toString().replace(/<CR>/g,"\n")
		task.Body = body_text

}

// function to chect state of unused tip box

function check_tips()
{
	left_tip = 0
	for(i=1;i<13;i++)
		{
		if(left_tip==0)
			{
			tip = eval("box2_"+i)
			if(tip==1)
				{
				left_tip = i
				}
			}
		}
	left_available_tips = 0
	if(left_tip!=0)
		{
		ok = 1
		for(i=left_tip;i<13;i++)
			{
			if(ok==1)
				{
				tip = eval("box2_"+i)
				if(tip==1)
					{
					left_available_tips++
					}
				else
					{
					ok = 0
					}
				}
			}
		}

	right_tip = 0
	for(i=12;i>0;i--)
		{
		if(right_tip==0)
			{
			tip = eval("box2_"+i)
			if(tip==1)
				{
				right_tip = i
				}
			}
		}
	right_available_tips = 0
	if(right_tip!=0)
		{
		ok = 1
		for(i=right_tip;i>0;i--)
			{
			if(ok==1)
				{
				tip = eval("box2_"+i)
				if(tip==1)
					{
					right_available_tips++
					}
				else
					{
					ok = 0
					}
				}
			}
		}

	print("Unused tip box state")
	print("From left side "+left_available_tips+" columns starting at column "+left_tip)
	print("From right side "+right_available_tips+" columns starting at column "+right_tip)
}


// function to chect state of used tip box

function check_spaces()
{
	left_space = 0
	for(i=1;i<13;i++)
		{
		if(left_space==0)
			{
			tip = eval("box8_"+i)
			if(tip==0)
				{
				left_space = i
				}
			}
		}
	left_available_spaces = 0
	if(left_space!=0)
		{
		ok = 1
		for(i=left_space;i<13;i++)
			{
			if(ok==1)
				{
				tip = eval("box8_"+i)
				if(tip==0)
					{
					left_available_spaces++
					}
				else
					{
					ok = 0
					}
				}
			}
		}

	right_space = 0
	for(i=12;i>0;i--)
		{
		if(right_space==0)
			{
			tip = eval("box8_"+i)
			if(tip==0)
				{
				right_space = i
				}
			}
		}
	right_available_spaces = 0
	if(right_space!=0)
		{
		ok = 1
		for(i=right_space;i>0;i--)
			{
			if(ok==1)
				{
				tip = eval("box8_"+i)
				if(tip==0)
					{
					right_available_spaces++
					}
				else
					{
					ok = 0
					}
				}
			}
		}

	print("Used tip box state")
	print("From left side "+left_available_spaces+" spaces starting at column "+left_space)
	print("From right side "+right_available_spaces+" spaces starting at column "+right_space)
}


// tips on function to check if enough tips and return location in box

function tips_on_check(columns)
{
	enough = 1
	position_in_box = 0

	check_tips()

	if(columns>0)
		{
		if(right_available_tips<columns)
			{
			enough = 0
			}
		else
			{
			position_in_box = right_tip - columns + 1
			}
		}
	else
		{
		if(left_available_tips<(0-columns))
			{
			enough = 0
			}
		else
			{
			position_in_box = left_tip
			}
		}

	if(enough != 1)
		{
		form_status = "CHANGE TIP BOXES"
		task.Title = "Check Tips"
		task.Body = "CHANGE TIP BOXES\n\nPress RESET on form and CONTINUE to reset to Full boxes or press PAUSE to manually edit."
		if(task.isSimulatorRunning())
			{
			task.Userdataentryintovariable = 1
			}
		else
			{
			task.Userdataentryintovariable = 0
			}
		alert()
		running = 0
		task.repeat()
		}
	else
		{
		form_status = last_status
		running = 1
		task.skip()
		}

	
}


// tips on function to check if enough empty spaces and return location in box

function tips_off_check(columns)
{
	enough = 1
	position_in_box = 0

	check_spaces()

	if(columns>0)
		{
		if(left_available_spaces<columns)
			{
			enough = 0
			}
		else
			{
			position_in_box = left_space
			}
		}
	else
		{
		if(right_available_spaces<(0-columns))
			{
			enough = 0
			}
		else
			{
			position_in_box = right_space + columns + 1
			}
		}

	if(enough != 1)
		{
		form_status = "CHANGE TIP BOXES"
		task.Title = "Check Tips"
		task.Body = "CHANGE TIP BOXES\n\nPress RESET on form and CONTINUE to reset to Full boxes or press PAUSE to manually edit."
		if(task.isSimulatorRunning())
			{
			task.Userdataentryintovariable = 1
			}
		else
			{
			task.Userdataentryintovariable = 0
			}
		running = 0
		alert()
		task.repeat()
		}
	else
		{
		form_status = last_status
		running = 1
		task.skip()
		}	
}

// function for tips on and to track tips used

function tips_on(column)
{
    task.Wellselection = [[1,position_in_box]]
                
    print("Tips on at column "+position_in_box)

    for(i=position_in_box;i<(position_in_box+Math.abs(column));i++)
        {
        eval("box2_"+i+" = 0")
        }
}
 
// function for tips off and to track spaces available

function tips_off(column)
{
    task.Wellselection = [[1,position_in_box]]
                
    print("Tips off at column "+position_in_box)

    for(i=position_in_box;i<(position_in_box+Math.abs(column));i++)
        {
        eval("box8_"+i+" = 1")
        }
}


// function to track tips on for cherry picking

function tips_on_cherry()
{
	task.Wellselection = [[tip_row,position_in_box]]
	
	print("Tips on at column "+position_in_box+" and row "+tip_row)
	
	if(tip_row == 8)
	{	
		for(i=position_in_box;i>(position_in_box-1);i--)
			{
			eval("box2_"+i+" = 0")
			}
	}
}

// function to tips off for cherry pick

function tips_off_cherry()
{
	task.Wellselection = [[tip_row,position_in_box]]
	
	print("Tips off at column "+position_in_box+ " and row "+tip_row)

	if(tip_row == 8)
		
		{
		for(i=position_in_box;i>(position_in_box-1);i--)
			{
			eval("box8_"+i+" = 1")
			}
		tip_row = 1
		}
	else
		{
		tip_row++
		}
}


print("Javascript functions declared")


// define variables

var headmode
var form_status
var columns_samples = parseInt(form_columns_samples)
var left_tip
var left_available_tips
var right_tip
var right_available_tips
var left_space
var left_available_spaces
var right_space
var right_available_spaces
var tip_row = 1
var hit = 1
var last_status
var position_in_box
var running = 1
var logdir = "c:\\VWorks Workspace\\VWorks\\Logs\\"
var logname = ""


// clear form status

update_status("")


// get headmode based on number of columns

print("The number of columns that will be processed is " + columns_samples + ".")

setHeadForRun(columns_samples)


// get protocol variables from CSV file
/*
print("Getting data from csv file....")

protocol_variable = new Array()
protocol_data = new Array()

protocol = new File()
protocol.Open(protocol_settings_filename)
protocol_contents = protocol.Read()
protocol.Close()

print(protocol_contents)

protocol_line = protocol_contents.split("\n")

num_var = 0
for(i=1;i<protocol_line.length;i++)
	{
	if(protocol_line[i]!="")
		{
		num_var++
		protocol_data[i] = protocol_line[i].split(",")
		}
	}

for(i=1;i<num_var+1;i++)
	{
	protocol_variable[i]=protocol_data[i][2]
	print("protocol_variable["+i+"] = "+protocol_variable[i]+" ("+protocol_data[i][0]+")")
	}
*/
// Log the run was started

// create log name from the time

	ltime = new Date()
	var lyear = ltime.getFullYear().toString()
	var lmonth = (ltime.getMonth()+1).toString()
	if(lmonth.length<2)
		{
		lmonth = "0"+lmonth
		}
	var ldate = ltime.getDate().toString()
	if(ldate.length<2)
		{
		ldate = "0"+ldate
		}
	var lhours = ltime.getHours().toString()
	if(lhours.length<2)
		{
		lhours = "0"+lhours
		}
	var lmins = ltime.getMinutes().toString()
	if(lmins.length<2)
		{
		lmins = "0"+lmins
		}
	var lsecs = ltime.getSeconds().toString()
	if(lsecs.length<2)
		{
		lsecs="0"+lsecs
		}

	logname = "XTLI_ILM_optionA("+lyear+"_"+lmonth+"_"+ldate+" "+lhours+"-"+lmins+"-"+lsecs+").log"

// Open log file

	logfile = new File()
	logfile.Open(logdir +logname,"w")

// Write header

	logfile.Write("On-Bead_ChIP-seq Option A Log file\n----------------------------\n\n")

// Write data

	logfile.Write("protocol:        "+task.getProtocolName()+" \n\n")

	var sim_on = "OFF"
	if(task.isSimulatorRunning()==1)
		{
		sim_on = "ON"
		}
	logfile.Write("simulation mode: "+sim_on+"\n\n")


	logfile.Write("started:         "+lyear+"\\"+lmonth+"\\"+ldate+" "+lhours+":"+lmins+":"+lsecs+"\n\n")
	logfile.Write("columns:         "+columns_samples+"\n\n")

	var alerts_on = "OFF"
	if(form_alerts_on==1)
		{
		alerts_on = "ON"
		}
	logfile.Write("audio alerts:    "+alerts_on+"\n\n")

	var incs_on = "ON"
	if(form_incubations_off==1)
		{
		incs_on = "OFF"
		}
	logfile.Write("incubations:     "+incs_on+"\n\n\n")

// write protocol data
/*	
	logfile.Write("Protocol Data:\n\n")
	for(i=1;i<num_var+1;i++)
		{
		logfile.Write("protocol_variable["+i+"] = "+protocol_data[i][2]+" ("+protocol_data[i][0]+")\n")
		}
	logfile.Write("\n\n")
*/
// Close log file

	logfile.Close()

// stratup script finished

print("Startup script complete")
```
