## Supplementary material for "Automated chromatin profiling with spa-ChIP-seq uncovers the impacts of condition variations": Cao-etal-revised-2025-biorxiv-automation-files: Adding a New Liquid Class to VWorks.pdf

1. Verify the system type, 32 or 64 bit, by going to Control Panel > System and Security > System

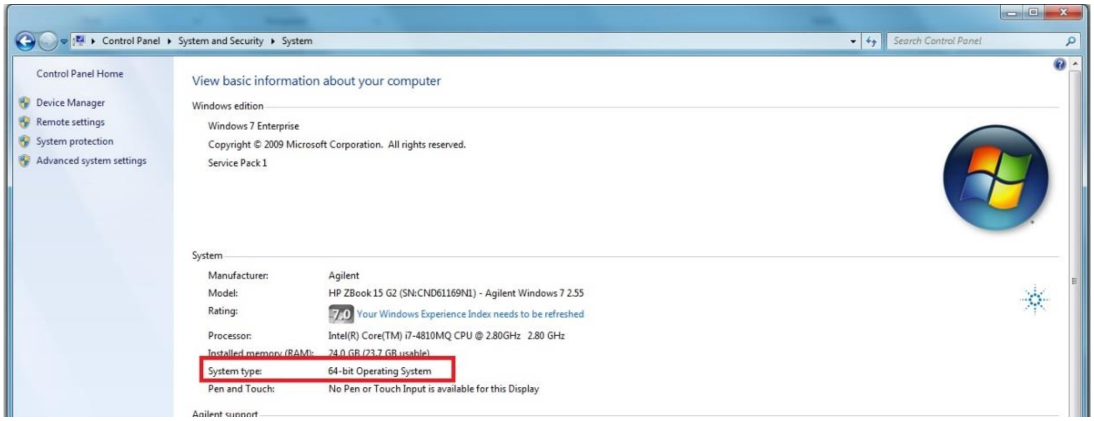


2. Find the appropriate file, 32 or 64 bit. If file is send as a .txt file, rename changing the extension to .reg
3. Double-click on the .reg file. You will get the following error

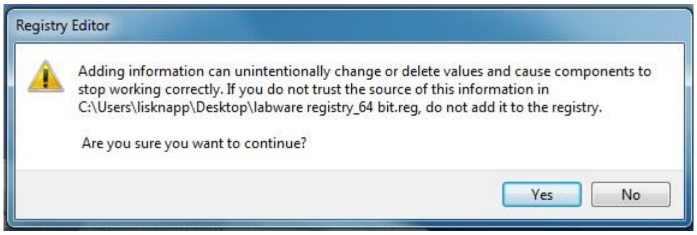


Select "Yes"

4. Once you have done this, you will get the following message

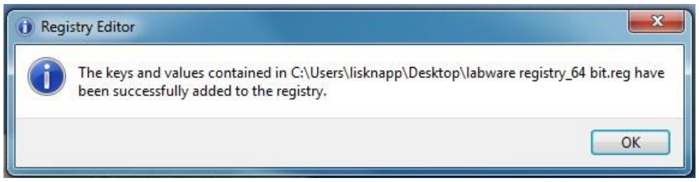
