## Supplementary material for "Automated chromatin profiling with spa-ChIP-seq uncovers the impacts of condition variations": Cao-etal-revised-2025-biorxiv-automation-files: Adding a New Piece of Labware to VWorks.pdf

1. Verify the system type, 32 or 64 bit, by going to Control Panel > System and Security > System

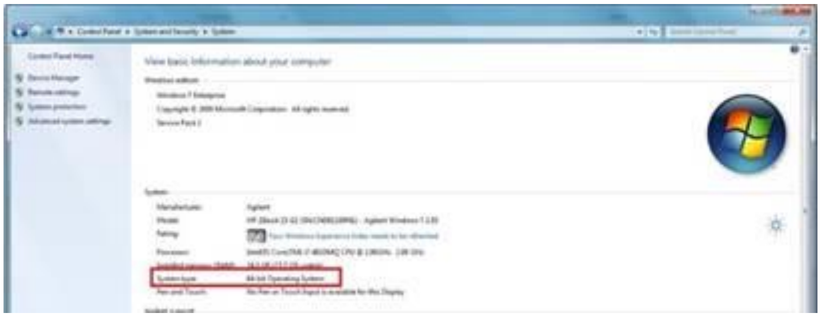

2. Find the appropriate file, 32 or 64 bit. If file is send as a .txt file, rename changing the extension to .reg

3. Double-click on the .reg file. You will get the following error

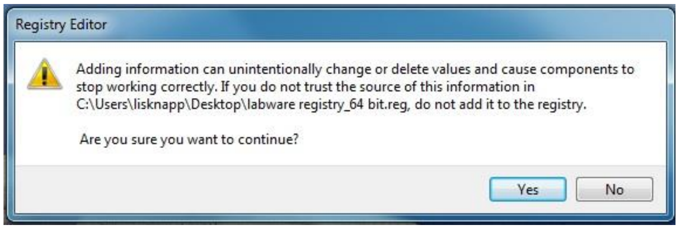


   Select “Yes”

4. Once you have done this, you will get the following message

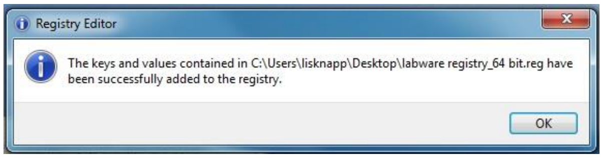


5. Once you have done this, go into Tools > Labware Editor. Follow the steps in the following image to make sure the labware is in the “Uses Standard Platepad” Labware Class.

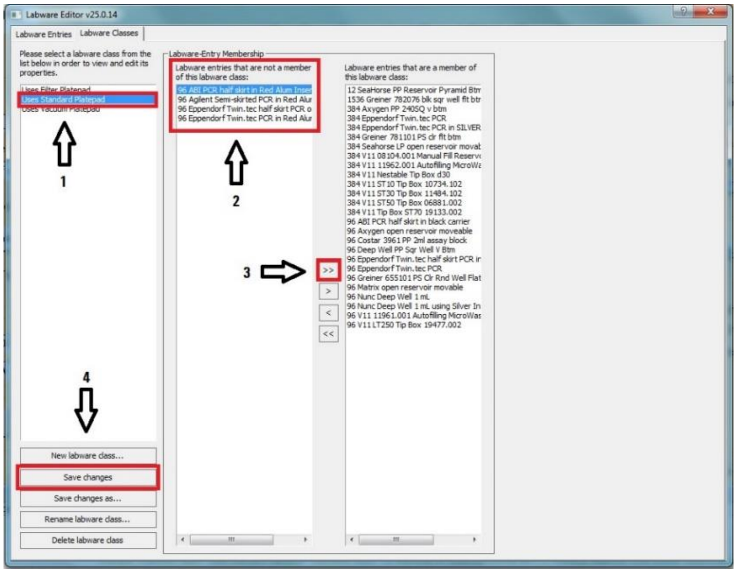
