## Supplementary material for "Automated chromatin profiling with spa-ChIP-seq uncovers the impacts of condition variations": Cao-etal-revised-2025-biorxiv-automation-files: Installing VWorks Software to Mimic NGS Option A Workstation.pdf

1. Install VSCodeUserSetup-x64-1.39.2.exe, Microsoft Visual C++ Redistributable for Visual Studio, x86 if not already installed on your computer. If it is not installed, you will receive an error when installing VWorks 13.1.0.1366

2. Verify Microsoft .NET Framework v2 is installed. It looks like this omission was due to a change in the latest release of Windows 10. In their infinite wisdom, Microsoft decided *not* to include version 2 of the .NET framework by default, probably figuring that all the cool kids were using .NET 4 these days. Previous builds of Windows 10 had .NET 2 and 4 both installed by default. If not installed, here is a link to download and install.
    a. Download ASP.NET MVC 2 RTM from Official Microsoft Download Center

3. Install VWorks 13.1.0.1366

4. Take the zip files named Sounds.zip and Inventory Backup.zip, unzip and place the folders in C:\VWorks Workspace

5. Install labware_64 bit.reg, liquid library_64 bit.reg, and the 3 profiles Bravo-Mag and Shaker.reg, Bravo-Red Insert Mag and Shaker, and Bravo-2 Inserts Mag and Shaker.reg following the instructions to install the liquid library.

6. Take the files named 500ng transfer.xml and AliquotLibraries.xml and place them in C:\VWorks Workspace\VWorks\Hit Picking\Format Files

7. In C:\VWorks Workspace\VWorks\Hit Picking\ create a folder named Pooling Files

8. Take the file named a.xml and save to C:\Program Files (x86)\Agilent Technologies\VWorks\Users. This creates the username “a” with no password.
