## Supplementary material for "Automated chromatin profiling with spa-ChIP-seq uncovers the impacts of condition variations": Cao-etal-revised-2025-biorxiv-automation-files: NGS Bravo Setup Consolidated Profile Document.pdf

**NGS Workstation Setup:**

Bravo Profiles: You will need two Bravo profiles for the NGS workstation. The first is: "Bravo-Mag and Shaker" and is taught to have V11 Inheco pads at positions 4 and 6. The other is "Bravo-2 Inserts Mag and Shaker" and is taught to have the Red PCR plate insert at position 4 and the Aluminum Nunc insert at Position 6.

***Bravo Teaching: Profile "*****Bravo-Mag and Shaker*****"***

All Z-axis teaching will be done using a 0.010 inch (0.254 mm) feeler gauge to set the height of the tips from the pad. To teach the height, load a column of tips and lower the head in 0.2 mm increments until the feeler gauge cannot fit under the tips then stop. Next raise the head by 0.05mm increments until you reach the lowest possible height that allows the 0.010" feeler gauge to slide under all of the tips without deflecting them.

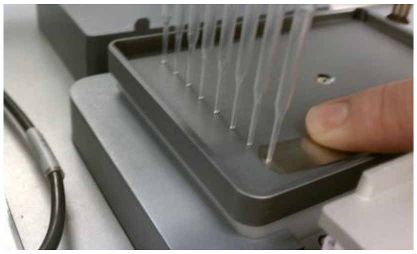

Position 1 – 3 (Standard Bravo Pads):
Teach position as normal and set height to a 0.010" gap between the tip and the pad.

Position 4 (Inheco Heating Pad):
1) Mount V11 pad onto Inheco heating Pad
2) Set Teaching insert on pad (10mm tall block with + at A1 position)
3) Teach position on teaching insert and set height to a 0.010" gap between the tip and the insert.
4) Raise Bravo head and Remove Teaching insert
5) Move to teachpoint
6) Lower Bravo by 10mm
7) Recheck height to ensure that tip is spaced 0.010" above the pad

Position 5 (VarioMag TeleShake):
Teach position as normal and set height to a 0.010" gap between the tip and the pad.
1) Set Teaching insert on pad (10mm tall block with + at A1 position)
2) Teach position on teaching insert and set height to a 0.010" gap between the tip and the insert.
3) Raise Bravo head and Remove Teaching insert
4) Move to teachpoint
5) Lower Bravo by 10mm
6) Set teachpoint here

Position 6 (Inheco Heating Pad):

1) Mount V11 pad onto Inheco heating Pad
2) Set Teaching insert (10mm tall block with + at A1 position)
3) Teach position on teaching insert and set height to a 0.010” gap between the tip and the insert.
4) Raise Bravo head and Remove Teaching insert
5) Move to teachpoint
6) Lower Bravo head by 10mm
7) Recheck height to ensure that tip is spaced 0.010” above the pad

Position 7 (Agencourt Magnet plate):
First securely mount Agencourt magnet to backing plate without included springs to eliminate any warping of Agencourt Magnet

1) Remove 4 corner screws from Agencourt magnet
2) Remove springs from between black plastic magnet mounting and metal backing plate.
3) Replace screws with shorter ones, or use small washers(size M3 works well) to fill the gap to allow screws to be tightened to press plastic magnet mounting firmly onto backing with stickerless side facing down.

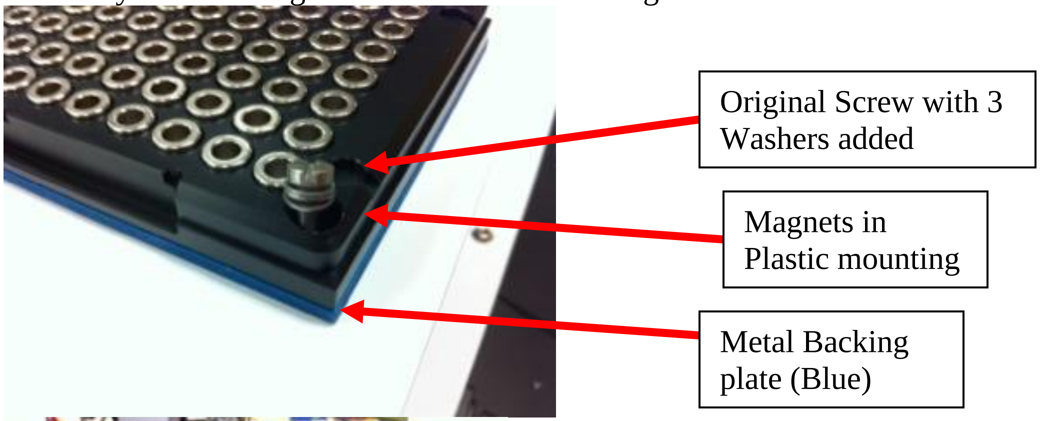


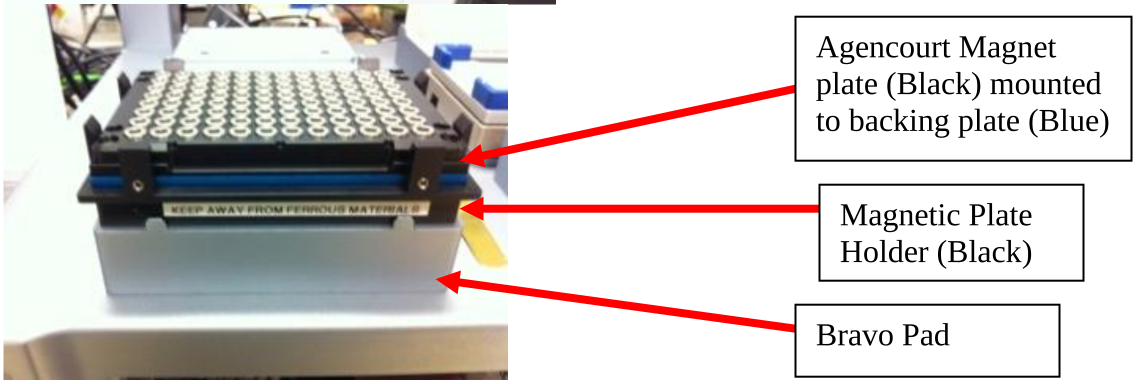


Teaching Position 7:

1) Teach X/Y position to the + on gray Bravo pad
2) Teach Z axis to 0.010” tip spacing above the gray Bravo pad
3) Raise head by 29.6 mm
4) Teach Position 7 Here
5) **Move to safe height at current X/Y position**

6) Mount Magnet to Plate holder
7) **Move** to Position 7 teachpoint
8) Adjust X and Y axes to center tips in Magnet rings.
    a. Recommended: Verify X/Y with a full head of tips.

Position 8 (Standard Bravo Pads):
Teach position as normal and set height to a 0.010” gap between the tip and the pad.

Position 9 (Mecour Thermal Pad):
Teach position as normal and set height to a 0.010” gap between the tip and the pad.
7) Set Teaching insert on pad (10mm tall block with + at A1 position)
8) Teach position on teaching insert and set height to a 0.010” gap between the tip and the insert.
9) Raise Bravo head and Remove Teaching insert
10) Move to teachpoint
11) Lower Bravo by 10mm
12) Set teachpoint here

***Teaching Bravo Profile: “*Bravo-Red Insert Mag and Shaker*”***

1) Close “Bravo-2 Inserts Mag and Shaker” profile
2) Remove tips from Bravo Head.
3) Delete old “Bravo-Red Insert Mag and Shaker” profile
4) Create a copy of “Bravo-Mag and Shaker” profile
5) Rename copy to: “Bravo-Red Insert Mag and Shaker”
6) Initialize “Bravo-Red Insert Mag and Shaker”
7) In the Jog/Teach tab:
    a. For Red PCR plate insert at Position 4:
        i. **Move** to Position 4
        ii. Raise the Bravo head (-z) 4.4 mm and set teachpoint.
8) In Profile tab Update this profile.

***Teaching Bravo Profile: “*Bravo-2 Inserts Mag and Shaker*”***

1) Close “Bravo-Mag and Shaker” profile
2) Remove tips from Bravo Head.
3) Delete old “Bravo-2 Inserts Mag and Shaker” profile
4) Create a copy of “Bravo-Mag and Shaker” profile
5) Rename copy to: “Bravo-2 Inserts Mag and Shaker”
6) Initialize “Bravo-2 Inserts Mag and Shaker”
7) In the Jog/Teach tab:
    a. For Red PCR plate insert:,
        i. **Move** to Position 4

        ii. Raise the Bravo head (-z) 4.4 mm and set teachpoint.
    b. For Nunc Plate SBS format Aluminum Insert
        i. **Move** to Position 6
        ii. Raise the Bravo head (-z) 3.3 mm and set teachpoint
            - For older style, non-SBS footprint aluminum insert that screws directly to CPAC heated position: Lower the Bravo head (+z) 4.45 mm and set teachpoint

8) In Profile tab Update this profile.

Double checking heights:
To Check that the Z-Height is properly set on these positions (especially Positions 4 and 6 Inheco pads, and Position 7 Magnet pad due to possible variations in inserts and magnets), take an Eppendorf PCR plate (for Position 4 or 7) or a Nunc Deep Well plate on the Inheco insert (for Position 6), and perform a Mix at 0.1 mm (Eppendorf) or 0.85 mm (Nunc) above the bottom of the plate. When the tips are all the way down, the plate should be able to move slightly from side to side, but there should not be any up and down movement. They plate should NOT be pinned to the position by the tips so that it cannot move at all. If the Plate is being pinned to the bottom of the position, adjust the teachpoint up slightly and test again. If the plate can be moved more than just side to side, and can be lifted slightly off of the position, adjust the teachpoint down slightly and test again.

***NOTES***
384 silver insert is 6.4 mm tall
