## Supplementary material for "Automated chromatin profiling with spa-ChIP-seq uncovers the impacts of condition variations": Cao-etal-revised-2025-biorxiv-automation-files: VWorks software computer requirements.pdf

If your organization uses a computer other than one configured by Agilent Technologies, make sure the computer meets the following minimum requirements.

### Computer

- Microsoft Windows 7 64-bit or Windows 10 64-bit operating system

  *Note:* The G5562A, G5563A Bravo Platform is verified only on the Windows 10 64-bit operation system.
- 2 GHz or faster 32-bit (x86) processor, multicore preferred
- 4 GB RAM
- 40 GB hard drive capacity with 10 GB free space
- 1280 x 1024 pixel screen resolution
- MySQL 5.1 (Windows 7) or MySQL 5.7 (Windows 10) for systems with labware storage devices, such as the Labware MiniHub
- Microsoft Excel (required for running AssayMAP calculators)
- One of the following browsers with JavaScript enabled (required for using the context-sensitive help and knowledge base):
  - Microsoft Edge on Windows 10
  - Microsoft Internet Explorer 6.x, 7.x, 8.x, 9.x, 10.x, 11.x
  - Mozilla Firefox 1.x,- 39.x

  *Note:* When using Internet Explorer to display the help topics, you might have to allow local files to run active content (scripts and ActiveX controls). To do this, open the Internet Options dialog box in Internet Explorer. Click the Advanced tab, locate the Security section, and select Allow active content to run in files on my computer.

  *Note:* You may use Google Chrome to view the help topics in the online web version of the knowledge base, but not to display the local help within the VWorks software.
- A PDF viewer, such as Adobe Reader (required for opening the user guide PDF files)

### Communications interface

Depending on the device requirements:

- Dedicated 10BaseT or faster Ethernet card (two network cards if connecting to your local area network)
- RS-232 DB9 serial port, if the device requires a serial connection

  Alternatively, you may connect using a USB port with USB-to-serial adapter.

To facilitate the setup process, a software installation CD is supplied. You can use the CD to install the necessary software and setup configurations.
