## Supplementary material for "Automated chromatin profiling with spa-ChIP-seq uncovers the impacts of condition variations": Cao-etal-revised-2025-biorxiv-automation-files: readme.txt

Readme for alert.exe
____________________

alert.exe is a small wav file player that does not have a GUI. It plays the file

alert.wav

located in the directory

C:VWorks Workspace\Sounds

The sound played can be altered by changing the alert.wav.

To run from VWorks use the run() function.

Note: There are two versions supplied one for .NET 3.5 and one for .NET 4 depending on the
customers installation.
 
_____________________
30/7/2011
M Bimson
Agilent Technologies
