## Supplementary figures and images for "Automated chromatin profiling with spa-ChIP-seq uncovers the impacts of condition variations"

### Img_Key_01a_8.png

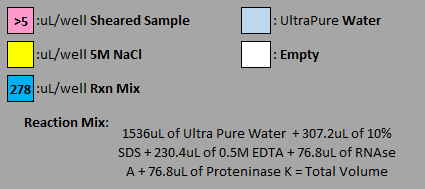

### Img_Key_01a_9.png

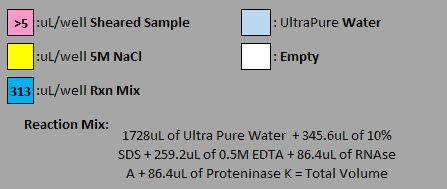

### Img_Key_01a_10.png

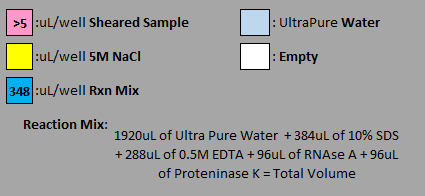

### Img_Key_01a_11.png

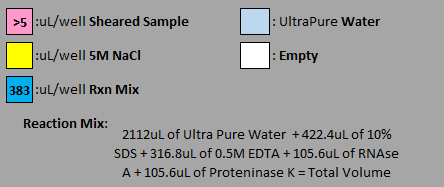

### Img_Key_01a_12.png

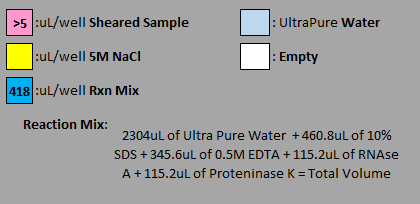

### Img_Key_01b_9.png

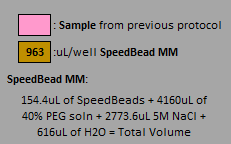

### Img_Key_05_8.png

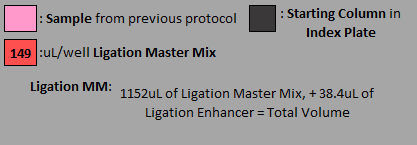

### Img_Key_05_9.png

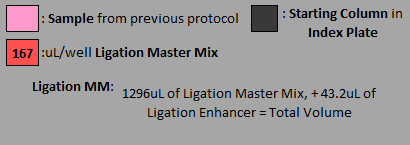

### Img_Key_05_10.png

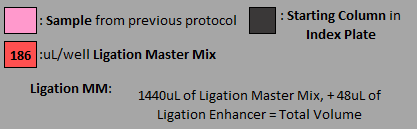

### Img_Key_05_11.png

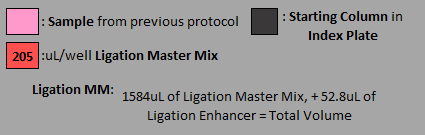

### Img_Key_05_12.png

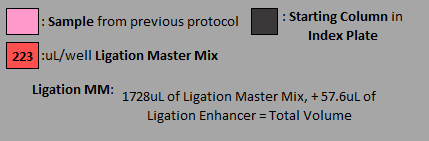

### Img_Key_06_9.png

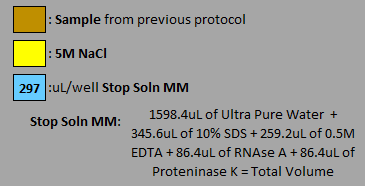

### Img_Key_08_1.png

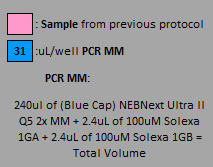

### Img_Key_08_2.png

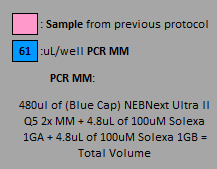

### Img_Key_08_3.png

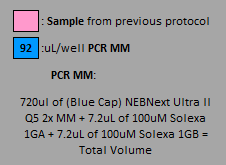

### Img_Key_08_6.png

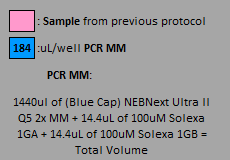

### Img_Key_08_7.png

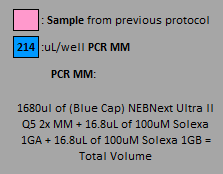

### Img_Key_09a_8.png

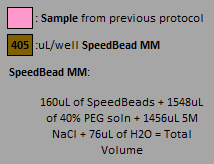

### Img_Key_09a_9.png

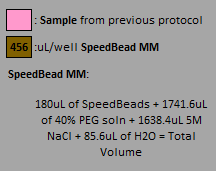

### Img_Key_09b.png

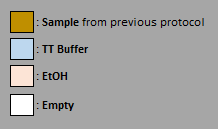

### Img_MM_01.png

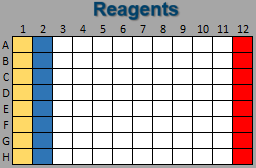

### Img_MM_02.png

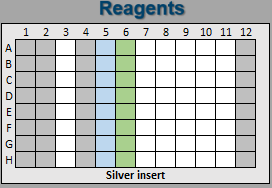

### Img_MM_03.png

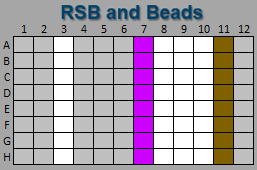

### Img_MM_05_10.png

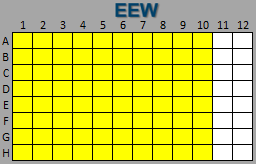

### Img_MM_05_11.png

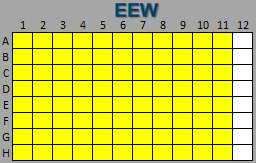

### Img_MM_05_12.png

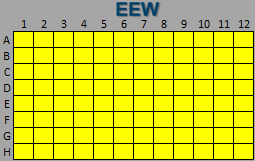

### Img_MM_05_A1.png

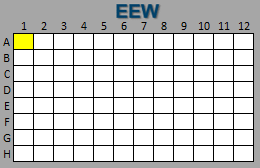

### Img_MM_06_red.png

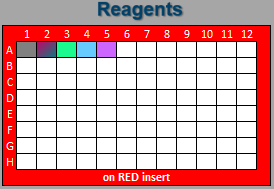

### Img_MM_07_red.png

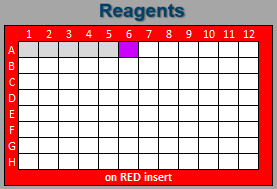
