## Supplementary figures and images for "Automated chromatin profiling with spa-ChIP-seq uncovers the impacts of condition variations"

### agilentlogo-home.bmp

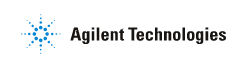

### alert_arrow.png

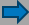

### alert_blank.png

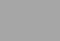

### Background Img Btm.png

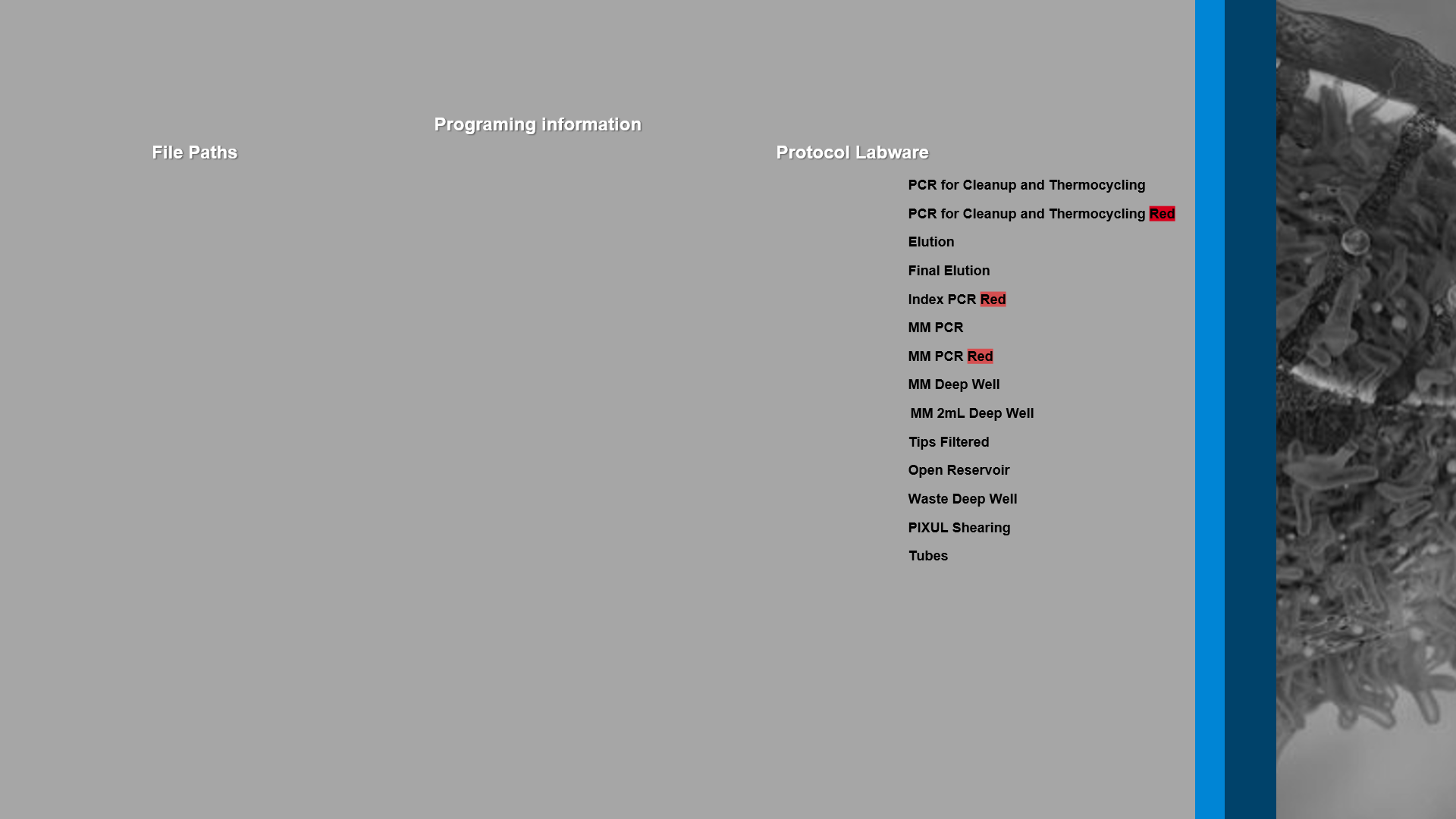

### Background Img Btm_PlateMap.png

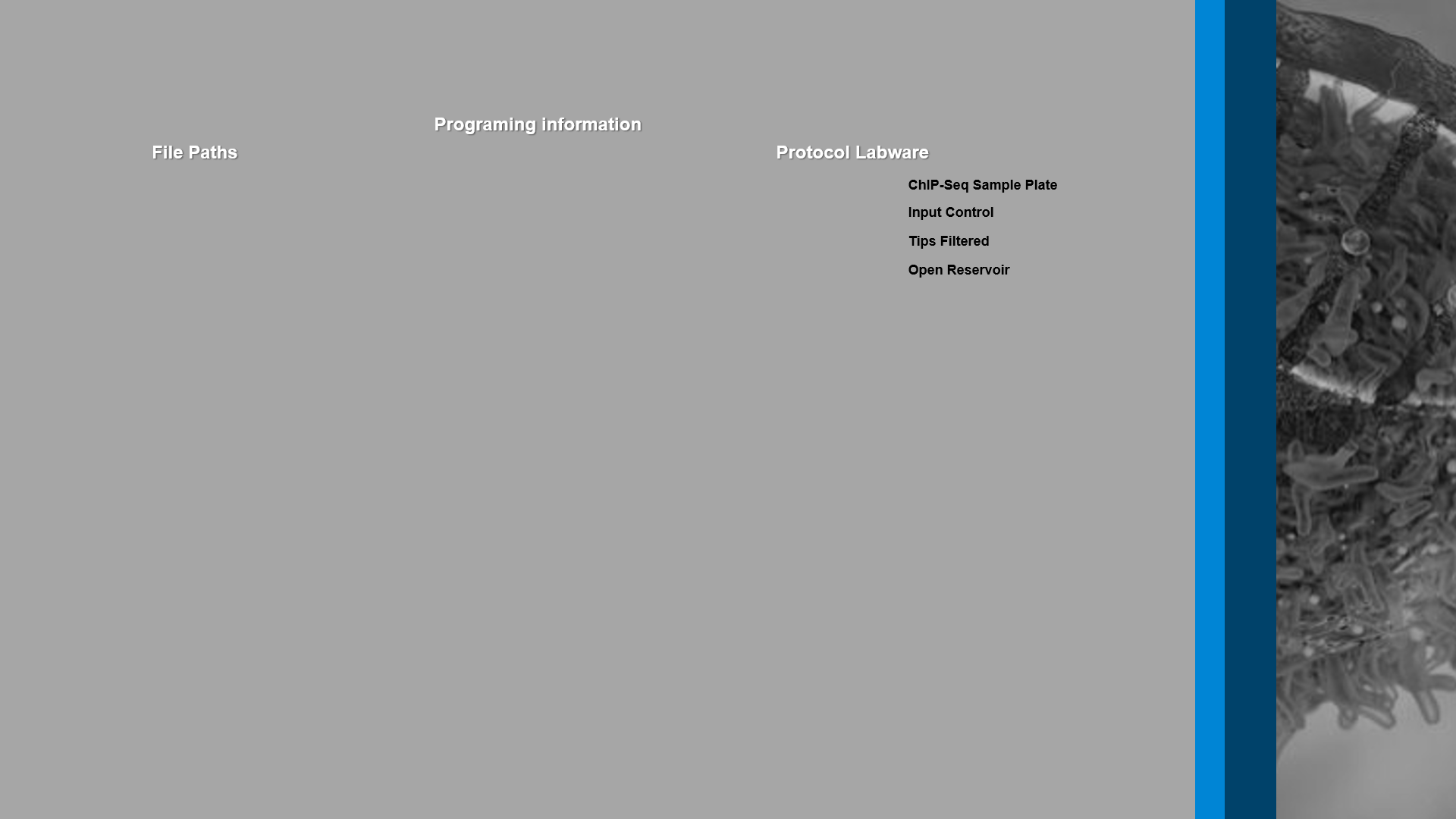

### Background Img Top.png

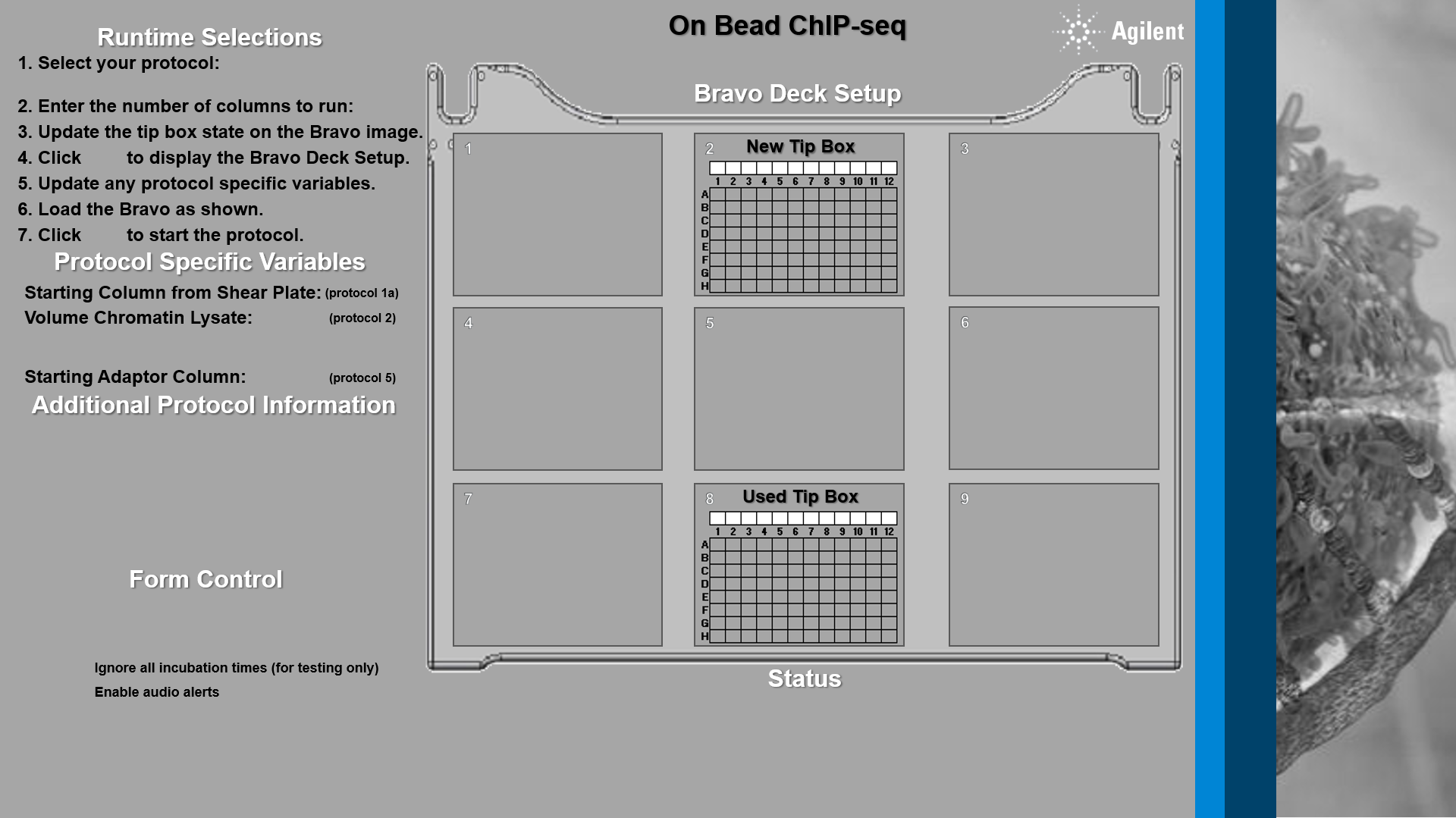

### Background Img Top_PlateMap.png

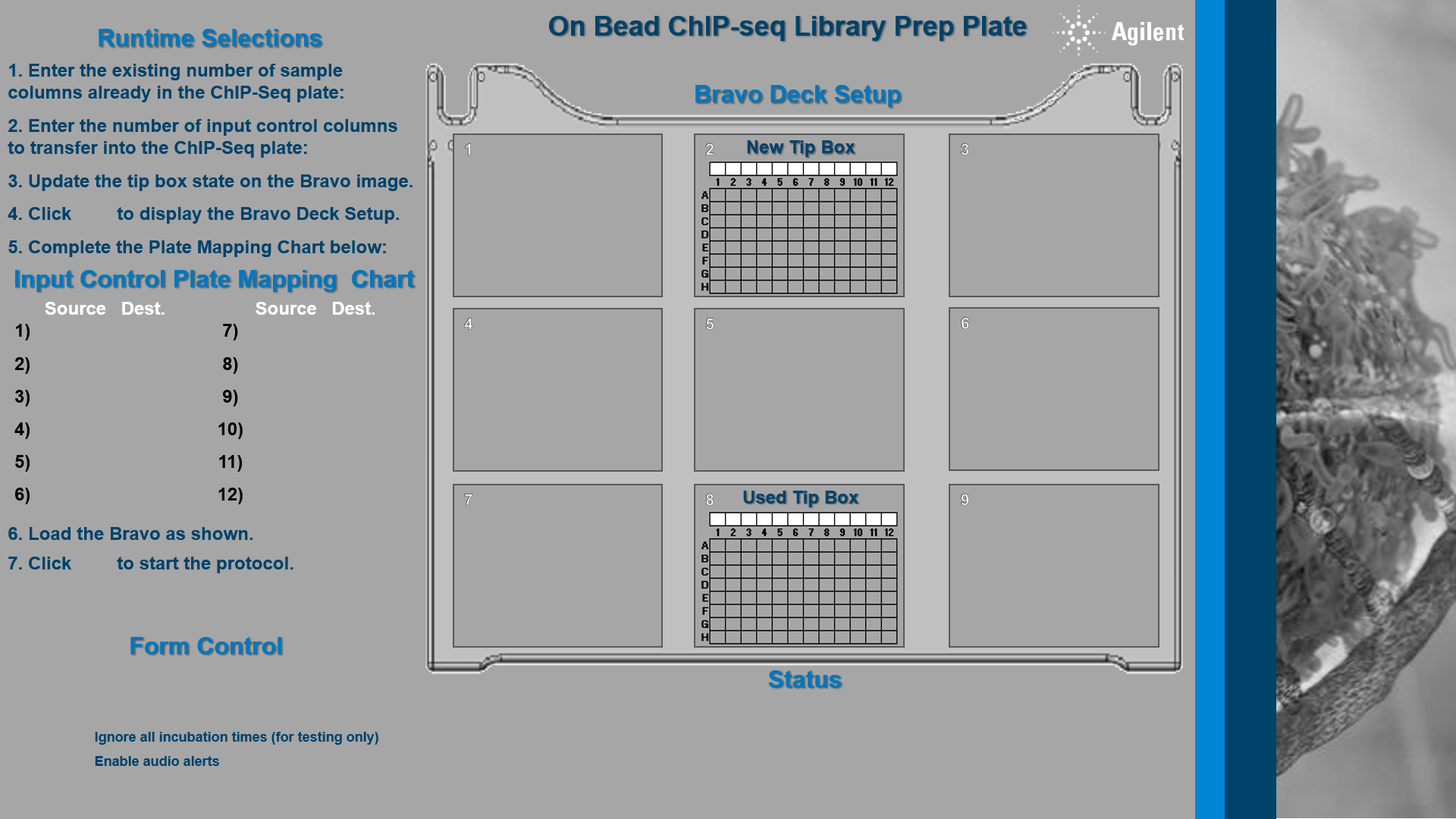

### Bravo Image-small Clear_Option A.png

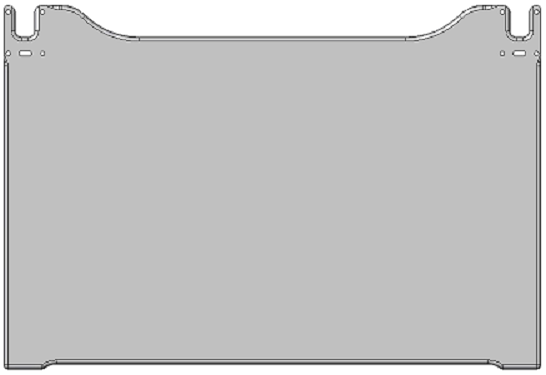

### icon_check_small.png

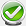

### img_Blank.png

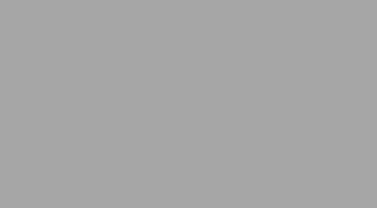

### img_Blue.png

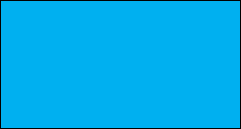

### img_Empty.png

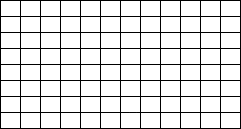

### img_Green.png

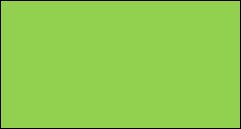

### Img_Key_01a_1.png

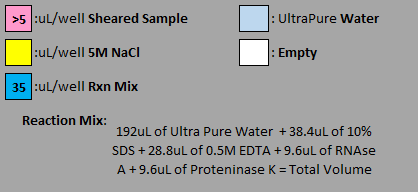

### Img_Key_01a_2.png

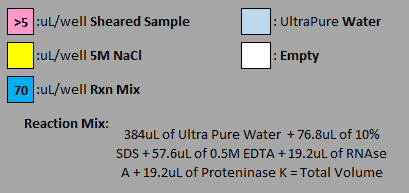

### Img_Key_01a_3.png

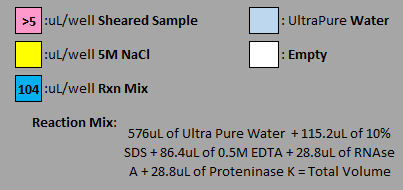

### Img_Key_01a_4.png

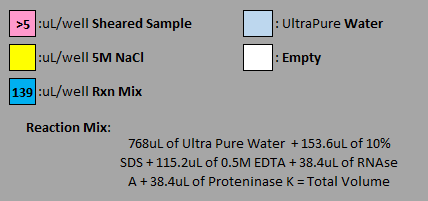

### Img_Key_01a_5.png

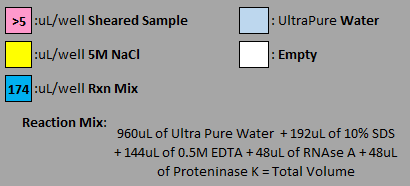

### Img_Key_01a_6.png

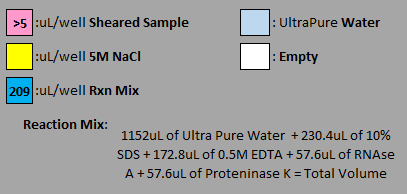

### Img_Key_01a_7.png

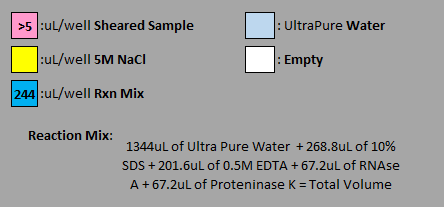

### Img_Key_01b_1.png

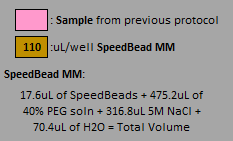

### Img_Key_01b_2.png

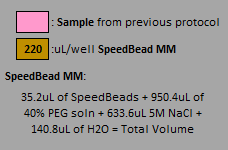

### Img_Key_01b_3.png

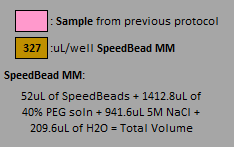

### Img_Key_01b_4.png

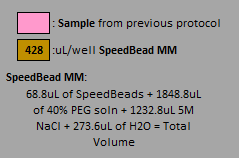

### Img_Key_01b_5.png

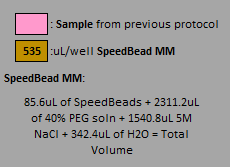

### Img_Key_01b_6.png

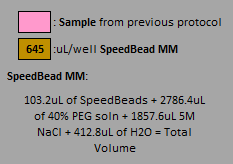

### Img_Key_01b_7.png

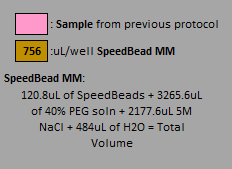

### Img_Key_01b_8.png

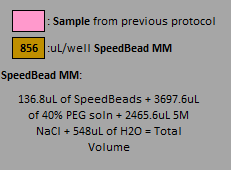

### Img_Key_01b_10.png

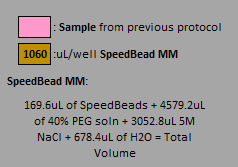

### Img_Key_01b_11.png

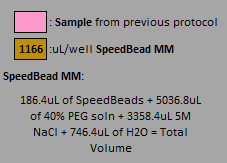

### Img_Key_01b_12.png

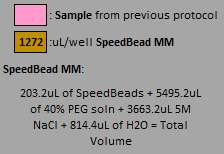

### Img_Key_01c.png

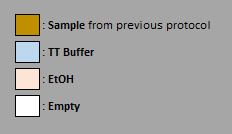

### Img_Key_02.png

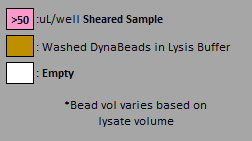

### Img_Key_03.png

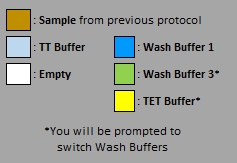

### Img_Key_04_1.png

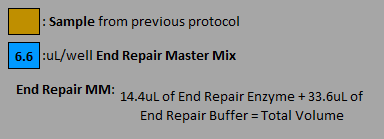

### Img_Key_04_2.png

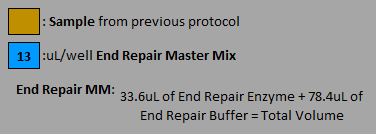

### Img_Key_04_3.png

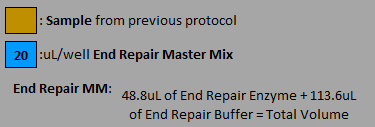

### Img_Key_04_4.png

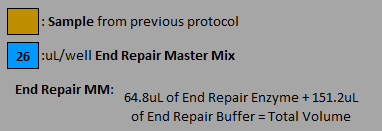

### Img_Key_04_5.png

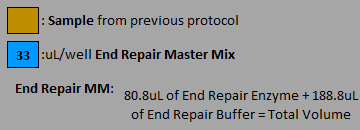

### Img_Key_04_6.png

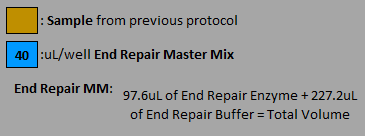

### Img_Key_04_7.png

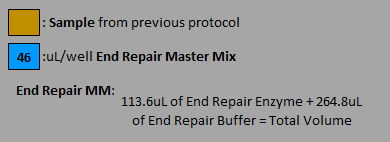

### Img_Key_04_8.png

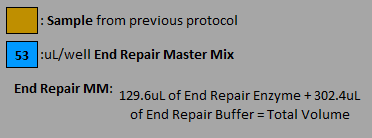

### Img_Key_04_9.png

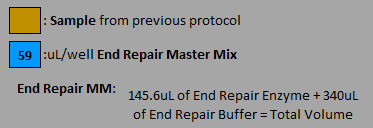

### Img_Key_04_10.png

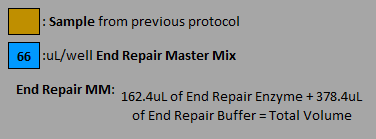

### Img_Key_04_11.png

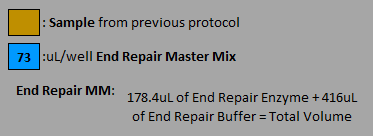

### Img_Key_04_12.png

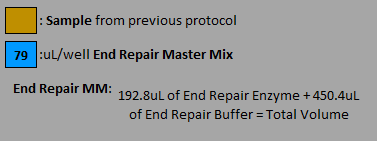

### Img_Key_05_1.png

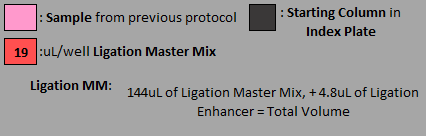

### Img_Key_05_2.png

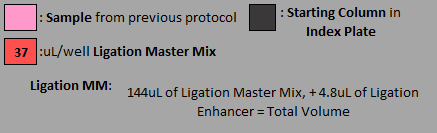

### Img_Key_05_3.png

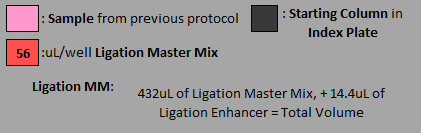

### Img_Key_05_4.png

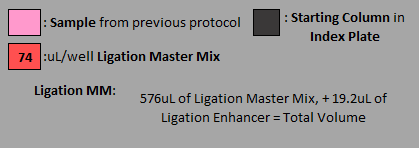

### Img_Key_05_5.png

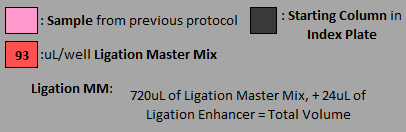

### Img_Key_05_6.png

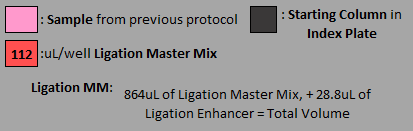

### Img_Key_05_7.png

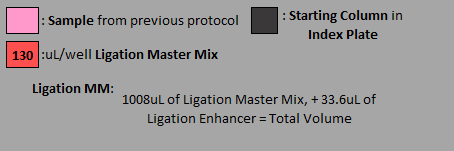

### Img_Key_06_1.png

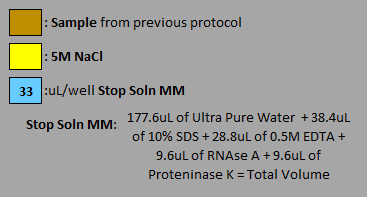

### Img_Key_06_2.png

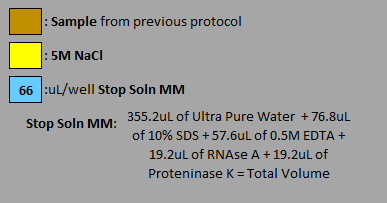

### Img_Key_06_3.png

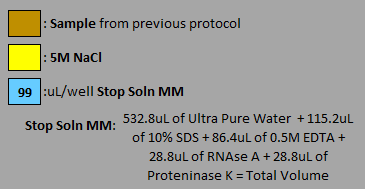

### Img_Key_06_4.png

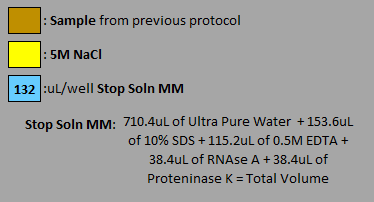

### Img_Key_06_5.png

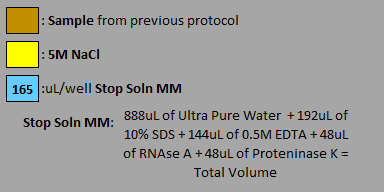

### Img_Key_06_6.png

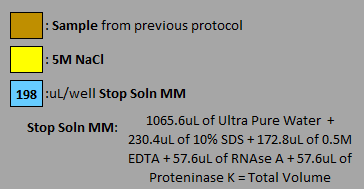

### Img_Key_06_7.png

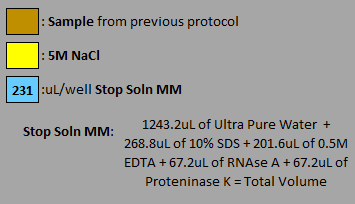

### Img_Key_06_8.png

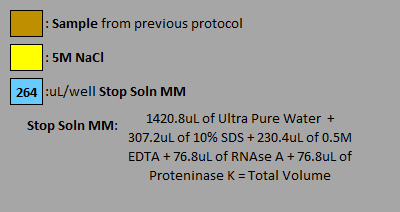

### Img_Key_06_10.png

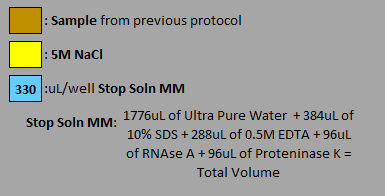

### Img_Key_06_11.png

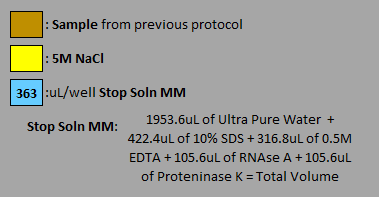

### Img_Key_06_12.png

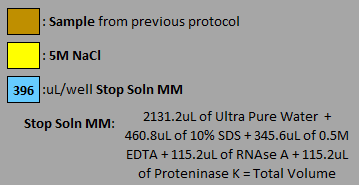

### Img_Key_07a_1.png

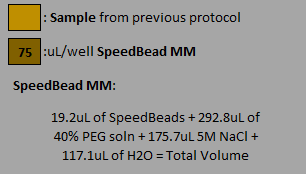

### Img_Key_07a_2.png

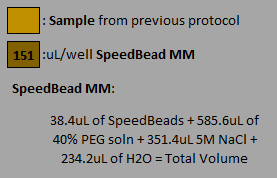

### Img_Key_07a_3.png

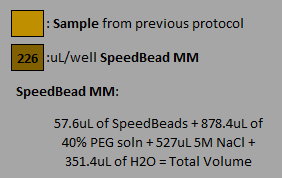

### Img_Key_07a_4.png

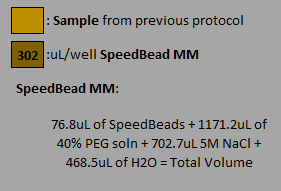

### Img_Key_07a_5.png

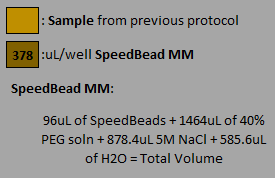

### Img_Key_07a_6.png

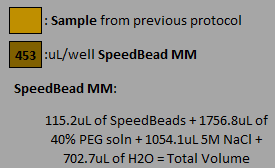

### Img_Key_07a_7.png

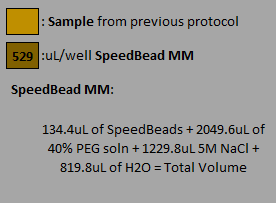

### Img_Key_07a_8.png

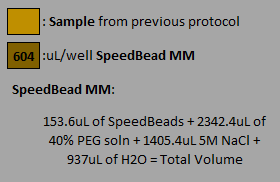

### Img_Key_07a_9.png

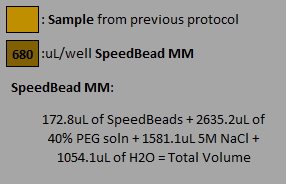

### Img_Key_07a_10.png

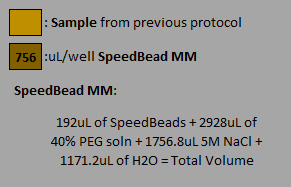

### Img_Key_07a_11.png

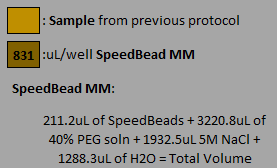

### Img_Key_07a_12.png

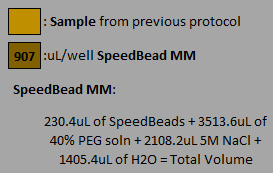

### Img_Key_07b.png

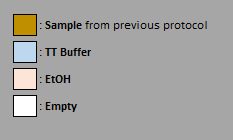

### Img_Key_08_4.png

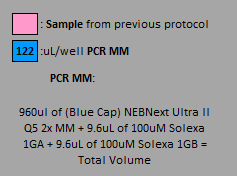

### Img_Key_08_5.png

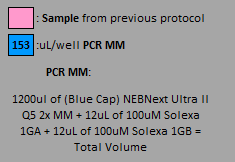

### Img_Key_08_8.png

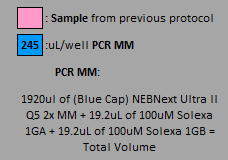

### Img_Key_08_9.png

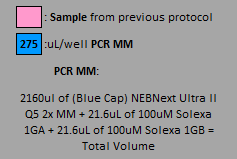

### Img_Key_08_10.png

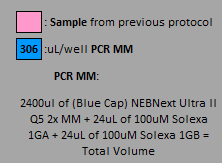

### Img_Key_08_11.png

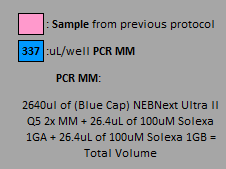

### Img_Key_08_12.png

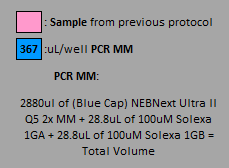

### Img_Key_09a_1.png

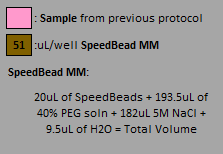

### Img_Key_09a_2.png

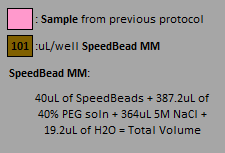

### Img_Key_09a_3.png

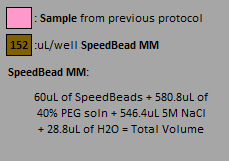

### Img_Key_09a_4.png

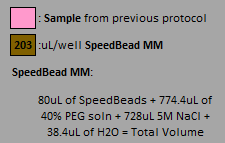

### Img_Key_09a_5.png

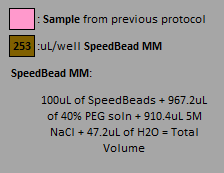

### Img_Key_09a_6.png

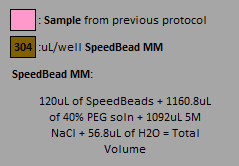

### Img_Key_09a_7.png

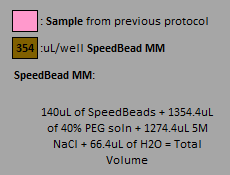

### Img_Key_09a_10.png

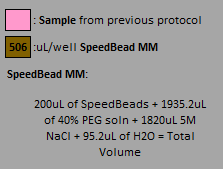

### Img_Key_09a_11.png

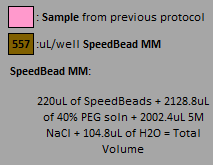

### Img_Key_09a_12.png

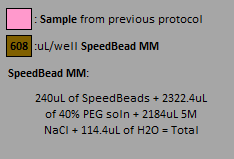

### img_LightBlue.png

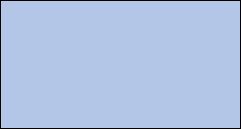

### Img_Mag.png

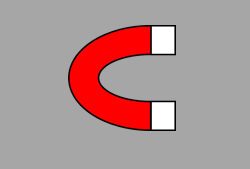

### img_Orange.png

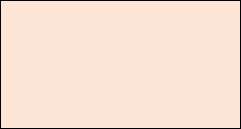

### img_Yellow.png

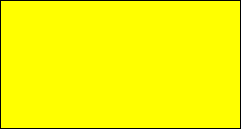

### Tips_no.png

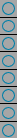

### Tips_yes.png

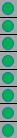
